# Elementary Dynamics of Neural Microcircuits

**DOI:** 10.64898/2026.05.29.728781

**Authors:** Stefano Masserini, Richard Kempter

## Abstract

In neural circuits with multiple excitatory (E) and inhibitory (I) cell types, qualitatively new dynamics arise when one population is silenced by the others, an effect linear theories cannot capture. Using threshold-linear networks, the simplest nonlinear population model, we derive closed-form conditions for multistability, oscillations, and paradoxical responses across microcircuits. In E-I circuits, strengthening E→I rather than I→E connections controls runaway excitation without silencing E cells. In a canonical neocortical circuit, two distinct gamma-oscillation mechanisms reproduce optogenetic silencing experiments, reconciling conflicting models. In hippocampal E-I-I circuits, cross-inhibition determines whether transitions between gamma rhythms are gradual, abrupt, or non-oscillatory. Two E populations or assemblies can cooperate, compete, or suppress each other depending on cross-versus self-excitation. Balanced inhibition prevents competition between E-I assemblies, unless I→I connections are more assembly-specific than I→E ones, a condition automatically met by somatostatin-positive interneurons, which lack mutual inhibition. Lateral inhibition yields winner-take-all dynamics even without assemblies. These testable conditions link connectivity to dynamics across brain areas.

## Introduction

Cell type diversity is a major direction in which systems neuroscience has expanded in the last decade, as the subtype categorization of excitatory (E) and inhibitory (I) neurons has reached a whole new level of detail (Tasic et al., 2018; Campagnola et al., 2022), and specific neuronal populations have often been hypothesized to play key roles in network dynamics (Evangelista et al., 2020; Lagzi and Fairhall, 2024; Veit et al., 2023). These advances have mostly been driven by new experimental techniques (Madisen et al., 2010; Lein et al., 2017), often inspiring circuit-specific modeling. However, many of these dynamics recur across cortical areas, suggesting that a common mathematical description is possible. Steps toward a general description have been taken by employing linear approximations to understand how connectivity shapes responses to perturbations from within or outside the network (Palmigiano et al., 2020; Miller and Palmigiano, 2020; Waitzmann et al., 2024).

Here, we go beyond linear approximations by studying microcircuits in the simplest nonlinear model, the threshold-linear network (TLN). It has been shown that introducing a simple threshold nonlinearity greatly extends the dynamical repertoire of purely linear networks, by allowing for multistability (Curto and Morrison, 2016; Curto et al., 2019, 2020) and oscillations (Tsodyks et al., 1997; Morrison et al., 2024). On the other hand, TLNs retain the fundamental simplicity of linear models, as conditions for each nonlinear regime can be computed in closed form and intuitively interpreted in terms of input and connectivity requirements. These findings, however, were so far mainly applied to all-inhibitory networks (Curto et al., 2024; Morrison et al., 2024) and not used in the context of population dynamics. Here, we extend these insights to excitatory-inhibitory microcircuits, to map previously unrelated systems neuroscience hypotheses to a common reference space and to gain new insights into the dynamics and function of various brain circuits.

Starting with a basic excitatory-inhibitory architecture with two populations (E-I), which has been characterized in the TLN model by Tsodyks et al. (1997), we address some less understood questions. It is intuitive that strong recurrent excitation can be balanced by a strong I→E connection, an approach which has also been taken in models of spiking networks, e.g. by Brunel (2000); but a strong E→I connection can also achieve the same purpose, by more strongly driving the inhibitory neurons. Are these two balancing strategies equivalent? What is the function of I→I connections, which are strongly expressed only by some interneuron types (Pfeffer et al., 2013)?

Next, we examine the most “canonical” expansion of excitatory-inhibitory networks, comprising parvalbumin-positive (PV), somatostatin-positive (SOM), and vasoactive intestinal peptide (VIP) interneurons. The microcircuit architecture involving these interneuron types appears to be stereotypical in different areas (Pfeffer et al., 2013; Letzkus et al., 2015; Canto-Bustos et al., 2022; Campagnola et al., 2022) and has received much attention from the scientific community, especially in its linear approximation (Litwin-Kumar et al., 2016; Waitzmann et al., 2024; Palmigiano et al., 2020). The active state of this circuit, however, is often characterized by highly coherent oscillations in the gamma band, for example in the visual cortex (Eckhorn et al., 1988), which are more poorly understood. Here, we explain the experimental findings by Veit et al. (2017), who have shown that optogenetic inactivation of SOM interneurons stops the oscillation, while inactivation of PV interneurons enhances it.

Oscillatory networks with two distinct inhibitory populations (E-I-I) have also been studied in the hippocampus, but in this case it was hypothesized that each inhibitory population, together with the excitatory population, produces a different gamma oscillation (Keeley et al., 2017). What are the requirements for this bistability of oscillatory states, and how does it relate to classical models of “perceptual” bistability in I-I circuits (Laing and Chow, 2002; Moreno-Bote et al., 2007; Shpiro et al., 2009) and to more recent hypotheses on active-inactive bistability in E-I-I networks (Evangelista et al., 2020)?

Cell-type diversity is not limited to I neurons, as E neurons can often also be classified into distinct, relatively homogeneous groups based on morphological features (Hunt et al., 2018), layer of location (De Kock et al., 2021), function (Kim et al., 2018), or belonging to distinct assemblies. In E-E-I networks, a central question revolves around the interactions between the excitatory populations: if they share the same pool of interneurons, when does each excitatory population excite (Bienenstock, 1995; Hunt et al., 2018) or inhibit (Sammons et al., 2025) the other one, and under which conditions can each population suppress the other, enabling winner-take-all bistability?

Interneurons, like principal cells, also often cluster in assemblies. Some interneurons, namely most neocortical PV interneurons, are strongly connected in both directions to E assemblies receiving correlated inputs (Znamenskiy et al., 2024), a connectivity scheme known as “balanced” inhibition, because it results in a fine-scale E-I balance between the currents received by each neuron (Wehr and Zador, 2003; Okun and Lampl, 2008). Balanced inhibition is useful to stabilize neural activity (Vogels et al., 2011), but can assemblies still compete with each other, through interneurons wired in this way? Other interneurons exhibit a different, “lateral”, connectivity scheme, with E neurons mostly exciting co-tuned I neurons, but I neurons mostly inhibiting competing E neurons, as observed for interneurons in the posterior parietal cortex (Kuan et al., 2024), or hypothesized for SOM interneurons in the orbitofrontal cortex (Lagzi et al., 2021). As lateral inhibition appears ideally suited to enable winner-take-all dynamics, can it even produce them on its own, even if E neurons are uniformly wired without any assembly structure?

By investigating these microcircuits, we both advance the mechanistic understanding of their function and dynamics and demonstrate the relevance of TLNs as a tool for explaining network states as a function of quantifiable connectivity parameters.

## Results

To outline the elementary dynamics of neural microcircuits, we first briefly define the mathematical framework of TLNs and then explain the behaviour of several circuit architectures.

### Mathematical Framework

We consider a general threshold-linear system with a vector of population firing rates *x* = (*x*_*p*_, *x*_*q*_, …*x*_*a*_, *x*_*b*_, …), where we indicate excitatory populations with the letters *p, q*… and inhibitory ones with the letters *a, b*, etc. The general equation is

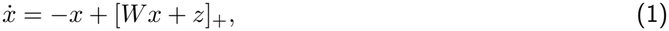

where *z* = (*z*_*p*_, …, *z*_*a*_, …) is the vector of the external inputs, *W* is the coupling matrix, with entries *w*_*ij*_ (*i* post-synaptic, *j* pre-synaptic), and [*x*]_+_ = max*{x*, 0*}* is the threshold-linear activation function. When referring to negative inhibitory weights, the notation |*w*_*ij*_| indicates the connection strength (without sign). We note that, even if these parameters are unitless, all of them can be estimated from spiking models or biological networks. An analysis of the dynamics of TLNs can be found in the Methods and is partially based on previous work on linear (Miller and Palmigiano, 2020) and threshold-linear (Curto et al., 2019) networks. In the Supplementary Materials (SM), we apply the general theory to specific networks of interest to obtain insights that we illustrate in the Results with figures and examples. All the parameters used in simulations are reported in Tables 1–5 of the SM.

**Table 1.** TLN parameters, model *a*–*b*.

|  | *w*<sub>*aa*</sub> | *w*<sub>*ab*</sub> | *w*<sub>*ba*</sub> | *w*<sub>*bb*</sub> | *z*<sub>*a*</sub> | *z*<sub>*b*</sub> |
|---|---|---|---|---|---|---|
| Figure 1C | −1.5 | −[0, 5] | *w*<sub>*ab*</sub> | −1.5 | [0, 200] | 100 |

### I-I Network

Networks with two inhibitory populations (*a* and *b*, Figure 1A) are well understood (Laing and Chow, 2002; Moreno-Bote et al., 2007; Shpiro et al., 2009; Curto and Morrison, 2016), so we use them as a starting point to illustrate the basic dynamics of TLNs. An I-I TLN can have two fundamentally different configurations, set apart by the sign of the expression (|*w*_*aa*_| + 1)(|*w*_*bb*_| + 1) *−* |*w*_*ab*_||*w*_*ba*_|, which is the determinant of the system’s Jacobian in the co-active state. This determinant represents the relative strength of the cross-connections (|*w*_*ab*_| and |*w*_*ba*_|) with respect to the self-connections and leaks (|*w*_*aa*_| +1 and |*w*_*bb*_| +1). If the determinant is positive (weak cross-connections), the two populations can be active together, in a stable fixed point that we indicate as *{a, b}* (SM, Equation (14)). In this case, changing the external input to one population results in smooth transitions to fixed points in which only a single population is active — either *{a}* or *{b}* (Figure 1Bi). These transitions happen when one population gets fully suppressed by the other one. For example, the suppression condition for *b* (i.e. *b* = 0 and *a >* 0) is *z*_*a*_|*w*_*ba*_| *> z*_*b*_(|*w*_*aa*_| + 1) (SM, Equation (12)), that is, the ratio between the inputs *z*_*a*_ and *z*_*b*_ is larger than the ratio between |*w*_*aa*_| + 1 (inhibition from *a* to itself, plus leak) and |*w*_*ba*_| (inhibition from *a* to *b*).

**Figure 1:**
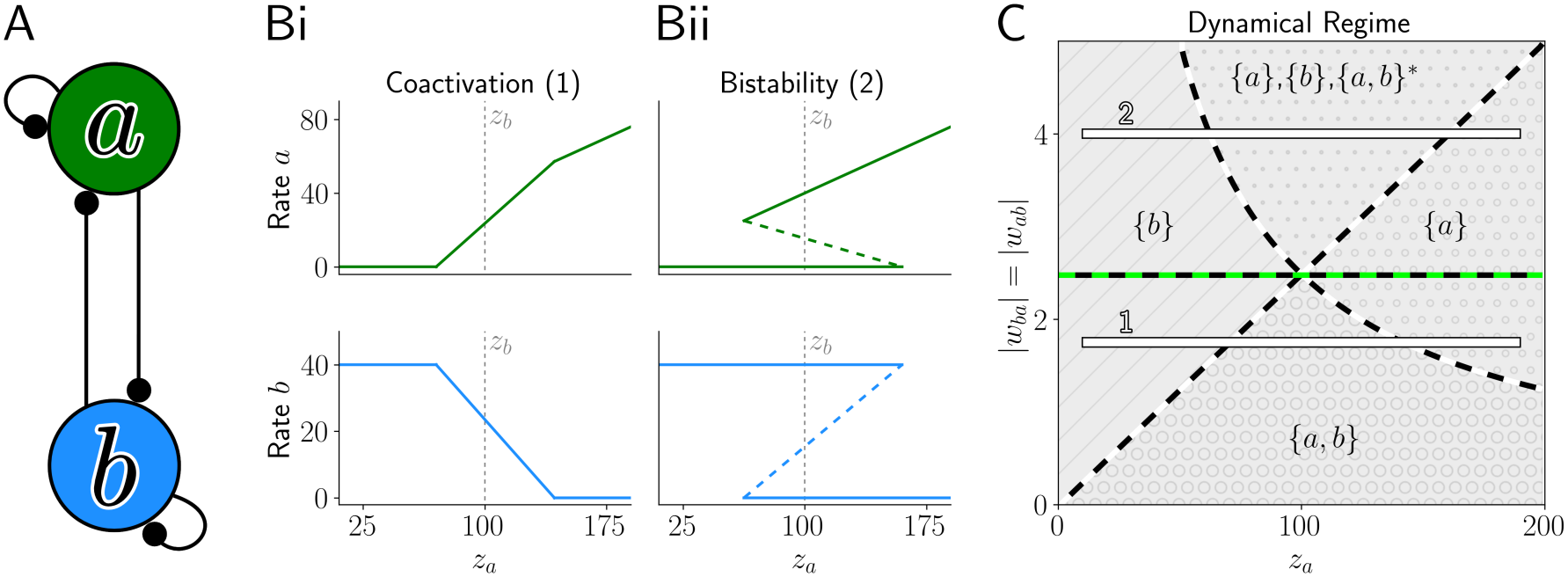
I-I network. (A) Microcircuit sketch with two inhibitory populations *a* and *b*. (B) Bi-furcation diagrams showing the firing rate of each population at stable (continuous lines) and unstable (dashed lines) fixed points when changing the input *z*_*a*_, for positive (Bi) or negative (Bii) determinant of the Jacobian. The value of *z*_*b*_ is marked in gray for comparison. (C) Phase diagram summarizing the different fixed point constellations (* indicates unstable points, same in the following figures). Dashed lines mark transitions between dynamical regimes (same in the following figures). Bars 1 and 2 indicate the cross-sections reported in B. Parameters in Table 1 of the SM.

On the contrary, if the determinant is negative (strong cross-connections), the network can present a bistability between the states *{a}* and *{b}*, separated by an unstable state *{a, b}*^*\**^. In this case, changing the inputs can lead to the sudden disappearance of one stable state through a saddle-node (SN) bifurcation (Figure 1Bii). These bifurcations occur when one population cannot fully suppress the other one anymore. Mathematically, this is the same suppression condition introduced in the case with positive determinant: for example, the stable state *{a}* and the unstable state *{a, b}*^*\**^ disappear together (leaving only *{b}*) as soon as *z*_*a*_|*w*_*ba*_| *< z*_*b*_(|*w*_*aa*_| + 1).

The two different configurations are summarized in Figure 1C: the horizontal green-black line represents where the determinant changes sign, as a function of the cross-connections. Below this line, the suppression conditions (dashed white-black lines), which depend on both the external input *z*_*a*_ and the cross-connections, provide smooth boundaries to the coactive region *{a, b}*. Above the determinant line, instead, the suppression conditions become saddle-node bifurcations and enclose the bistable region.

This relatively simple network shows that TLNs are an ideal tool to understand microcircuits with potentially complicated multistable landscapes, since they allow for explicit closed-form conditions for each possible fixed point of the system and for their bifurcations (Methods, Equation (6)), informing about the role of each parameter.

### E-I Network

Another classical architecture, comprising one excitatory (*p*) and one inhibitory (*a*) population (Figure 2A) has been analyzed in the TLN model (with saturation) by Tsodyks et al. (1997). Here, on the one hand, we briefly reiterate some key findings of that analysis that have broader implications for more elaborate microcircuits, while, on the other hand, we gain novel insights on the role of connections that stabilize neural activity.

**Figure 2:**
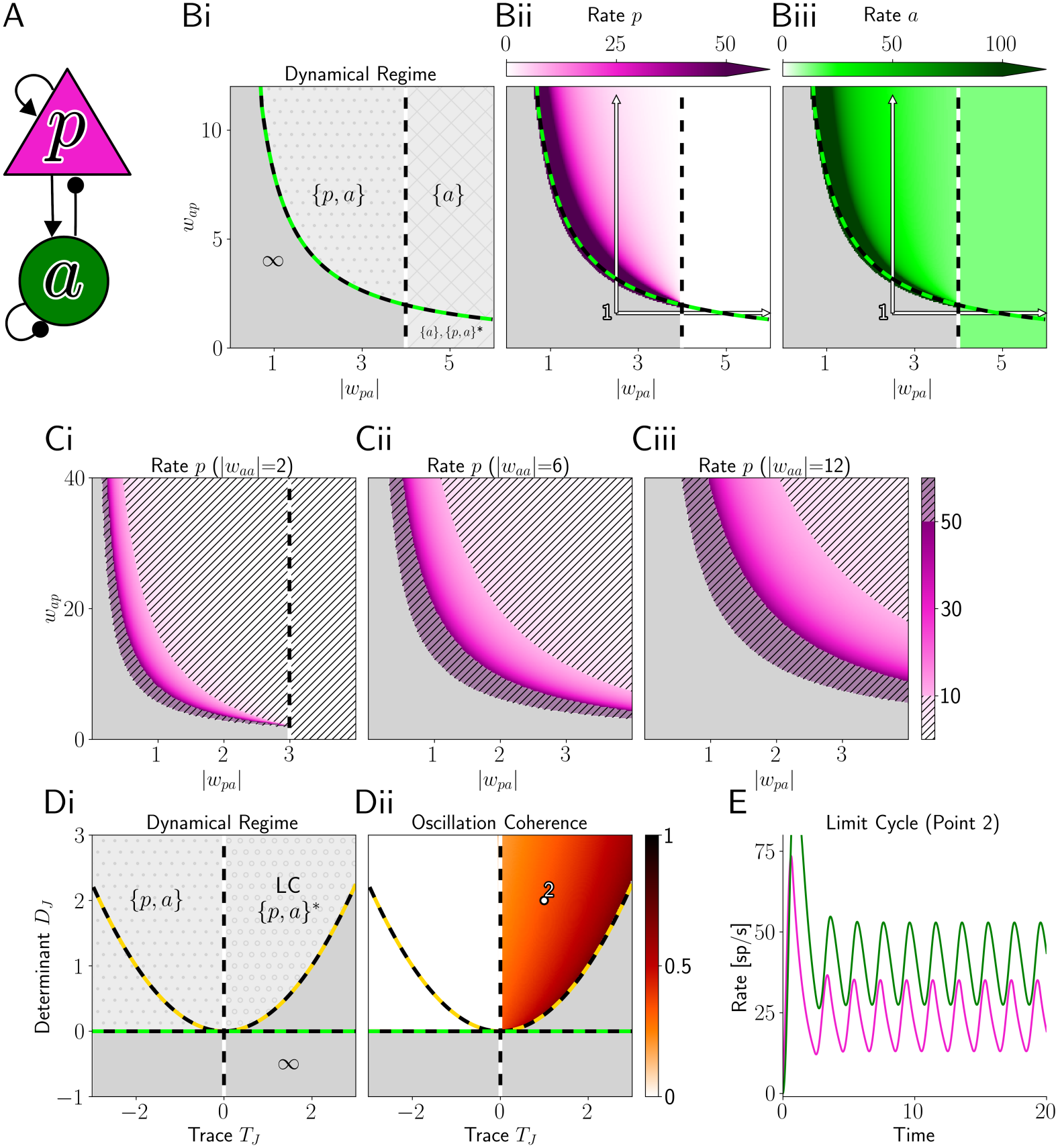
E-I Network. (A) Microcircuit sketch, where triangular shapes indicate excitatory populations/connections and circular shapes inhibitory ones (same in the following figures). (B) Phase diagram (i) (*∞* and gray color represent divergence) and steady state firing rates (ii–iii) as a function of |*w*_*pa*_| and *w*_*ap*_, with arrows indicating different balancing strategies. (C) Steady state firing rates of *p* for different values of the recurrent inhibition *w*_*aa*_. Rates above 50 or below 10 spikes/s are crossed out. (D) Phase diagram (i) and power spectrum coherence (a metric defined in Equation (10)) (ii) as a function of the trace *T*_*J*_ := *w*_*pp*_ *−* |*w*_*aa*_| *−* 2 and the determinant *D*_*J*_ := *w*_*ap*_ |*w*_*pa*_|*−* (*w*_*pp*_ *−*1)( |*w*_*aa*_| + 1). LC marks the limit cycle region. The green dashed line corresponds to zero determinant, the golden one to zero discriminant, and the white one to zero trace (Hopf bifurcation). (E) Network activity for dot ‘2’ in D. Parameters in Table 2 of the SM.

### Properties of the Inhibition-Stabilized Regime

In an E-I TLN, if *w*_*pp*_ *>* 1, the excitatory population *p* can self-sustain its own growth. Therefore, a stable fixed point {*p, a*} can only exist if the inhibitory feedback, i.e. the coupling between *p* and *a*, is strong enough. Like in the I-I network, this requires the determinant *D*_*J*_ = *w*_*ap*_|*w*_*pa*_| *−* (*w*_*pp*_ *−* 1)(1 + |*w*_*aa*_|) to be positive (see SM, section “E-I Network”), which results in an “inhibition-stabilized regime” (Tsodyks et al., 1997) (see also Figure 2B, where the green dashed line indicates *D*_*J*_ = 0). Tsodyks et al. (1997) have shown that, in the inhibition-stabilized regime, external inputs to *a decrease* its own firing in the steady state, a “paradoxical” effect that has found broad experimental confirmation (Sanzeni et al., 2020). The reason for this is that the decrease in the recurrent drive that *a* receives from *p* is larger than the increase in the external current. This paradoxical effect, like the sign and size of any perturbation response, can be determined analytically in TLNs for any fixed point (Methods, section “Perturbations”).

**Table 2.** TLN parameters, model *p*–*a*.

|  | *w*<sub>*pp*</sub> | *w*<sub>*pa*</sub> | *w*<sub>*ap*</sub> | *w*<sub>*aa*</sub> | *z*<sub>*p*</sub> | *z*<sub>*a*</sub> |
|---|---|---|---|---|---|---|
| Figure 2B | 3 | [0, −6] | [0, 12] | −3 | 50 | 50 |
| Figure 2C | 3 | [0, −4] | [0, 40] | −2, −6, −12 | 50 | 50 |
| Figure 2D | [0, 10] | −2 | 20 | −[0, 10] | 100 | 50 |

### Role of Connections in Inhibitory Stabilization

The weights *w*_*ap*_ and *w*_*pa*_, since they are multiplied together in *D*_*J*_, seem to play similar roles in preventing runaway excitation, as can be seen from the hyperbolic shape of the boundary to divergence in Figure 2B. However, these two connections do not have the same effect, as only *w*_*pa*_ is also involved in the suppression condition for *p* (SM, Equation (22), vertical dashed line in Figure 2B). This suppression condition takes the same form as the one presented for the I-I network: in other words, for given positive values of the external inputs, there is a specific value of |*w*_*pa*_| beyond which *p* becomes fully suppressed. This means that a diverging network (point ‘1’ in Figure 2Bii-iii) gains a stable state with *p* silent once *w*_*pa*_ crosses this threshold (horizontal arrow), while divergent trajectories disappear when further crossing the line where the determinant changes sign (green dashed line). Therefore, changing *w*_*pa*_ alone can result in a trade-off between runaway activity and full suppression. On the contrary, increasing *w*_*ap*_ (Figure 2Bii-iii, vertical arrow) allows flexible control of the activity of the excitatory and inhibitory populations, which can span the whole range of possible firing rates without ever getting fully inactivated.

The capability of *a* to confine the firing rate of *p* to a desired intermediate range depends on the recurrent inhibition *w*_*aa*_. Larger |*w*_*aa*_| broadens the region of the parameter space in which the steady-state firing of *p* is within a certain range (SM, Equation (35)). As an example, Figure 2C highlights the hyperbolic “slice” of *w*_*ap*_ and *w*_*pa*_ combinations keeping the firing rate of *p* between 10 and 50 spikes/s — the slice size increases with |*w*_*aa*_|. Thus, larger recurrent inhibition |*w*_*aa*_| increases the robustness of the network to variations in connectivity, which might explain why interneurons that are believed to primarily stabilize the excitatory activity, such as PV interneurons (Veit et al., 2017; Palmigiano et al., 2020), form strong recurrent connections with each other (Pfeffer et al., 2013).

### Gamma Oscillations

When the excitatory-inhibitory loop (*w*_*ap*_|*w*_*pa*_|) is strong enough, the E-I TLN can generate oscillatory solutions, as already highlighted by Tsodyks et al. (1997). Stronger recurrent inhibition |*w*_*aa*_| hampers these oscillations, whereas stronger recurrent excitation *w*_*pp*_ favors them. From a mathematical perspective, this result can be understood as a function of the determinant *D*_*J*_ and of the trace *T*_*J*_ := *w*_*pp*_ *−* |*w*_*aa*_| *−* 2. Oscillatory solutions arise for 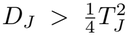 (Figure 2D, region above gold dashed line). For *T*_*J*_ *<* 0, oscillations decay, while for *T*_*J*_ *>* 0 they expand — but unlike in fully linear systems they converge to a limit cycle. The resulting oscillation (Figure 2E) is conceptually a pyramidal-interneuronal network gamma oscillation (PING, Whittington et al. (2000)), since it emerges from the coupling between the E and I populations. The fact that strong recurrent inhibition can prevent stable PING oscillations from emerging has implications for microcircuits with multiple interneuron classes, as we will show in the next section.

### Canonical Circuit

Studies across the neocortex have shown that PV, SOM, and VIP interneurons often present a stereotypical connectivity. PV and SOM cells are both reciprocally connected to excitatory (E) cells, but PV interneurons also make synapses among themselves, unlike SOM interneurons. Finally, SOM also projects to PV, but not the other way around (Pfeffer et al., 2013; Letzkus et al., 2015; Canto-Bustos et al., 2022; Campagnola et al., 2022) (Figure 3A). VIP interneurons mainly project to SOM (Karnani et al., 2016), so here we consider them only as part of the background input to these cells.

**Figure 3:**
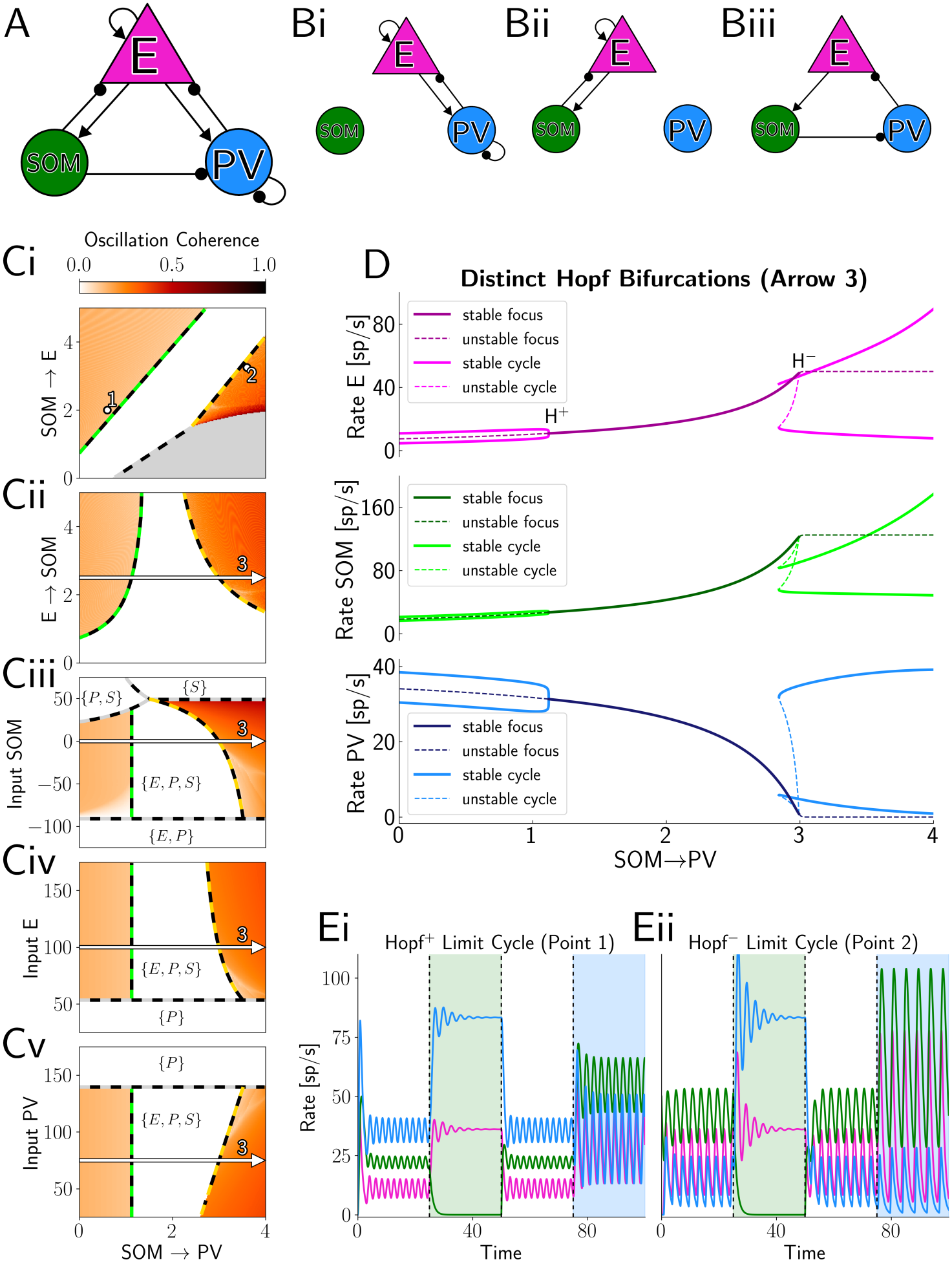
Canonical circuit. (A) Microcircuit sketch. (B) Schematic of the three loops appearing in the Hopf bifurcation (SM, Equation (40)). (C) Coherence of the power spectrum, highlighting oscillatory and non-oscillatory regions, as a function of relevant parameters. Regions with diverging dynamics appear gray. Stable fixed points are highlighted as combinations of E, P (PV) and S (SOM). Dashed lines mark bifurcations and other regime transitions. (D) Bifurcation diagram reporting fixed points and limit cycles when moving along arrow 3 in Cii–v, highlighting the supercritical (H^+^) and subcritical (H^*−*^) Hopf bifurcations. (E) Network activity for dots 1 and 2 in Ci. Colored regions represent optogenetic inhibition of the SOM (green) and PV (blue) populations. Parameters in Table 3 of the SM.

In the visual cortex (Fries, 2009), as well as in other sensory areas (Buzsáki and Draguhn, 2004), sensory stimulation causes gamma oscillations that involve all these cell types. In V1, it has been shown that optogenetic inactivation of SOM interneurons stops the oscillatory activity, while inactivation of PV interneurons enhances it (Veit et al., 2017) — a finding that so far has not been mechanistically explained. In our model, we can analyze the conditions for the canonical circuit to produce stable oscillatory activity, under the assumption that this stems from a PING-like mechanism, i.e. arising from the interactions between E and I cells. Interneuronal network gamma (ING) oscillations relying on recurrent inhibition (Van Vreeswijk et al., 1994), on the contrary, require a delay in the connection from I cells to themselves, which is not included in our model, and thus we exclude them from the analysis.

**Table 3.** TLN parameters, model *p*–*a*–*b*. For the canonical circuit figures, we employ the notation from the SM: *p* for E, *a* for SOM, and *b* for PV.

|  | *w*<sub>*pp*</sub> | *w*<sub>*pa*</sub> | *w*<sub>*pb*</sub> | *w*<sub>*ap*</sub> | *w*<sub>*aa*</sub> | *w*<sub>*ab*</sub> |
|---|---|---|---|---|---|---|
| Figure 3Ci | 4 | [0, −5] | −2.5 | 2 | 0 | 0 |
| Figure 3Cii | 4 | −2 | −2.5 | [0, 5] | 0 | 0 |
| Figure 3Ciii | 4 | −2 | −2.5 | 2.5 | 0 | 0 |
| Figure 3Civ | 4 | −2 | −2.5 | 2.5 | 0 | 0 |
| Figure 3Cv | 4 | −2 | −2.5 | 2.5 | 0 | 0 |
| Figure 4C–F | 4 | −2 | −2 | 10 | −1 | [0, −4] |
| Figure S2A–C | 6 | −2.5 | −2.5 | 11 | −1 | [0, −5] |
|  | *w*<sub>*bp*</sub> | *w*<sub>*ba*</sub> | *w*<sub>*bb*</sub> | *z*<sub>*p*</sub> | *z*<sub>*a*</sub> | *z*<sub>*b*</sub> |
| Figure 3Ci | 6 | [0, −4] | −2.5 | 100 | 0 | 75 |
| Figure 3Cii | 6 | [0, −4] | −2.5 | 100 | 0 | 75 |
| Figure 3Ciii | 6 | [0, −4] | −2.5 | 100 | [−125, 75] | 75 |
| Figure 3Civ | 6 | [0, −4] | −2.5 | [25, 175] | 0 | 75 |
| Figure 3Cv | 6 | [0, −4] | −2.5 | 100 | 0 | [25, 175] |
| Figure 4C–F | 10 | *w*<sub>*ab*</sub> | −1 | 100 | [−50, 50] | −*z*<sub>*a*</sub> |
| Figure S2A–C | 11 | [0, −5] | −1 | 100 | [−100, 100] | −*z*<sub>*a*</sub> |

As a first approach to understanding the experiments by Veit et al. (2017), let us first consider the two subsystems obtained by removing either inhibitory population from the circuit in Figure 3A. From the analysis of E-I networks in the previous section (subsection “Gamma Oscillations”), we know that PING oscillations emerge when recurrent inhibition is comparably weak. The experiments would be consistent with a scenario in which the E-PV subsystem, with strong recurrent inhibition (Figure 3Bi), has a stable fixed point, and the E-SOM subsystem, with no recurrent inhibition (Figure 3Bii), has a limit cycle. However, considering subsystems in isolation does not explain how oscillations arise in the fully connected network, in which the E population is shared by both interneuron classes, and the additional SOM→PV connection is present.

### A first type of oscillation emerges when the SOM→PV connection is weak

In order to understand the dynamical regime of the full network, we examine the eigenvalues of the fixed point {E,PV,SOM}, in which all the populations are active. In this case, we can analytically determine the condition for a Hopf bifurcation in which this fixed point loses stability and a limit cycle emerges (SM, Equation (40)). This condition can be decomposed into three terms, each corresponding to a different loop in the circuit (summarized by Figure 3Bi–iii). Consistently with their role in the isolated subsystems, the E→ SOM→E loop favors the oscillation, while the E→ PV→E loop contributes a negative (desynchronizing) term, when the PV→PV recurrent inhibition is high, as we assume it to be. The disinhibitory E→ SOM→ PV→E loop also hinders the oscillation, which can be intuitively understood in the following way: within this loop, an increase in the E activity causes an increase of the SOM activity, which causes a decrease of the PV activity and thus further disinhibition of the E activity. As activity is always modulated in the same direction, this loop cannot be oscillatory. The SOM→PV connection only appears in the latter, desynchronizing, loop (SM, Equation (40)) and must therefore be weak in order for this oscillation to emerge. We can see this effect e.g. in Figures 3Ci–v, where the above-described Hopf bifurcation (green-black lines) can be found on the left of the non-oscillatory regions, i.e. towards weak SOM→PV connection strengths.

### A second type of oscillation emerges when the SOM→PV connection is strong

Interestingly, in Figure 3C, we can also see a different oscillatory region, arising through a separate bi-furcation (gold-black lines), which instead requires the SOM→PV connection to be *strong*. In order to understand this second oscillation, we perform a semi-analytical bifurcation analysis (Figure 3D, corresponding to the cross-section 3 in Figure 3Cii–v) in a smooth approximation of the TLN system with the softplus activation function (SM, Equation (46)). In the bifurcation diagram, we can see that this second bifurcation appears (on the right) when the {E,PV,SOM} fixed point loses the PV interneurons, which become fully suppressed by the SOM ones. Since the E-SOM subsystem is in the oscillatory configuration, the new fixed point {E,SOM} is an unstable focus, surrounded by a limit cycle. However, although the oscillation focus is two-dimensional, this does not imply that PV interneurons do not fire at all: on the contrary, they still activate at high rates along the trajectory of the limit cycle (Figure 3D). A prerequisite for the existence of this limit cycle is that the E-SOM system has a fixed point (the oscillation focus) with finite values. Otherwise, as soon as PV deactivates, activity diverges, as in Figure 3Ci (gray region). The bifurcation diagram also shows that this second Hopf bifurcation is subcritical: it produces an unstable limit cycle that disappears together with the stable one for a slightly smaller value of the SOM→PV connection. This results in a narrow bistable region (as can be seen in Figures S1D, E).

Since the subcritical Hopf bifurcation occurs when the PV population deactivates, we can determine it analytically like we did for the other suppression conditions discussed in the I-I and E-I sections above. In this case, PV gets suppressed by the joint activity of E and SOM (gold-black subcritical Hopf line in Figure 3C) when increasing the E→SOM (Figure 3Cii) or SOM→PV connections (all panels), or when decreasing SOM→E (Figure 3Ci) or E→PV (SM, Equation (44)). External inputs also appear in this condition: increased inputs to SOM (Figure 3Ciii, also Figure S1Cii for a bifurcation diagram) and to E (in most cases, SM, Equation (43), Figure 3Civ) facilitate it, while increased input to PV (Figure 3Cv) hinders it.

### Both oscillation types are compatible with the experimental results

The effects of the external inputs on the second (subcritical) bifurcation are consistent with the experiments by Veit et al. (2017): if a network exhibits the limit cycle for strong SOM→PV, inhibition of SOM can push it back to the non-oscillatory region (Figure 3Ciii), while inhibition of PV can push it deeper into the oscillatory regime (Figure 3Cv). This behaviour is also illustrated in simulations of optogenetic inhibition of SOM or PV (Figure 3Eii, corresponding to point 2 in Ci). However, a network exhibiting the limit cycle at weak SOM→PV behaves in the same way (Figure 3Ei, point 1 in Ci), even though the external inputs are not directly involved in the supercritical bifurcation. This is because the oscillation can also be stopped by turning off E or SOM, which can happen by either increasing the input to PV (turns off both E and SOM, Figure 3Cv, left), or decreasing the input to SOM (leaves an {E,PV} stable fixed point, Figure 3Ciii, left) — note that SOM does not exhibit a paradoxical response unless the E-PV subsystem is unstable (Palmigiano et al., 2020). In the TLN model, the two oscillation types could in principle be distinguished based on whether a non-oscillatory {E,PV,SOM} state exists for an intermediate value of the input to SOM (Figure 3Ciii, vertical boundary on the left vs non-vertical boundary on the right). As soon as the activation function is smooth, however, this rigid distinction disappears. For example, in a network with the softplus activation function (otherwise characterized by very little discrepancy with the TLN, as seen in Figure S1A), the oscillatory region ends where the activity of E or SOM is not fully zero yet (Figures S1Bi–iv, Ci).

In conclusion, two different mechanisms can explain the results by Veit et al. (2017): an inherently three-dimensional mechanism based on the relative strength of oscillatory and non-oscillatory loops, or a fundamentally two-dimensional mechanism relying on the E-SOM loop and on the suppression of PV interneurons in the oscillation focus. This insight clarifies apparently contradictory results from the modeling literature: on the one hand, spiking network simulations by Parker et al. (2025) identify a gamma oscillation emerging at high values of the SOM→PV connection, while on the other hand a mean field model by Tahvili et al. (2026) finds gamma oscillations for *low* values of the SOM→PV connection, albeit a previous paper by the same authors claimed that such an oscillation emerges in a spiking network when this connection is *strong* (Tahvili et al., 2025). Our results reconcile these seemingly contradictory insights and clarify the mechanism supporting each oscillation, while indicating possible ways to identify the oscillation mechanism in V1. Specifically, we show that the two mechanisms can neither be distinguished based on the activity of the PV interneurons, which fire in both oscillations, nor based on experiments activating or deactivating each population. Manipulating the strength of the SOM→PV synapses, on the contrary, could reveal the underlying mechanism, as one kind of limit cycle would be stopped by strengthening it, the other one by weakening it (Figure 3C). Manipulations of the SOM→E connection could also distinguish between the two mechanisms, as this connection favors the three-dimensional mechanism but hampers the two-dimensional one (Figure 3Ci).

### Limit Cycle Bistability in the E-I-I Network

Neural microcircuits can alternate between different oscillatory states, for example at different frequencies, even without changes in the global brain state. A prominent example is the hippocampal CA1 field, which in awake and moving animals can exhibit gamma oscillations at clearly distinct frequencies (Colgin et al., 2009). Keeley et al. (2017) have put forward the hypothesis that these fast and slow gamma oscillations emerge when the network activity is dominated respectively by a fast and a slow class of interneurons, each receiving their own source of external input. To grasp the basic constituents of these seemingly complex dynamics, we first study an E-I-I circuit without any time constant heterogeneity, in which two identical I populations — *a* and *b* — are identically connected to the E population *p* (Figure 4A). Then, we examine how dynamics change when *a* and *b* have different time constants.

**Figure 4:**
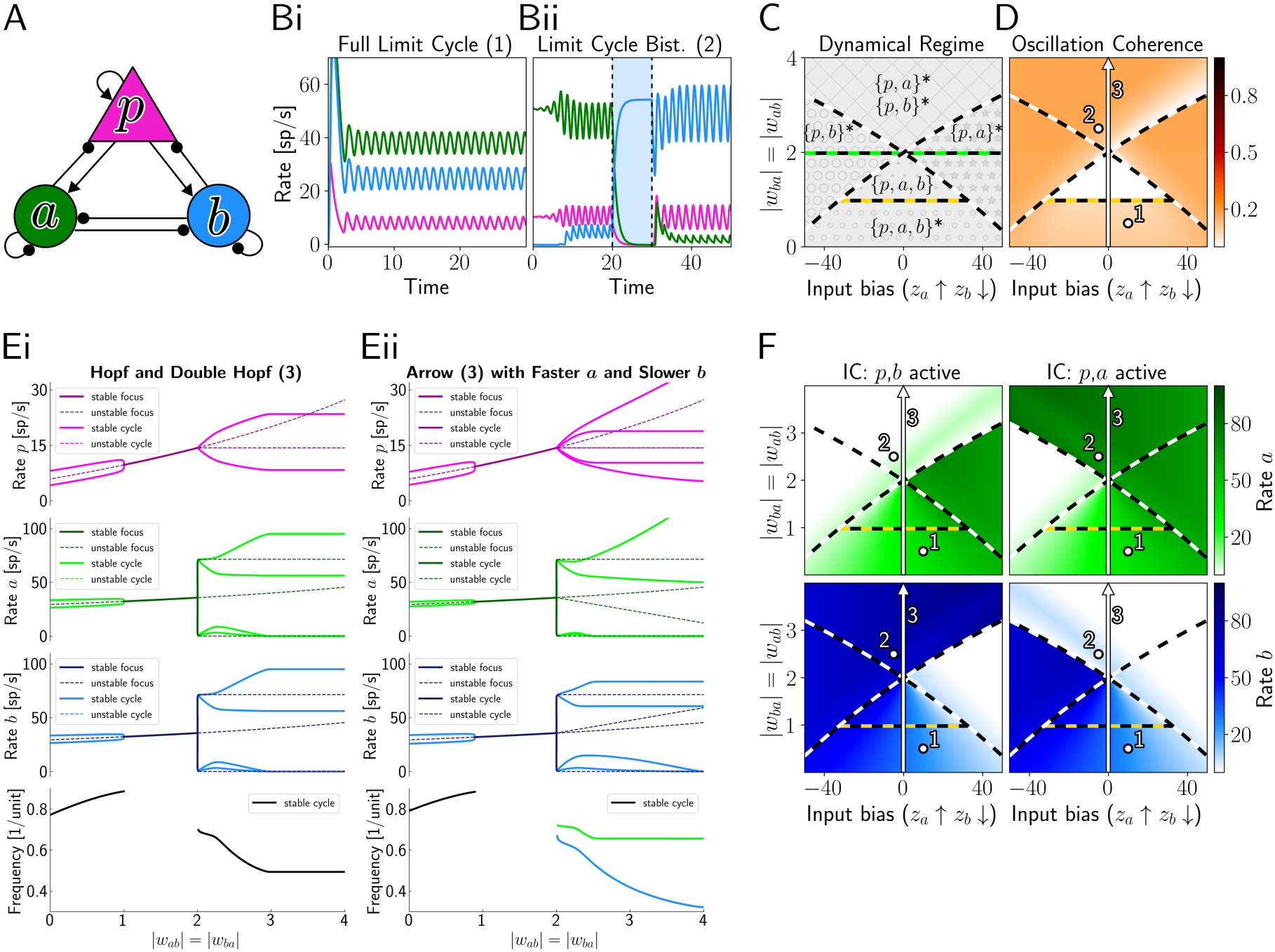
E-I-I network, intermediate recurrent excitation strength. (A) Network sketch. (B) Network activity for example parameters (corresponding to dots 1 and 2 in D,F). The shaded blue region represents a transient stimulation of population *b*. (C) Phase diagram, reporting fixed points and their stability, but not limit cycles. At the black-green line the determinant of {*p, a, b*} changes sign, while at the black-gold line the oscillatory-instability condition for {*p, a, b*} stops holding. Which line is higher determines whether the intermediate non-oscillatory region exists. (D) Power spectrum coherence. (Ei) Bifurcation diagram corresponding to arrow 3. (Eii) Same diagram with time constant heterogeneity. Specifically, time constants are 1 unit for *p*, 0.75 for *a*, 1.5 for *b*. Defining the time constant of population *p* (e.g. 10 ms) allows converting unitless frequencies into actual frequencies. (F) Mean steady state firing rates for different initial conditions (IC). Parameters in Table 3 of the SM.

### Bistability of limit cycles is analogous to simpler kinds of bistability

We find that the symmetric E-I-I circuit can exhibit either a single oscillatory state (Figure 4Bi) or a bistability of limit cycles (Figure 4Bii). This bistability is conceptually analogous to the bistability of fixed points examined in the section on the I-I network and, if *a* and *b* are completely symmetrical, the condition for it is exactly the same (negative determinant of the *a*–*b* subsystem — SM, Equation (49) and following text). As the TLN theory can determine analytically only fixed points (Figure 4C) and the onset of oscillations, we rely on network simulations to verify which states are oscillatory (Figure 4D) and when the network expresses different states if initialized at different initial conditions (Figure 4F). These simulations show that the onset of the bistable region (top region of phase diagrams in Figure 4D,F) through folds of limit cycles (FLC) coincides with the bifurcation lines for the unstable fixed points {*p, a*}^*\**^ and {*p, b*}^*\**^ in Figure 4C around which each limit cycle is centered.

### Interactions between bistability conditions and conditions for oscillations shape the dynamical landscape

The dynamical landscape, however, is complicated by the fact that the cross-inhibitory weights *w*_*ab*_ and *w*_*ba*_ do not only affect the determinant of the *a*–*b* subsystem, but also the stability of the point {*p, a, b*}. As we have seen for the canonical circuit, cross-inhibitory connections favor non-oscillatory dynamics by strengthening desynchronizing loops (SM, Equation (39) for the full expression not specific to the canonical circuit). We first consider the case that recurrent excitation is not too strong, as assumed in Figure 4. In this case, oscillations are produced by this network when the connections *w*_*ab*_ and *w*_*ba*_ are weak (Figure 4C,D, bottom, label {*p, a, b*}^*\**^), oscillations disappear for intermediate values of these connections (Figure 4C,D, middle, label {*p, a, b*}), while further strengthening them gives rise to the two limit cycles with 2-dimensional foci (Figure 4C,D, top, labels {*p, a*}^*\**^ and {*p, b*}^*\**^). A semi-analytical bifurcation analysis validates these insights (Figure 4Ei) and confirms that the appearance of the two limit cycles (on the right) coincides with the pitchfork-like bifurcation of the fixed points. When performing the same analysis in a network with time constant heterogeneity, we find the same bifurcations, but the *a*-dominated and the *b*-dominated cycles now oscillate at different frequencies. In the alternative scenario of strong recurrent excitation (SM, Equation (52)), the limit cycles with 2-dimensional focus can directly emerge from the limit cycle with 3-dimensional focus, through a pitchfork of limit cycles (Figure S2Di). In this case, we note that, away from the pitchfork, the two FLC bifurcations do not exactly coincide with the SN bifurcation of the fixed points (Figure S2C, Dii).

Since CA1 is characterized by a rather weak recurrent excitation (Deuchars and Thomson, 1996), the scenario considered in Figure 4 appears more relevant. In this case, a changing proportion between the inputs *z*_*a*_ and *z*_*b*_ could, depending on the strength of cross-inhibition, induce a transition between the *a*-dominated and the *b*-dominated oscillation in three fundamentally distinct ways: through an oscillation with intermediate features (Figure 4D,F, bottom), through an intermediate non-oscillatory state (Figure 4D,F, middle), or through a sudden switch (Figure 4D,F, top). Provided that fast and slow gamma are indeed associated with different classes of interneurons inhibiting each other, further experiments could manipulate also the strength of cross-inhibitory connections. Together, changing these weights and/or the inputs to inhibitory neuron classes would allow testing the model predictions and determining how state transitions occur.

### E-E-I Network

We now transition to studying networks with one inhibitory (*a*) and two excitatory (*p* and *q*) populations (Figure 5A). This microcircuit motif is very versatile and can describe the clustering of excitatory neurons into phenotypically distinct subpopulations with different functions (Kim et al., 2018; Hunt et al., 2018; De Kock et al., 2021), or also into assemblies processing specific features. In either case, the activation of an excitatory population can result in the subsequent activation of another excitatory population, like in the case of synfire chains (Bienenstock, 1995), or in their inhibition or even suppression (Sammons et al., 2025). Mutual suppression of competing assemblies can result in a multi-stability that, in the presence of noise, can produce the metastable dynamics typical of spontaneous activity in different brain regions (Kenet et al., 2003; O’Hashi et al., 2018; Sasaki et al., 2007). Here, we analyze the requirements for these cooperative or competitive interactions. We first consider the case of a symmetric network, which is best suited for describing two E assemblies with roughly analogous features, and then extend the analysis to asymmetric networks, most relevant when the E populations represent distinct neuron types.

**Figure 5:**
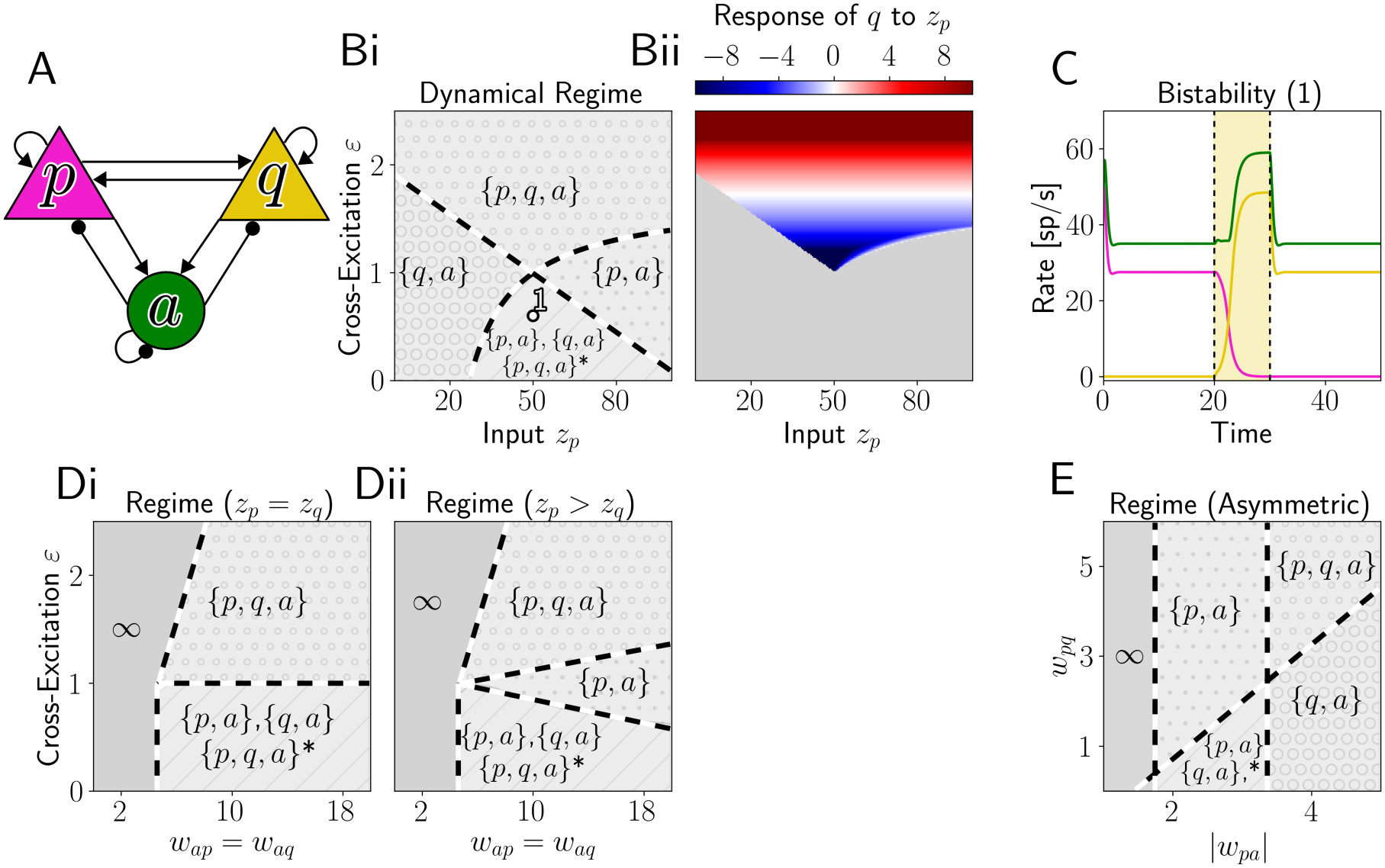
E-E-I network. (A) Microcircuit sketch. (Bi) Phase diagram. (Bii) Response of *q* to an extra input of 10 delivered to *p*. Regions in which the baseline state is not {*p, q, a*} appear gray. (C) Example of bistable network configuration. The yellow shaded region represents a transient stimulation of *q*. (D) Phase diagrams for equal (i) and biased (ii) inputs. (E) Phase diagram for an asymmetric network (* represents the unstable {*p, q, a*}). Parameters in Table 4 of the SM.

### Symmetric Network

We assume that the connections involving *p* and *q* are symmetrical: *w*_*qq*_ = *w*_*pp*_, *w*_*qp*_ = *w*_*pq*_, *w*_*aq*_ = *w*_*ap*_, and *w*_*qa*_ = *w*_*pa*_, and refer to their value as the first element of each pair. In addition, we define the scaling factor 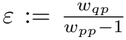, which represents the strength of cross-excitation with respect to the self-excitation. When *w*_*pp*_ *<* 1, *ε* is *<* 0, but we primarily focus on *w*_*pp*_ *>* 1 (always resulting in *ε >* 0), since indications of the inhibition-stabilized regime have been found throughout most sensory areas (Sanzeni et al., 2020; Adesnik, 2017; Kato et al., 2017).

**Table 4.** TLN parameters, model *p*–*q*–*a*.

|  | *w*<sub>*pp*</sub> | *w*<sub>*pq*</sub> | *w*<sub>*pa*</sub> | *w*<sub>*qp*</sub> | *w*<sub>*qq*</sub> | *w*<sub>*qa*</sub> |
|---|---|---|---|---|---|---|
| Figure 5B | 3 | [0, 5] | −3 | *w*<sub>*pq*</sub> | 3 | −3 |
| Figure 5Di | 3 | [0, 5] | −3 | *w*<sub>*pq*</sub> | 3 | −3 |
| Figure 5Dii | 3 | [0, 5] | −3 | *w*<sub>*pq*</sub> | 3 | −3 |
| Figure 5E | 3 | [0, 6] | [−1, −5] | 1.5 | 3 | −3 |
| Figure S6 | 1.6 | 3.2 | −4 | [0, 4.8] | [0, 3.2] | −4 |
|  | *w*<sub>*ap*</sub> | *w*<sub>*aq*</sub> | *w*<sub>*aa*</sub> | *z*<sub>*p*</sub> | *z*<sub>*q*</sub> | *z*<sub>*a*</sub> |
| Figure 5B | 8 | 8 | −6 | [0, 100] | 50 | 25 |
| Figure 5Di | [0, 20] | *w*<sub>*ap*</sub> | −6 | 50 | 50 | 25 |
| Figure 5Dii | [0, 20] | *w*<sub>*ap*</sub> | −6 | 55 | 50 | 25 |
| Figure 5E | 8 | 8 | −6 | 50 | 50 | 25 |
| Figure S6 | 7.2 | 0.8 | −8 | 24 | 32 | 40 |

In our analysis, we consider a “resting state” in which the external inputs *z*_*p*_ and *z*_*q*_ are not biased toward a specific assembly (e.g. both 50 at the center of Figure 5B) and study how this state changes when the inputs become biased toward one E assembly. Based on the value of the cross-excitation *ε*, we can set apart three different scenarios. Below a first threshold, *ε*_1_ = 1, the resting state has two competing stable states: {*p, a*} and {*q, a*} (Figure 5B, bottom center), which both disappear for *ε > ε*_1_ (SM, Equation (62)). This upper bound for the states {*p, a*} and {*q, a*} changes when the two inputs are not perfectly balanced: the one for {*p, a*} increases with *z*_*p*_, while the one for {*q, a*} decreases (SM, Equation (67)). As a result, starting from the bistable configuration, inputs to either population (e.g. *p*) can force the network into the state (e.g. {*p, a*}) in which the other E population (*q*) is suppressed (see also Figure 5C). When the input terminates, the network can still remain in the same configuration, due to hysteresis.

If *ε >* 1, on the contrary, populations *p* and *q* can be both active at the resting state. Below a second threshold 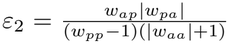 (always larger than *ε*_1_, if E-I balance conditions are satisfied), an increasing input to one E population results in a continuous transition from co-activation to suppression of the other population (SM, Equation (65)). In the *ε*_1_ *< ε < ε*_2_ band, the interaction between the excitatory populations is negative, meaning that a positive perturbation of one of them has a negative effect on the other one (Figure 5Bii, bluish region). This is because the excitatory pathway from one population to the other, *ε*(*w*_*pp*_ *−*1), is weaker than the inhibitory pathway 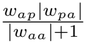 mediated by *a* (SM, Equation (65)). Above *ε*_2_, the *p*–*q* interaction turns positive and increasing *z*_*p*_ does not turn off *q* anymore, but rather increases its rate too (Figure 5Bii, reddish region). Increasing *w*_*ap*_ broadens the mono-stable regions if the drives are different, while it has no effect if the drives are the same (Figure 5D). In conclusion, we have shown that the strength of cross-excitation with respect to self-excitation determines whether cooperation, competition, or mutual suppression occurs in a symmetric E-E-I network, and we have predicted the specific values at which these transitions would occur.

### Asymmetric Network

The analysis on symmetric E-E-I networks can be extended to asymmetric ones, where *p* and *q* are not identical. At the resting state (equal drives *z*_*p*_ = *z*_*q*_), whether one E population suppresses the other depends on two comparisons (SM, Equation (57)). Taking suppression of *p* as an example, the first ingredient is the comparison between the excitation that *q* sends to *p* (*w*_*pq*_) and the excitation that *q* sends to itself (*w*_*qq*_ *−* 1, including the leak): the weaker *w*_*pq*_, the easier it is for *q* to suppress *p*. The second ingredient is the comparison between the inhibition that *a* sends to the two populations (|*w*_*pa*_| vs |*w*_*qa*_|): the more *a* targets *p* rather than *q*, the easier the suppression of *p*. Figure 5E illustrates both dependencies: *p* is suppressed (bottom right) when *w*_*pq*_ is small and |*w*_*pa*_| is large, i.e. below and to the right of the diagonal line. The suppression of *q*, instead, does not depend on *w*_*pq*_, so in this panel it is set by |*w*_*pa*_| alone and marked by a vertical line. Together, the intersection between the two lines gives rise to the same four regimes found in the symmetric network in Figure 5Bi.

If one of the two comparisons is an equality, the other one determines the outcome. If *a* inhibits *p* and *q* equally (|*w*_*pa*_| = |*w*_*qa*_|), *p* is suppressed exactly when *q* excites *p* less than it excites itself (*w*_*pq*_ *< w*_*qq*_ *−* 1). Conversely, if *q* excites *p* as strongly as itself (*w*_*pq*_ = *w*_*qq*_ *−* 1), *p* is suppressed exactly when *a* inhibits *p* more strongly than *q* (|*w*_*pa*_| *>* |*w*_*qa*_|). As an application, we consider the connectivity between “thorny” and “athorny” pyramidal cells in the CA3 region of the hippocampus. It was shown that athorny neurons target each other much more often than they target thorny ones, while thorny neurons target athorny neurons more often than they target other thorny ones (Sammons et al., 2025). Assuming that these populations are equally targeted by the interneurons, this connectivity would imply that there is a single stable point, in which only athorny cells and interneurons are active. This is indeed observed during the first part of sharp-wave events, when athorny cells activate without the thorny ones (Hunt et al., 2018). Similar predictions could be made about any network in which the connectivity between two E subpopulations is known, for example in the visual cortex (Kim et al., 2018) or in the somatosensory cortex (De Kock et al., 2021).

### E-E-I-I Network

In the previous section, we have shown that, if two E assemblies share a single pool of interneurons (“blanket” inhibition), the strength of cross-excitation is the main parameter that determines competitive or cooperative interactions. This is a major drawback, since cognitive processes might require more flexibility. For example, two weakly connected assemblies may need to activate together to form new associations, or two strongly connected assemblies may need not to activate together. Here, we investigate whether the clustering of interneurons into inhibitory assemblies with specific connectivity with the E assemblies allows for more flexible dynamics.

We consider a network with two E assemblies *p* and *q* and two I assemblies *a* and *b*. We regard each of the pairs *p*–*a* and *q*–*b* as an E-I module that is activated by the same stimuli and therefore receives correlated inputs. We simplify the system by assuming that the two E-I modules are identical, with only four independent parameters *w*_*pp*_ = *w*_*qq*_, *w*_*pa*_ = *w*_*qb*_, *w*_*ap*_ = *w*_*bq*_, and *w*_*aa*_ = *w*_*bb*_. We also assume that the two modules exchange symmetric cross-connections. Four independent parameters express the specialization of each pair of cross-connections with respect to the within-module connections (Figure 6A). Specifically, the parameter 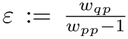, already considered in the E-E-I section, scales the strength of cross-excitation between the E assemblies. As before, we primarily focus on *w*_*pp*_ *>* 1 and therefore on *ε >* 0. Lower specialization of excitatory connections is thus represented by a higher value of *ε*. The same applies to three more parameters. The parameter 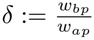 determines how strongly an E assembly targets the “other” I assembly compared to the “own” one, and we would typically have *δ <* 1 in most networks with inhibitory assemblies. The parameter 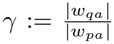, on the contrary, in combination with *δ <* 1, sets apart two cases both well represented in the brain. *γ <* 1 corresponds to “balanced” inhibition, according to which I assemblies inhibit their own E assembly more strongly (Figure 6A), while *γ >* 1 corresponds to “lateral” inhibition, according to which I assemblies inhibit the *other* E assembly more strongly (Figure 7A). Finally, 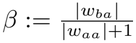 determines the degree of competition between the I assemblies. Note that the value of *β* corresponding to unspecialized I→I connections grows with *w*_*aa*_, because the denominator includes the leak.

**Figure 6:**
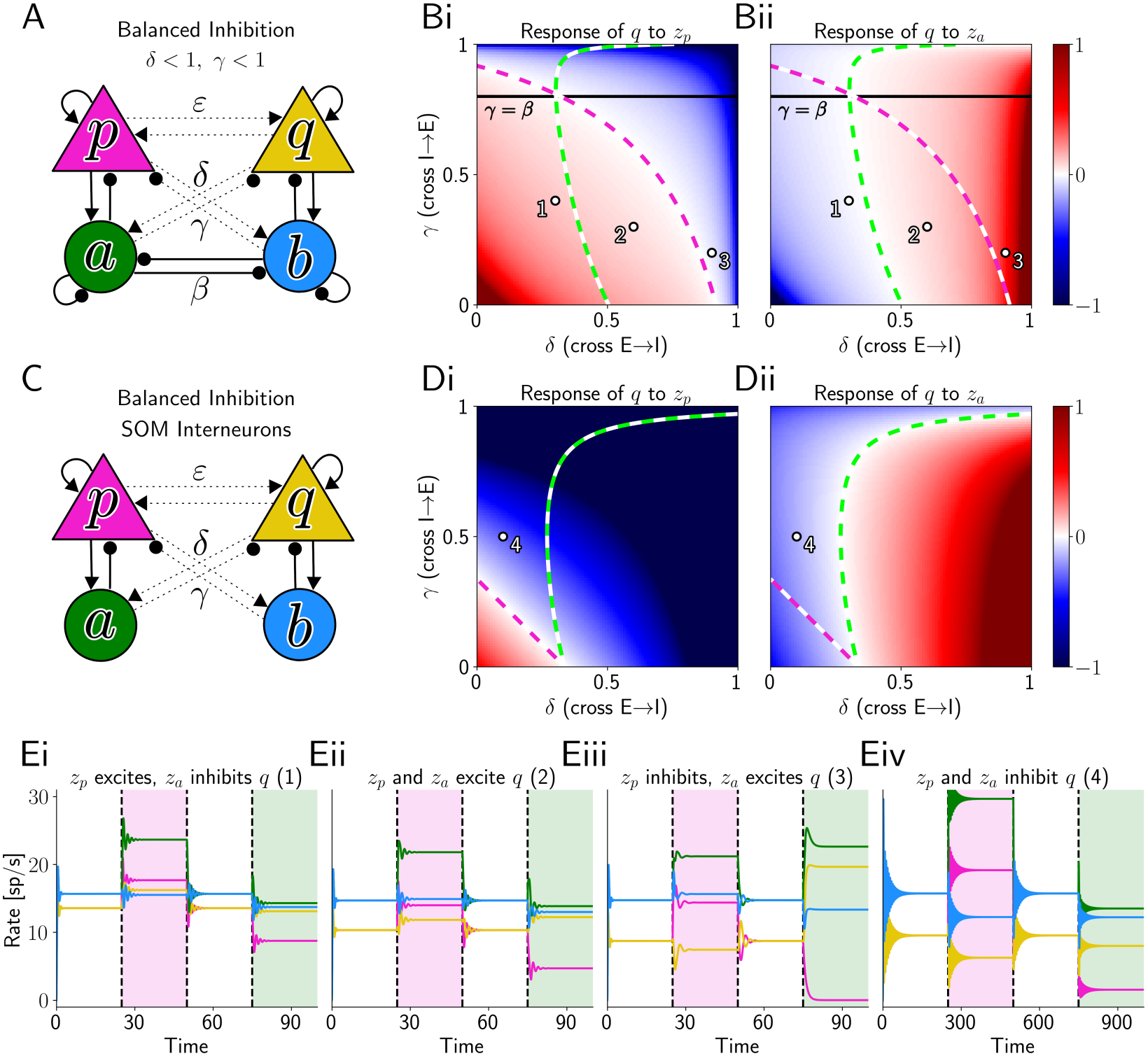
Input responses of the E-E-I-I network with balanced inhibition. (A) Sketch of a network with balanced inhibition, in which dashed arrows represent weaker connections. (B) Response of *q* to an extra input of 1 delivered to either *p* (i) or *a* (ii). Parameter values at which these responses change sign are marked by white-magenta and white-green lines respectively. (C) Sketch of a network without recurrent inhibition. (D) Same as B, for the network in C. (E)Network activity for the dots in B and D. The colored regions represent a transient stimulation of *p* (magenta) or *a* (green). Parameters in Table 5 of the SM.

**Table 5.**
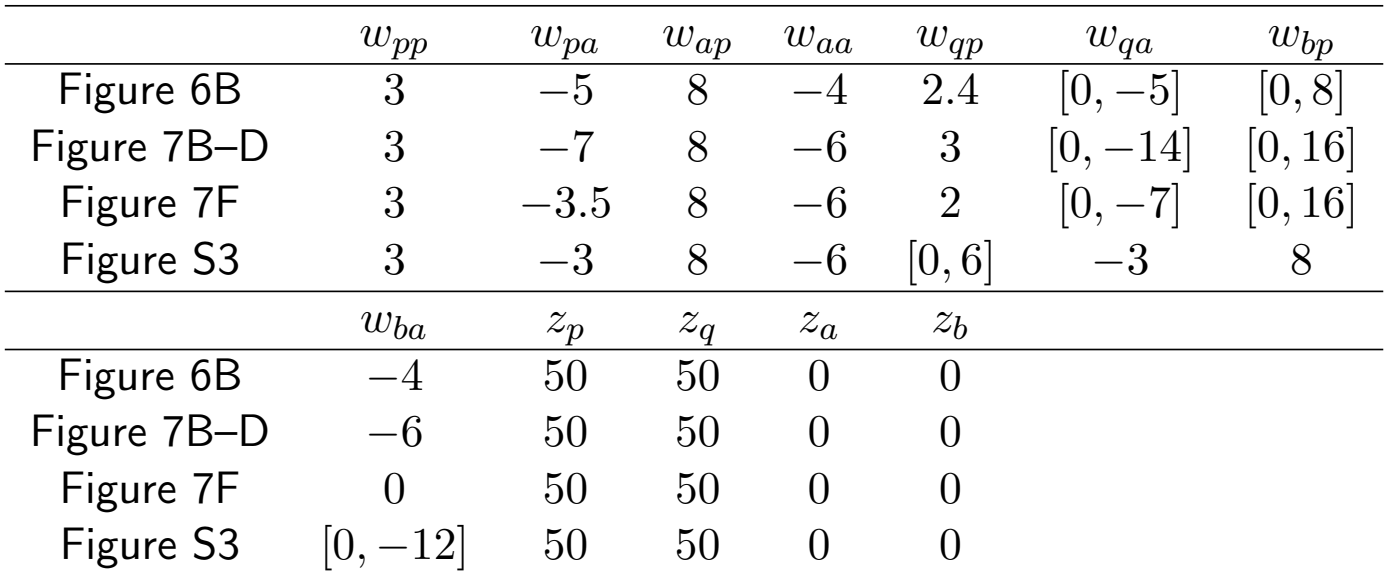
TLN parameters, model *p*–*q*–*a*–*b*. Each connection parameter is identical to its symmetric counterpart.

|  | w<sub>pp</sub> | w<sub>pa</sub> | w<sub>ap</sub> | w<sub>aa</sub> | w<sub>qp</sub> | w<sub>qa</sub> | w<sub>bp</sub> |
|---|---|---|---|---|---|---|---|
| Figure 6B | 3 | −5 | 8 | −4 | 2.4 | [0, −5] | [0, 8] |
| Figure 7B–D | 3 | −7 | 8 | −6 | 3 | [0, −14] | [0, 16] |
| Figure 7F | 3 | −3.5 | 8 | −6 | 2 | [0, −7] | [0, 16] |
| Figure S3 | 3 | −3 | 8 | −6 | [0, 6] | −3 | 8 |
|  | w<sub>ba</sub> | z<sub>p</sub> | z<sub>q</sub> | z<sub>a</sub> | z<sub>b</sub> |  |  |
| Figure 6B | −4 | 50 | 50 | 0 | 0 |  |  |
| Figure 7B–D | −6 | 50 | 50 | 0 | 0 |  |  |
| Figure 7F | 0 | 50 | 50 | 0 | 0 |  |  |
| Figure S3 | [0, −12] | 50 | 50 | 0 | 0 |  |  |

**Figure 7:**
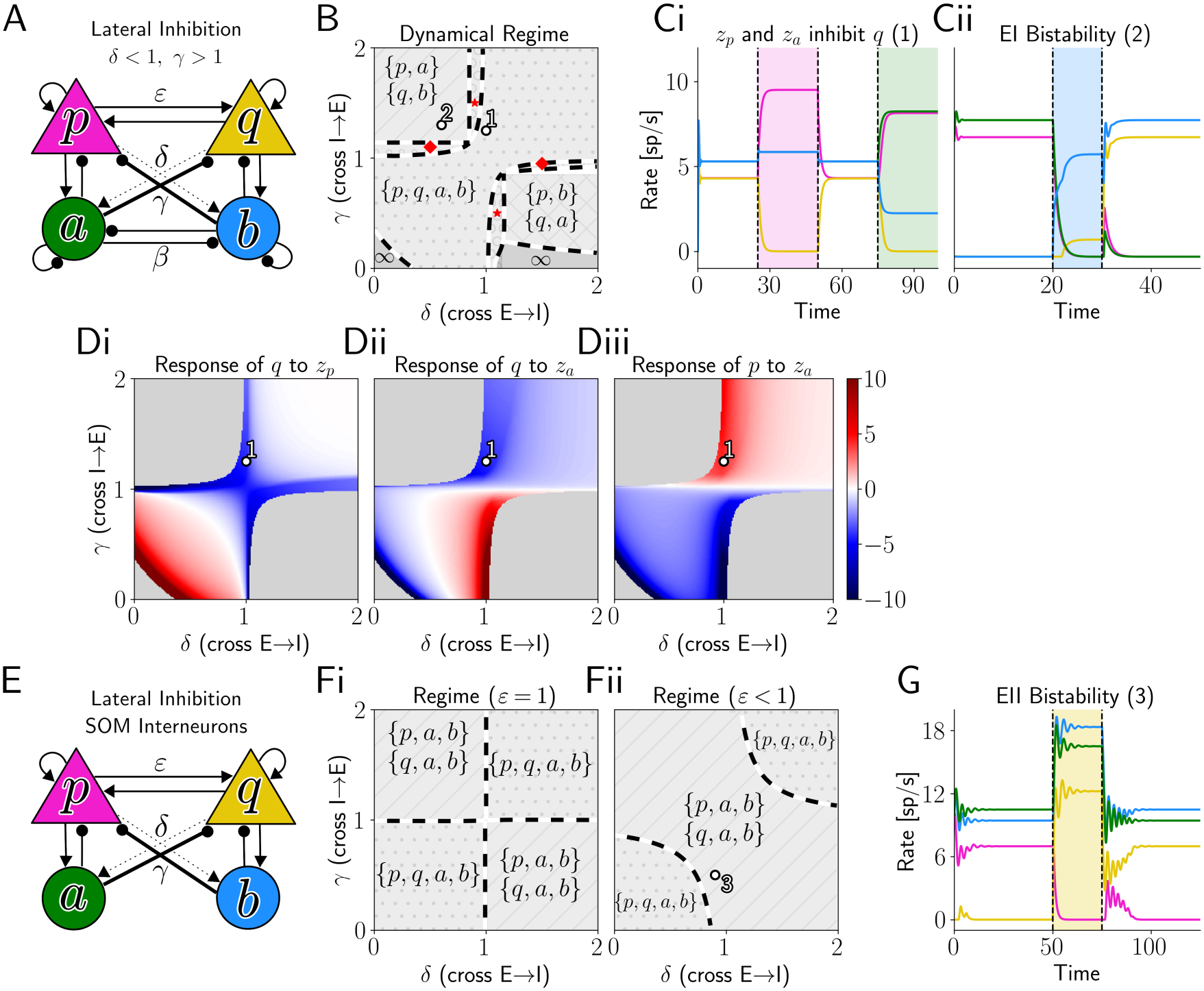
Dynamical regimes of E-E-I-I networks as a function of cross-connections. (A) Sketch of a network with lateral inhibition, in which thick and dashed arrows represent stronger and weaker connections, respectively. (B) Phase diagram (only stable states are displayed). Red stars mark regions of {*p, a, b*} - {*q, a, b*} bistability, red diamonds {*p, q, a*} - {*p, q, b*} bistability. (C) Network activity for dots 1 and 2 in B. The colored regions represent a transient stimulation of the population of matching color. (Di-ii) Response of *q* to an extra input of 10 delivered to either *p* (i) or *a* (ii). (Diii) Same for the response of *p* to input to *a*. (E–G) Same as in A–C, for a network without recurrent inhibition.

The SM reports a full characterization of the fixed-point configurations that can emerge in this network and of the resulting interaction between assemblies. In the next subsections, we explore non-trivial effects of the specialization parameters, focusing on regimes typically encountered in the brain, namely balanced (Figure 6) and lateral (Figure 7) inhibition.

### Balanced Inhibition

We show here that the specialization of the I→E connections (*γ <* 1), typical of networks with balanced inhibition, hampers the competition between E-I modules. We define competition between E-I modules in the following way: (i) input to *p* decreases the firing rate of *q*; (ii) input to *a* decreases the firing rate of *q*. These requirements ensure that any coordinated input to *p* and *a* coherently decreases activity in the competing E population. We prove that competition between E-I modules is only possible if *γ > β*, i.e. if the amount of I→E cross-inhibition is large in comparison to the I→I cross-inhibition (SM, Equation (79) and following). Somewhat surprisingly, this lower bound for *γ* is unaffected by the cross-excitation *ε* and by the specialization *δ* of the E→I connections.

Figure 6B illustrates the different ways in which E-I modules can fail to develop competitive interactions when *γ < β*. On the one hand, when *δ* is small, input to *a* inhibits the competing E population *q* (Figure 6Bii), but input to *p* excites it (Figure 6Bi, example (1); see also Figure 6Ei). On the other hand, when *δ* is large (but still *<* 1), input to *p* inhibits *q*, but input to *a* disinhibits it (example (3); see also Figure 6Eiii). These two scenarios are separated by an intermediate region in which *q* is excited by input to either *p* or *a* (example (2); see also Figure 6Eii). The boundaries at which these two perturbation responses change sign intersect on the line *γ* = *β* (Figure 6B, here *β* = 0.8). Therefore, only when *γ > β* is there a regime in which any coordinated input to the E-I module *p*–*a* is guaranteed to decrease the firing rate of *q*.

For interneurons with strong recurrent inhibition (e.g. PV), *β* represents the specialization of I→I connections. When such connections are weak, however, *β* is small regardless of specialization, and is 0 if such connections are absent, as for SOM interneurons. Therefore, in a network of E-SOM assemblies (Figure 6C), the constraint *γ > β* vanishes and competition is always possible even at low values of *γ* (Figure 6D, example (4); see also Figure 6Eiv). This makes SOM interneurons ideally suited to support competition, whatever their wiring.

To summarize, a network with balanced inhibition that nevertheless needs E-I modules to compete with each other requires weak connectivity between the I assemblies. For interneurons with strong recurrent inhibition, such as PV interneurons, this requirement means a direct comparison between the specialization of E→I and I→I connections. This prediction highlights the importance of structured I→I connections, so far understudied in the experimental literature, which has mostly focused on E→I and I→E specialization (Znamenskiy et al., 2024; Kuan et al., 2024; Ogando et al., 2025). In contrast, interneurons with weak recurrent inhibition, such as SOM interneurons, fulfill the condition automatically and can always support competition.

### Lateral Inhibition

Altogether, the results presented so far demonstrate the limitations of balanced inhibition with respect to competition between assemblies. Therefore, it is not surprising that networks for which the suppression of competing assemblies is a priority, e.g. the decision-making circuits of the posterior parietal cortex, often exhibit a different connectivity scheme (Kuan et al., 2024), which we termed lateral inhibition (*δ <* 1, *γ >* 1, Figure 7A). This asymmetry in the specialization of E→I and I→E connections causes each E-I module to act as a single unit that inhibits the other module, naturally resulting in competition and bistability (SM, Equation (85)).

In addition to supplementing an existing assembly structure, lateral inhibition is by itself sufficient to induce functional E-I modules in an otherwise unstructured network. Specifically, we ask whether a network with completely unspecialized E→E and I→I connections, i.e. without assemblies in either the E or the I population, settles into a co-active state or into competing ones at rest, that is, when both modules receive the same external input. Lateral inhibition alone is sufficient for the latter: through the combined effect of *γ >* 1 and *δ <* 1, the network becomes bistable between the winner-take-all states {*p, a*} and {*q, b*} (Figure 7B, top left region, example in Figure 7Cii). We also observe that both requirements for competition between E-I modules (inputs to *p* and *a* both inhibiting *q*) are satisfied for lateral inhibition whenever the four assemblies are active together (Figure 7Di-ii, top left quadrant, example in Figure 7Ci). In addition, input to *a* even increases the firing rate of its own E assembly *p* (Figure 7Diii).

If connections between the two I assemblies are absent, for example for assemblies of SOM interneurons, which lack recurrent inhibitory connections altogether (Figure 7E), even slightly lateral inhibition is sufficient for resting-state bistability, as long as *ε ≤* 1 (SM, Equation (83)). Figure 7Fi shows this for the limiting case *ε* = 1: the whole lateral inhibition quadrant is bistable between {*p, a, b*} and {*q, a, b*}, with both I assemblies always active. When *ε <* 1, bistability is possible even with balanced inhibition (Figure 7Fii, example in Figure 7G) We do not focus on *δ >* 1, which is less relevant biologically, but we predict that such networks would exhibit the same states with the E-I pairing swapped, in both cases considered in Figure 7. Additional multi-stable configurations can be seen in Figure S3 and are discussed in the SM.

In conclusion, lateral inhibition can by itself produce winner-take-all dynamics, even in networks that lack E- and I-assemblies in the traditional sense. When I→I cross-connections are strong, the losing E-I module is silenced as a whole, whereas when they are weak only the losing E assembly is suppressed. In addition, when such connections are absent, like in an E-SOM network, we predict that even slightly lateral inhibition would produce a resting state in which single E assemblies activate alone, rather than activity being spread equally through the network. Since lateral inhibition is so well suited to competition, we suggest that balanced E-I networks that cannot support competition due to the *γ > β* constraint (see subsection above) could include a different interneuron type that instead specializes in lateral inhibition. From this perspective, the prediction of Lagzi et al. (2021) that, as a consequence of the plasticity measured in the orbitofrontal cortex, PV interneurons would develop balanced, and SOM interneurons lateral clustering, is especially suggestive. However, if cross-excitation between assemblies is weak, SOM interneurons can support bistability even with balanced inhibition

## Discussion

In this study, we have shown that multiple, previously unrelated hypotheses in systems neuroscience can be understood within a unified theoretical framework, which exploits mathematical insights into the dynamics of threshold-linear networks (TLNs). This includes a broad class of population dynamics that rely on undelayed network interactions between homogeneous neuronal populations. Here, we have applied this theory to push forward our understanding of elementary microcircuits.

In a classical E-I circuit, we have found that E→I connections are more effective than I→E connections in controlling the activity of a population with strong recurrent excitation, without suppressing it. Strong I→I connections on the one hand broaden this “useful” range of inhibition, while on the other hand they hinder the emergence of PING oscillations, two effects that are consistent with recent evidence on the role of PV interneurons, which are strongly coupled to each other (Campagnola et al., 2022), in balancing excitation (Palmigiano et al., 2020). Consistently with experimental evidence from V1 (Veit et al., 2017), inserting a third population of SOM interneurons in the E-PV circuit can recover the PING oscillation through its bidirectional coupling with the pyramidal cells, while the connection from SOM to PV cells strengthens a non-oscillatory loop. Further increasing the latter connection, however, gives rise to a different type of oscillation, with a two-dimensional focus. Experimental manipulations of synapses from SOM to PV would help identify which mechanism underlies oscillations in V1. However, we stress that the second mechanism (high SOM→PV) appears less plausible as an explanation for gamma oscillations in V1, because it would require the E-SOM subsystem to have a fixed point with finite values, and some arguments exist against this. Firstly, Veit et al. (2017) have shown that total suppression of the PV population causes an explosion of activity, and, secondly, the fact that PV interneurons show paradoxical responses to stimulation (Palmigiano et al., 2020) suggests that the E-SOM subsystem has a negative determinant and therefore that the oscillation focus blows up. Such a mechanism, however, might underlie gamma oscillations in different areas, or even different dynamics: since it relies on one I population being suppressed at the fixed point of an E-I-I network, it is conceptually analogous to the “sharp-wave” state postulated by Evangelista et al. (2020) for region CA3 of the hippocampus. In a circuit with two mutually inhibiting interneuron types, instead, strong cross-inhibition produces a bistability between two PING oscillations, which could underlie the sudden transitions between different gamma rhythms in the hippocampal CA1 subfield, as hypothesized by Keeley et al. (2017).

Bistability can also emerge between two excitatory populations or assemblies competing through the interneurons. Namely, when the two E populations receive a comparable background input, the relative strength of self- and cross-excitation determines whether each E population excites, inhibits, or fully suppresses the other one. If both populations suppress each other, the network is bistable. Determining which of these three cases applies to each E population could be a standard approach for investigating networks with two or more distinct and homogeneous populations of E neurons. In case interneurons cluster into assemblies that are more strongly excited and more strongly inhibit the E assembly co-tuned to the same stimulus (“balanced” inhibition), we proved that competition between the two E-I modules is only possible if I assemblies inhibit each other less than they inhibit the competing E assembly. SOM interneurons always meet this condition, while for PV interneurons it depends on the assembly-specificity of I→I connections, which is still largely unmeasured. On the contrary, if the I assemblies project more strongly to the not co-tuned E population (“lateral inhibition”), winner-take-all bistability between the E–I modules can appear even when E→E and I→I connections are completely unspecialized, meaning that there are no E and I assemblies in the traditional sense. When I→I cross-inhibition is absent, as it would be for assemblies of SOM interneurons, even slightly lateral inhibition would be sufficient for bistability. Since competitor-suppression is in general a useful feature for E-I assemblies, we predict that some interneuron types should specialize in lateral inhibition, possibly alongside interneuron types specializing in balanced inhibition to stabilize neural activity. This prediction would match the functional specialization predicted for SOM and PV interneurons, respectively, by the plasticity rules experimentally determined by Lagzi et al. (2021).

The theoretical foundation of this study has relied and expanded on previous work by other theoreticians. Namely, Curto and Morrison (2016) and Curto et al. (2019) have comprehensively characterized multistability in TLNs, but only limited to all-inhibitory networks and not in the perspective of explaining the mean-field dynamics of neural populations. Palmigiano et al. (2020); Miller and Palmigiano (2020) and Waitzmann et al. (2024), instead, have given precise mathematical criteria to determine the effects of perturbing specific neural populations, but only in the mono-stable case in which all those populations are active and do not oscillate.

Compared to the most widespread model of the same complexity, the Wilson-Cowan model with a sigmoidal activation function (Wilson and Cowan, 1972), TLNs not only allow for an increased analytical tractability and interpretability, but are also closer to the dynamics of real neural populations in the physiological regime, as they do not artificially introduce an additional stable “up-state”, a saturation of the firing rate which mostly only occurs in epileptic or other unphysiological circumstances. Support for coarsely threshold-linear activation functions of neural populations has come in recent years from experimental measurements (LaFosse et al., 2024), and such functions are increasingly used in models (Sadeh and Clopath, 2020, 2021). Future work needs to assess the extent to which the microcircuit dynamics that we explored in this simple model can be found, under equivalent conditions, also in spiking networks. Some parameters of TLNs, such as the above-rheobase input and the connection strengths, can be directly calculated from the parameters of spiking networks, while parameters like gains and time constants can be easily incorporated into the TLN theory without altering the analytical tractability. In this framework, TLNs could be employed as a roadmap to find specific dynamics of interest within the parameter space of more complex spiking networks, and to explain their dynamics in both mathematical and biophysical terms.

In order to expand on the present study, we indicate several possible directions. Firstly, a large body of theoretical studies has shown that multistability easily turns into a slow oscillation, in the presence of adaptive mechanisms. For example, I-I bistability can give rise to stochastic oscillations associated with perceptual bistability (Laing and Chow, 2002; Shpiro et al., 2007, 2009), and E-I-I bistability was proposed to produce alternations between sharp waves and inactive states in the resting hippocampus (Evangelista et al., 2020). Expanding TLNs with a slow variable representing an adaptive feedback would be a promising way to investigate such hypotheses in an analytically tractable way. Secondly, the model could be complemented by synaptic delays, which would allow capturing additional oscillatory phenomena like ING oscillations (Van Vreeswijk et al., 1994; Wang and Buzsáki, 1996). This could be done either with delay-differential equations or with multiple synaptic filters introducing an implicit delay, as was done with the sigmoidal model by Lu and Rinzel (2024). Lastly, many additional network phenomena can be investigated and understood within the TLN model even in its basic version, including the effects of varying inputs, gains (neuromodulation), or time constants. We believe that the number of microcircuits worth analyzing within this framework is only going to increase in the upcoming years, due to the developments of optogenetics and recording techniques.

## Acknowledgments

We thank Gaspar Cano, Atilla Botond Kelemen, John Rinzel, and Archili Sakevarashvili for helpful discussions, and Gaspar Cano and Liz Weerdmeester for carefully proofreading the manuscript. This work was supported by the Deutsche Forschungsgemeinschaft (DFG), project number 327654276 — SFB 1315. The authors declare that they have no competing interests. All data needed to evaluate the conclusions in the paper are present in the paper and/or the Supplementary Materials. All code used to generate the simulations and figures will be deposited on Zenodo upon publication.

## Author contributions

Conceptualization: S.M. Methodology: S.M. Software, formal analysis, and visualization: S.M. Writing — original draft: S.M. Writing — review & editing: S.M., R.K. Supervision: R.K.

## Materials and Methods

### Fixed Points, Bifurcations, and Stability

We consider a general threshold-linear network (TLN), where the firing rates *x* = (*x*_1_, …, *x*_*n*_) of *n* populations evolve according to the equation

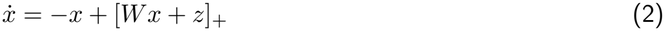

where *z* = (*z*_1_, …, *z*_*n*_) is the vector of the external inputs, *W* is the *n × n* coupling matrix, with entries *w*_*ij*_ (coupling from population *j* to population *i*), and [*x*]_+_ = max{*x*, 0} is the threshold-linear activation function.

It is enlightening to partition the domain of the TLN in two different ways. Firstly, for every *σ ⊆ {*1, …, *n*}, we can indicate as *X*^*σ*^ = {*x ∈* ℝ^*n*^ | *x*_*i*_ *>* 0 *∀i ∈ σ ∧ x*_*i*_ = 0 *∀i ∉ σ*} the subspace in which all and only the populations with an index in *σ* are active. Secondly, we indicate as *L*^*σ*^ = {*x ∈* ℝ^*n*^ | *Σ* _*j*_ *w*_*ij*_*x*_*j*_ + *z*_*i*_ *>* 0 ∀*i* ∈ *σ* ∧Σ_*j*_ *w*_*ij*_*x*_*j*_ + *z*_*i*_ *<* 0 *∀i ∉ σ*} the subspace in which all and only the net *inputs* received by the populations in *σ* are positive. The second partition excludes the hyperplanes *H*_*i*_ = {*x ∈* ℝ^*n*^ | Σ_*j*_ *w*_*ij*_*x*_*j*_ + *z*_*i*_ = 0}, where the net input received by population *i* is exactly zero and Equation (2) is not differentiable. These hyperplanes mark the borders between the different *L*^*σ*^, in each of which the system is linear and can be written as

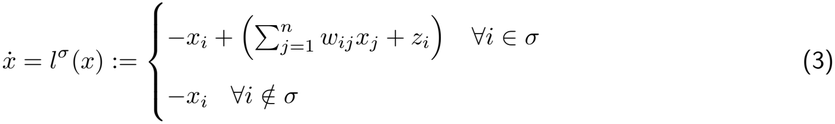

where we indicate each of these linear systems as *l*^*σ*^(*x*). Therefore, over each *L*^*σ*^, the Jacobian is constant, and we can denote its value as *J*^*σ*^ with an *upper* index. In particular, we will often refer to the negative of these Jacobians, which we indicate as 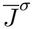 and has two types of rows:

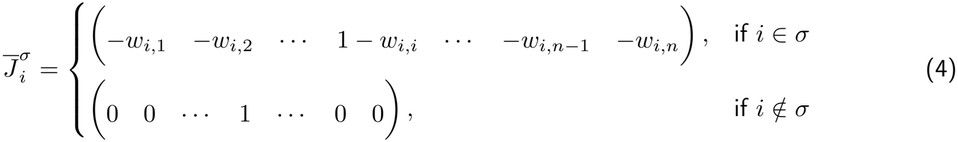

This matrix has the same determinant as the matrix that we obtain by re-arranging rows and columns so that those in *σ* come first, which is

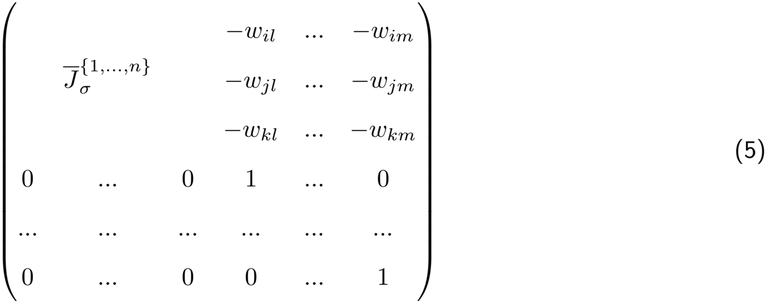

where with the *lower* index *σ* we indicate the restriction of a matrix to the rows and columns in *σ*, and where *i, j, k ∈ σ* and *l, m ∉ σ*. We can see that the determinant of this matrix (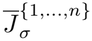 valid in the region *L*^*σ*^) is simply the determinant of 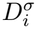, (the negative of the Jacobian of the full system, restricted to *σ*). Therefore, from now on we indicate both determinants with the common notation *D*^*σ*^.

In addition, we employ the notation 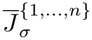 to indicate the determinant of the matrix obtained by replacing the *i*-th column of 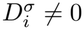 with the vector *z* (restricted to *σ*) of the external inputs. Following Curto et al. (2019), we say that a system is non-degenerate if two conditions are met:

- *D*^*σ*^ *≠* 0 for each *σ ⊆ {*1, …, *n*}
- 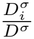 for each *σ ⊆ {*1, …, *n*} and for each *i ∈ σ*

The first condition guarantees that all the solutions of the systems *l*^*σ*^(*x*) exist, while the second one guarantees that they do not lie on the hyperplanes *H*_*i*_. Such conditions have probability 1, so we can assume that they are always met.

Reasoning like Curto and Morrison (2016), Curto et al. (2019), and Curto et al. (2020), it is possible to prove that Equation (2) has a fixed point in the subspace *X*^*σ*^ if and only if the solution of the linear system *l*^*σ*^ lies within its validity region *L*^*σ*^, which happens if and only if

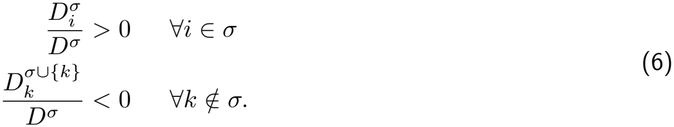

These conditions guarantee that none of the active populations suppresses the others (first set of conditions), while all of the inactive populations get suppressed by the active ones (second set). Several corollaries ensure that, if such a fixed point exists, then it is the only fixed point in *X*^*σ*^, and that a fixed point in *X*^*σ*^ must be also inside *L*^*σ*^. Therefore, since this is the only fixed point that there can be in either *X*^*σ*^ or *L*^*σ*^, we directly indicate it as *σ*. Due to Cramer’s rule, the value of population *i* in the fixed point *σ* is given by 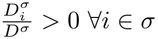. These results have been proved by Curto and Morrison (2016) and Curto et al. (2019) for all-inhibitory TLNs, but the fact that they are inhibitory was not used in the proofs, which thus hold for any TLN.

While the ratios in Equation (6) allow one to determine the existence and the value of the fixed points, the denominators *D*^*σ*^ are related to their stability. In particular, any stable fixed point needs to have a Jacobian in which all the eigenvalues have a negative real part, and therefore the negative of the Jacobian should have all eigenvalues with a positive real part, thus a positive determinant. We have shown already (below Equation (5)) that the determinant of the negative of the Jacobian of the system in any subspace coincides with the determinant *D*^*σ*^ of the restriction of the negative of the full Jacobian to that subspace, so *D*^*σ*^ needs to be positive in order for *σ* to be stable. So, all the fixed points for which both terms in the ratios 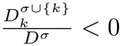 are negative, are unstable, namely saddles. As a consequence, if *σ* and *σ ∪ {k*}, with *k ∉ σ*, are both fixed points, then at least one of them must be unstable, because in the conditions for there is 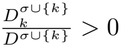, but in the conditions for there is 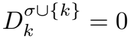, so *D*^*σ*^ and *D*^*σ∪{k*}^ must have opposite signs. This also means that, if two such fixed points exist, they annihilate in a saddle-node (SN) bifurcation where 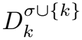. Instead, if *D*^*σ*^ and *D*^*σ∪{k*}^ are both positive (or both negative), and all the conditions for *σ* are met, then *σ ∪ {k*} cannot exist, but it comes into existence when 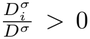 turns from negative to positive. In this way, the fixed point *σ* becomes *σ ∪ {k*}. In this way, *σ* can smoothly transition into *σ ∪ {k*} without changing its stability (but it is also possible that one of the two points is stable and the other one unstable - just, with an even number of unstable modes).

As *D*^*σ*^ *>* 0 is a necessary but not sufficient condition for *σ* to be stable, we need additional requirements, which need to be evaluated for the Jacobian restricted to *σ*. For a 2-dimensional subsystem, the additional condition is that the trace of the Jacobian is negative. In 3- and 4-dimensional systems, closed-form expressions for the stability conditions can be obtained by calculating the eigenvalues through Cardano’s and Ferrari’s solutions for equations of order 3 and 4, but such solutions do not exist in higher dimensions. However, a powerful aid to determine the stability of any fixed point in any *n*-dimensional network comes from the Routh-Hurwitz criterion (Routh, 1877): it is sufficient to calculate the coefficients of the characteristic polynomial of the Jacobian (which in general will be sums of minors) in the subsystem *l*^*σ*^(*x*), and to construct the corresponding Hurwitz matrix. *σ* is stable if and only if all the leading principal minors of the Hurwitz matrix are positive.

### Perturbations

From the conditions outlined so far, we obtain “for free” a perturbation theory. As explained in the previous section, the TLN in Equation (2) has a fixed point *σ*, in which population *i* has the value 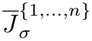 if and only if the conditions in Equation (6) are met. Importantly, the determinant 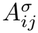, in the numerator of Equation (6), can be Laplace-decomposed as

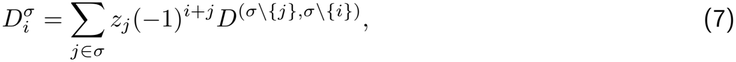

where (*−*1)^*i*+*j*^*D*^(*σ\{j*},*σ\{i*})^ are the (signed) minors obtained by removing the *j*-th row and *i*-th column from 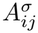. This expansion provides several fundamental insights, which are outlined in the following separately for stable and unstable fixed points.

If we are in a stable fixed point (*D*^*σ*^ *>* 0 in the denominator of Equation (6)), and if we deliver a perturbation to population *j*, therefore changing *z*_*j*_, then the effect of this perturbation scales with the signed minor (*−*1)^*i*+*j*^*D*^(*σ\{j*},*σ\{i*})^. Therefore, provided that the perturbation is small enough not to push the system into a different fixed point, it is sufficient to calculate the sign of these minors to know whether the perturbation will excite or inhibit any other population. This result has been employed by multiple studies adopting a linear activation assumption in order to model perturbations in a widespread “canonical” circuit with three classes of interneurons (Palmigiano et al., 2020; Richter and Gjorgjieva, 2022; Wu and Gjorgjieva, 2023; Waitzmann et al., 2024). As the matrix with such minors as entries is known in linear algebra as the adjugate matrix, we simplify the notations by indicating (*−*1)^*i*+*j*^*D*^(*σ\{j*},*σ\{i*})^ as 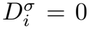.

If instead *D*^*σ*^, appearing at the denominator in Equation (6), is negative (so the point *σ* is a saddle), then the same 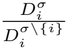 which results in positive (negative) perturbation responses in the stable case produces negative (positive) responses. This effect can be understood in the following way: if the point *σ* is a saddle, we have seen in the previous section that it can have a saddle-node bifurcation with the point *σ \{i*}, in case the latter exists and is stable, and that this happens when 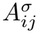. The ratio 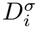 is negative, signifying that *i* is suppressed in the point *σ \{i*}. If a term 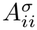, appearing in 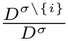, is positive, this indicates that population *j* works for the liberation of population *i* from suppression in the point *σ \ {i*}. At the same time, it makes sense that population *j* has a negative effect on *i* in the saddle *σ*, because the value of *i* in the saddle needs to decrease to 0 in order for it to have a saddle-node bifurcation with *σ \{i*}, resulting in the liberation of *i* from suppression. Therefore, these perturbation responses are consistent with the fact that, if *σ* were stable, *j* would have a positive effect on *i*.

This elegant solution to the perturbation problem (Equation (7)) implies that the response 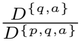 of a population *i* to its own perturbation scales like the determinant *D*^*σ\{i*}^ of the system without *i*. This scaling might appear counterintuitive, but it accounts for the fact that the effect of *z*_*i*_ is divided by *D*^*σ*^, the determinant accounting for the overall self-interactions in the system *σ*, but is not directly affected by the terms appearing in *D*^*σ\{i*}^, as these express self-interactions not involving *i*.

The scaling of the input *z*_*i*_ by the determinant *D*^*σ\{i*}^ is at the origin of the “paradoxical” effects: as highlighted by Miller and Palmigiano (2020), a population *i* has a paradoxical response to its own perturbation (its firing rate decreases when injecting a positive current) if and only if the minor *D*^*σ\{i*}^ is negative, so the paradoxical effect implies that the network without *i* is unstable. As pointed out by Miller and Palmigiano (2020), the converse implication does not hold: if the network without population *i* is unstable, population *i* can also respond non-paradoxically, if (and only if) the Jacobian without *i* has an even number of real positive eigenvalues. Moreover, we shall dispel another possible misunderstanding: the statement “the network without population *i* is unstable” might evoke in the reader the association with a specific kind of instability: the one in which excitatory activity would grow unbounded and inhibitory feedback is needed to control it, as in the most prominent and well-studied example of a paradoxical effect (Tsodyks et al., 1997). However, the dynamical possibilities reach far beyond that: an unstable state might for example consist of two inhibitory populations with strong reciprocal connections, so that they cannot be simultaneously active (see also I-I section of the Results). An excitatory population *p* might stabilize this network by providing the suppressed population with enough excitation to activate (Figure S5A). Stimulating *p* would result in a paradoxical reduction of its firing, because it would drive the growth of this additional inhibitory population. An excitatory population can also stabilize a network in the traditional way: for example, let us consider a network with an excitatory population *p* and an inhibitory population *a*, which is unstable, because the feedback loop is not strong enough to control the runaway excitation (Figure S5B). If an additional excitatory population *q* receives a strong drive from *p*, and in turn drives *a*, then it can help close the loop and stabilize the activity of *p*. Stimulation of *q* excites *a*, which inhibits *p*, which is the main driver of *q* itself, whose firing rate therefore decreases. Excitation-stabilization can therefore also exist and, as much as inhibition-stabilization, is a complex and multifaceted phenomenon that cannot be reduced to the mere prevention of runaway activity.

Once we accept that the effect of a perturbation *z*_*i*_ on the population *i* which receives it (in a fixed point *σ*) is indeed given by 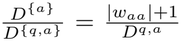, the other response terms can be interpreted in a very intuitive way. For example, we again consider a network with two excitatory populations *p* and *q*, and one inhibitory population *a* (Figure S5C). According to the formula, the effect on *q* of an input *z*_*p*_ delivered to *p* is (see also the section about E-E-I networks)

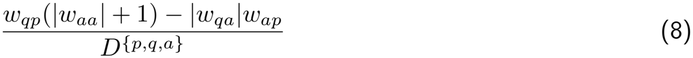

It is easy to see that the sign of this response term is given by the difference between the strength of the excitatory *p → q* pathway and that of the inhibitory *p → a → q* pathway, but it is not immediately clear why the former scales with |*w*_*aa*_| + 1. This becomes clear by decomposing the excitatory path in Equation (8) into 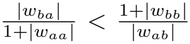 (effect of *z*_*p*_ on *p* itself) times *w*_*qp*_ (strength of the *p → q* pathway), times 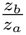 (effect of this excitation on *q*), seen as an external input to the subsystem without *p*. The inhibitory term, reasoning in the same way, can instead be decomposed as

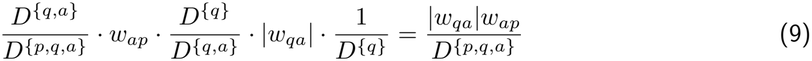

In this case, since this pathway comprises the whole network, all the normalization terms cancel out and the final result is simply divided by the determinant of the whole system.

In case the point {*p, q, a*} is stable, the denominator of Equation (8) is positive, and so the sign of the response term is simply given by the numerator *w*_*qp*_(|*w*_*aa*_| + 1) *−* |*w*_*qa*_|*w*_*ap*_. As one could expect, this term grows monotonically with *w*_*qp*_, so increasing this connection can only change the sign of the response term from negative to positive. In this sense, we can think of this connection as having a positive impact on the *p → q* interaction. However, as an important general remark, this does not necessarily mean that *w*_*qp*_ always increases the response term (8), because it also affects the determinant at the denominator: therefore, it might have a paradoxical negative effect on the perturbation. In particular, if the numerator is negative, and *w*_*qp*_ decreases the denominator (which we assume to be positive), then *w*_*qp*_ makes the numerator less negative and the denominator smaller, and its overall effect can be negative. In the determinant at the denominator, *w*_*qp*_ is scaled by |*w*_*pa*_|*w*_*aq*_ *− w*_*pq*_(|*w*_*aa*_| + 1), which is positive when *q* in turn has a negative effect on *p*: cross-excitation has an ambiguous effect when one population (*p*) inhibits the other (*q*), but not vice-versa. This happens because an increase of *w*_*qp*_ stimulates *q*, but this in turn excites *p*, which inhibits *q*. Figure S6 shows an example of a network in which increasing *w*_*qp*_ can either switch the perturbation sign from negative to positive, or “paradoxically” make it more negative. *w*_*qq*_, which does not appear in the numerator in Equation (8), independently affects the denominator, reducing its positivity: therefore it determines whether the impact of *w*_*qp*_ is stronger on the numerator or on the denominator.

### Limit Cycles

Trajectories in a TLN do not always converge to a stable fixed point. All-inhibitory TLNs with at least three populations can exhibit limit cycles, while with more populations they can even have chaotic attractors, as shown by Morrison et al. (2024). When introducing excitation, limit cycles are already possible in an E-I network, while E-I-I and E-E-I can have multiple ones. Limit cycles can either appear from saddle-node bifurcations when a change in inputs or connections turns some population on or off, or from Hopf bifurcations when a fixed point loses stability as an effect of changes in the connectivity. Hopf bifurcations occur when the real part of a pair of complex eigenvalues changes its sign from negative to positive and they can be found by a set of conditions based on Hurwitz’s criterion (Liu, 1994). We will discuss this criterion in the 2- and 3-dimensional cases. In the 2-dimensional E-I TLN, it can also be proved that this Hopf bifurcation indeed produces a limit cycle (Tsodyks et al., 1997), while in the 3-dimensional TLN we rely on numerical simulations to show that on the other side of this Hopf bifurcation there is indeed a limit cycle (rather than an outward spiral or chaotic attractor).

### Divergence

In TLNs with some excitatory units, trajectories can also diverge, which in our framework we interpret as the onset of unphysiological epileptic-like activity. In the divergent regime, TLNs lose their value as tools to understand biological networks, since the conditions that determine an epileptic saturation of activity are not the same that determine divergence in a TLN: for example, in a TLN a fixed point can never blow up due to an increase of the external input, while biological networks can saturate due to it. In order to understand saturation, it would be interesting to consider a variant of the TLN in which the activation function has an additional upper threshold, beyond which it becomes flat again. This framework would allow for a fully analytical description of the conditions under which saturated fixed points appear, including distinctions between different saturated points (for example, combinations for which one population might saturate, while another one is suppressed, or takes an intermediate value). However, this characterization is beyond the scope of this paper.

In order to understand just some fundamental features of divergence in a TLN, we present a basic criterion. Trajectories can diverge within one region *L*^*σ*^ only if the largest eigenvalue is real and positive, and the eigenvector associated with it has only components with the same sign. This is because otherwise trajectories cannot stay within *L*^*σ*^ when time goes to *∞*. Nevertheless, this is not a sufficient criterion for divergence, because, even if it is verified, trajectories can still cross the border to a different *L*^*σ*^, by activating or inactivating one population, and in that region they might end up at an attractor. In addition, there might also be ways to diverge without permanently staying within the same *L*^*σ*^, but rather moving back and forth between two or more of them. Solving these complications requires additional theoretical work — however, in our simulations the eigenvalue-eigenvector criterion has always been sufficient to identify divergent regimes.

### Simulations

All simulations were performed by integrating the respective dynamical system with the Euler method and a step size of 0.01 units. Each simulation panel reports a 200 *×* 200 parameter sweep. The existence and stability of each fixed point was determined by numerical evaluation of the conditions in Equation (6). Oscillatory states were identified through a “coherence” metric defined as

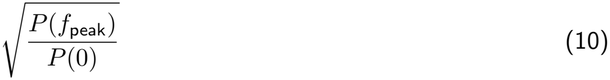

where *P* (*f* ) is the power spectral density and *f*_peak_ *>* 0 is the dominant oscillation frequency. As the signal, we used *x*_*p*_ in the E-I, canonical, and E-I-I network.

## Supplementary Materials 1: Supplementary Text

In what follows, we apply the general theory discussed in the Methods to derive conditions and prove results for each of the specific architectures covered in the Results.

### I-I Network

We consider a TLN with two inhibitory populations, *a* and *b*.

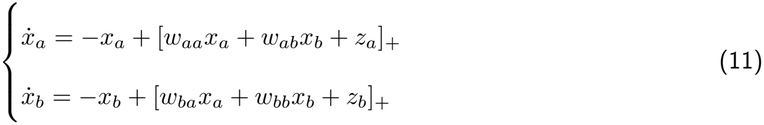

where all the weights are negative. Based on the general solution (6), the fixed point {*a*} exists if and only if

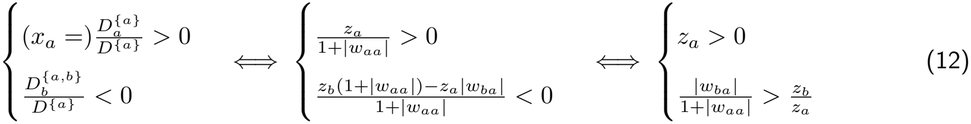

The conditions for {*b*} are completely symmetrical

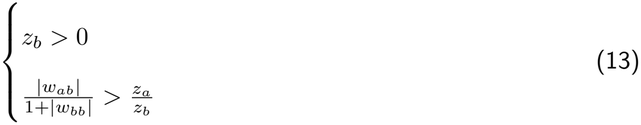

The conditions for {*a, b*} are

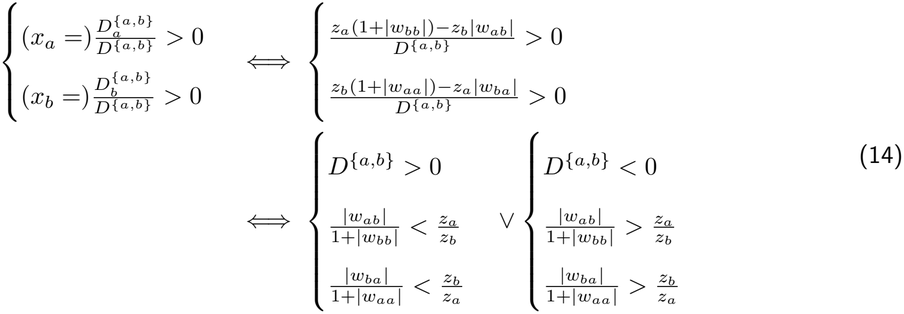

In both cases, both inputs need to be positive. We note that {*a*} and {*b*}, when they exist, are always stable because *D*^{*a*}^ = 1 + |*w*_*aa*_| and *D*^{*b*}^ = 1 + |*w*_*bb*_| are always positive, while {*a, b*} is stable only if *D*^{*a,b*}^ = (1 + |*w*_*aa*_|)(1 + |*w*_*bb*_|) *−*|*w*_*ab*_||*w*_*ba*_| is positive. In addition, we see that {*a*} and {*b*} can only exist in the same network if *D*^{*a,b*}^ *<* 0, because otherwise 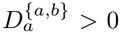, and 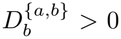 cannot be in between them. Therefore, only these combinations are possible: if *D*^{*a,b*}^ *>* 0, there can either be {*a*} alone, {*b*} alone, or a stable {*a, b*}, depending on whether *a, b* or neither of them suppresses the other population; if *D*^{*a,b*}^ *<* 0, there can either be {*a*} alone, {*b*} alone, or an unstable {*a, b*} co-existing with a stable {*a*} and {*b*}. The negative determinant means that the cross-inhibition between *a* and *b* is stronger than their recurrent inhibitions and leak term, therefore enabling bistability.

The cases that we described correspond to different nullcline configurations in the *x*_*a*_–*x*_*b*_ phase plane (Figure S7). In particular, the linear parts of the nullclines, expressed as functions of *a*, are for the *a*-nullcline

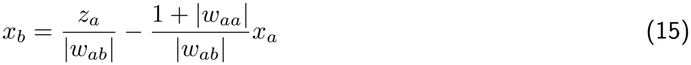

and for the *b*-nullcline

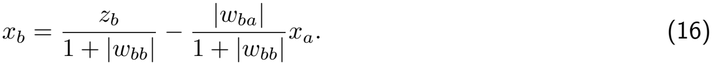

Therefore, *D*^{*a,b*}^ *>* 0 means that the slope of the *a*-nullcline (as a function of *a*) is more negative than the one of the *b*-nullcline, and therefore the *a*-nullcline can meet the *b*-nullcline in the linear part without also meeting it in the zero part; in contrast, if *D*^{*a,b*}^ *<* 0 this is impossible. Moreover, the system moves toward the *a*-nullcline in the horizontal direction, and toward the *b*-nullcline in the vertical direction, so their intersection is only stable if the *a*-nullcline has a more negative slope, or otherwise it is a saddle.

We have said that, when the point {*a, b*} exists and is stable (*D*^{*a,b*}^ *>* 0), {*a*} and {*b*} cannot exist, because 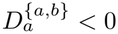 and 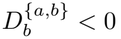 (Equations (12)–(14)). On the contrary, if {*a, b*} exists and if both 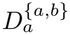 and 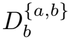, we have shown that *D*^{*a,b*}^ *<* 0. Therefore, if both the suppression conditions 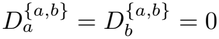 and 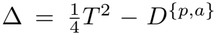 change sign at the same time (like in Figure 1C, when crossing the intersection point from the lower to the upper region along the central axis), the determinant *D*^{*a,b*}^ also needs to change sign. Indeed, if 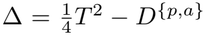, then

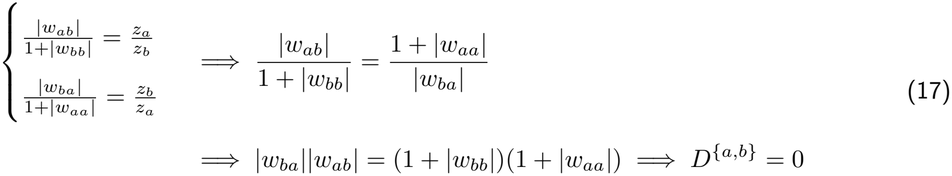

So at this point we have a pitchfork bifurcation, beyond which {*a, b*} keeps existing but becomes unstable and the stable {*a*} and {*b*} appear.

### E-I Network

We consider a TLN with one excitatory population *p* and one inhibitory population, *a*

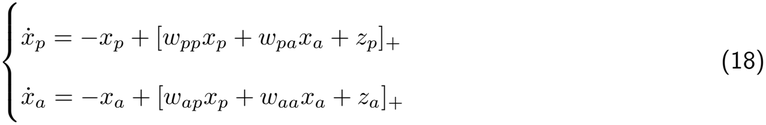

where the weights coming from *p* are positive and those coming from *a* are negative. The linear part of the *a*-nullcline, as a function of *x*_*p*_ is

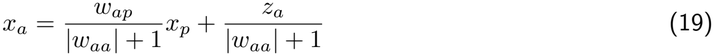

while the linear part of the *p*-nullcline, also as a function of *x*_*p*_, is

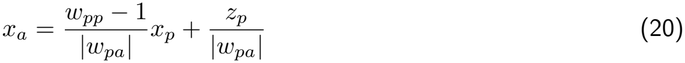

We set apart a case with “subcritical” *p* (*w*_*pp*_ *<* 1) and a case with “supercritical” *p* (*w*_*pp*_ *>* 1). We adopt these terms because the expression “inhibition-stabilized regime” would become ambiguous as soon as we have more than 2 populations. From the nullcline equation, we see that the linear part of the *p*-nullcline is a decreasing function of *x*_*p*_ in the subcritical case, and an increasing function in the supercritical one, which can result in different potential fixed point configurations. In what follows, we will examine the possible fixed point combinations of this network, associating each with the corresponding nullcline configuration in Figure S8. By applying the general theory, we find that the fixed point {*p*} exists if and only if

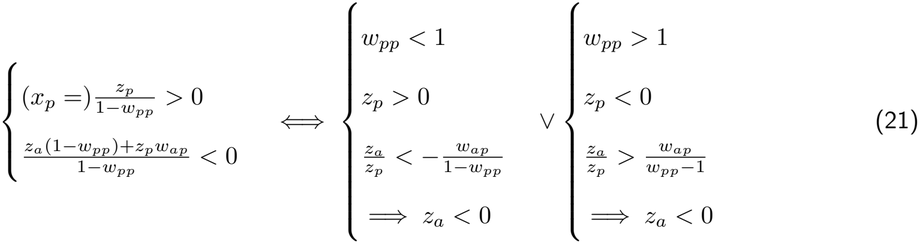

where we distinguish the subcritical case, which needs *z*_*p*_ positive and *z*_*a*_ negative enough for *a* to stay silent in spite of the *p → a* excitation (Figure S8, panel 1), and the supercritical one, which needs *z*_*p*_ and *z*_*a*_ both negative (Figure S8, panel 2). The subcritical case is stable, while the supercritical one is unstable, since the determinant of the negative of the Jacobian in the subspace *L*^{*p*}^ is simply 1 *− w*_*pp*_, while the trace of the Jacobian is *w*_*pp*_ *−* 2. Therefore, a stable {*p*} cannot exist if *p* is supercritical, consistently with the meaning that this population would need to be inhibition-stabilized. Moving forward, {*a*} exists if and only if

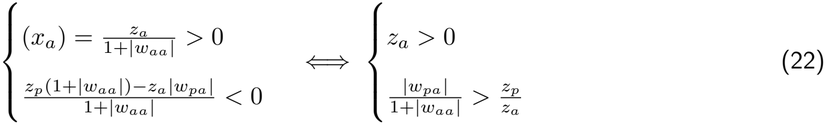

which is a single-population “suppression term”, detailing the condition for one population to suppress another one, analogously to the conditions found in the previous section (“I-I Network”). If {*a*} exists, it is always stable. If {*a*} exists, it cannot co-exist with {*p*} (Figure S8, panel 3). The conditions for {*p, a*} instead are

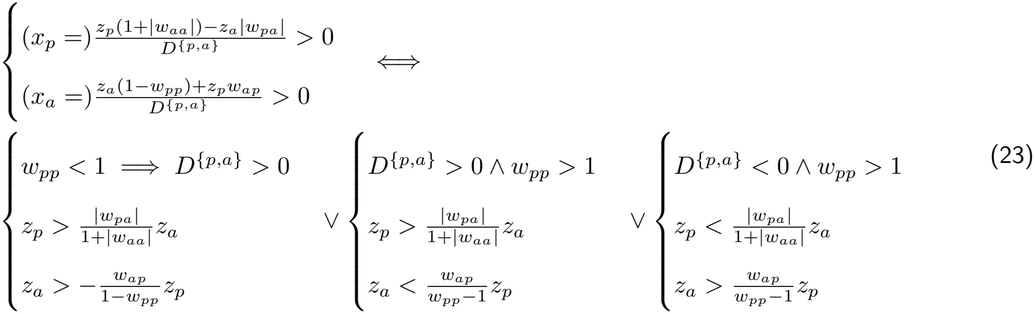

where we have used that the value of *D*^{*p,a*}^ is (1 *− w*_*pp*_)(1 + |*w*_*aa*_|) + |*w*_*pa*_|*w*_*ap*_, so, if *w*_*pp*_ *<* 1, it cannot be negative. We further distinguish five subcases based on the sign of the inputs:

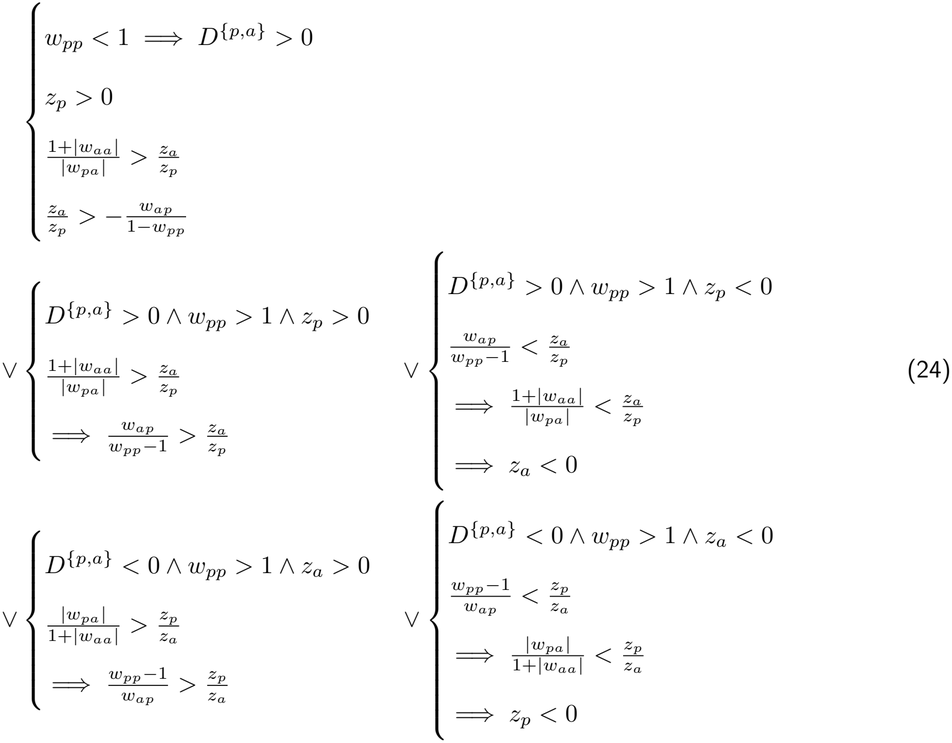

The first case (Figure S8, panel 4) corresponds to a subcritical {*p, a*} state, which is always stable, because *D*^{*p,a*}^ *>* 0 and the trace of the Jacobian is *w*_*pp*_ *−* |*w*_*aa*_| *−* 2 *<* 0. Such a point also cannot co-exist with {*p*} or {*a*}, as its conditions negate one of their conditions. In particular, the input to *a* needs to be strong enough for this population to fire, and weak enough not to suppress *p*. Seen otherwise, *p* must provide enough excitation to *a* and not receive too much inhibition from it. In addition, from the LHSs of the conditions in Equation (23) we can also easily see that delivering a stimulation to any population has exactly the effect that one would non-paradoxically imagine: stimulating *p* increases the rate of both *p* and *a*, while stimulating *a* increases the rate of *a* and decreases the rate of *p*.

The second case (Figure S8, panel 5), instead, produces a supercritical {*p, a*} point, which is stable under the additional condition that the trace of the Jacobian is negative. In this case, the non-suppression condition for *a* follows from the other conditions, since the supercritical {*p, a*} can only have a positive determinant if *a* is also active. In such a point, stimulating *p* increases the firing of both *p* and *a*, while stimulating *a decreases* both of them — the well-known paradoxical effect, that can be clearly appreciated in Equation (23).

The third case (Figure S8, panel 6) also produces a supercritical {*p, a*}, but we distinguish it because this occurs when both inputs are negative. While the supercritical {*p, a*} with positive *z*_*p*_ cannot co-exist with any other fixed point, when *z*_*p*_ (and consequently *z*_*a*_) is negative, this point always co-exists with an unstable {*p*} and, additionally, with the point {*∅*}, in which both rates are 0 (which trivially exists if and only if both inputs are negative). In this case, if {*p, a*} has a negative trace, there is a bistability between {*∅*} and {*p, a*}, while {*p*} is unstable, a case that has been extensively discussed by Jercog et al. (2017) as a candidate dynamical landscape underlying neocortical slow oscillations.

In the fourth case (Figure S8, panel 7), {*p, a*} is also supercritical, but the negative determinant *D*^{*p,a*}^ automatically ensures its instability, and in addition *z*_*a*_ *>* 0, so this point co-exists with a stable {*a*}.

In the fifth case, for *z*_*a*_ *<* 0 (Figure S8, panel 8) also *z*_*p*_ *<* 0, and the unstable {*p, a*} co-exists with the stable {*∅*}.

In addition, it is also possible that no fixed point exists (Figure S8, panel 9). The conditions for this to happen can be obtained by negating all the other combinations at the same time. This only leaves

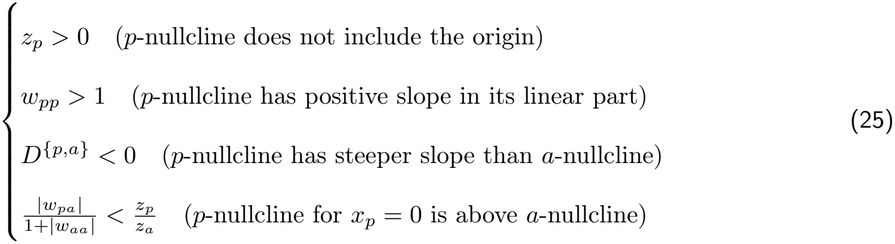

In the last three cases, and in general every time that *w*_*pp*_ *>* 1 and *D*^{*p,a*}^ *<* 0, there are diverging trajectories, even if there are additional stable points {*a*} or {*∅*}. In fact, in this case there is one positive and one negative eigenvalue (which are real, since the discriminant 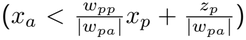 is necessarily positive), and the eigenvector associated with the positive eigenvalue is

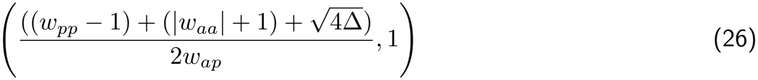

where 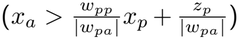 is the discriminant, and *T* = *w*_*pp*_ *−* |*w*_*aa*_| *−* 2 the trace of the Jacobian (note that *D*^{*p,a*}^, defined as the determinant of the negative Jacobian, is also the determinant of the Jacobian, because this fixed point is 2-dimensional). Not only both components of this vector are positive, so the necessary criterion for divergence within *L*^{*p,a*}^ is satisfied, but we can also ensure that trajectories that start far enough from hyperplanes *H*_*a*_ and *H*_*p*_ stay within *L*^{*p,a*}^. In fact, the equation of *H*_*p*_ is

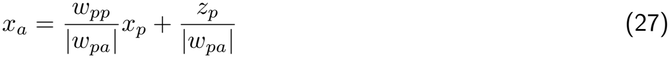

and this hyperplane separates the region in which *p* receives a positive net input 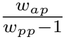 from the region in which it receives a negative net input 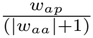. The slope of the relevant eigenvector (in the plane with *x*_*p*_ on the horizontal axis and *x*_*a*_ on the vertical axis) is

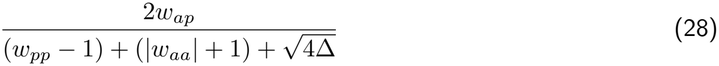

and since *D*^{*p,a*}^ *<* 0 we have 4Δ *> T* ^2^ = (*w*_*pp*_ *−* |*w*_*aa*_| *−* 2)^2^, so

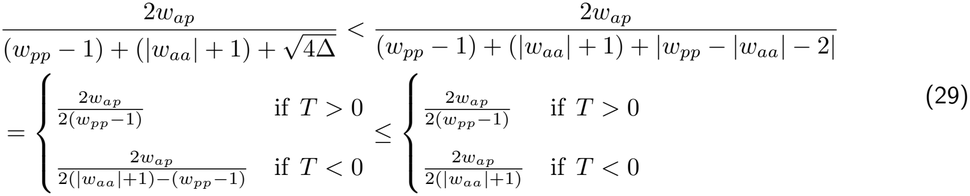

In case *T >* 0, we then have that the slope of the eigenvector is smaller than 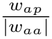. Since *D*^{*p,a*}^ *<* 0, then *w*_*ap*_|*w*_*pa*_| *<* (*w*_*pp*_ *−* 1)(|*w*_*aa*_| + 1), but then, using that *T >* 0

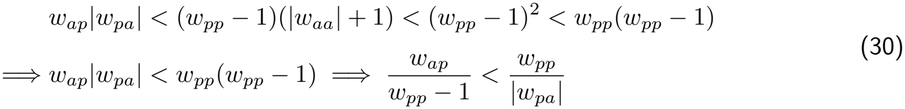

So the term which is larger than the slope of the eigenvector is smaller than the slope of *H*_*p*_, given by Equation (27). If instead *T <* 0, we get that the slope of the eigenvector is smaller than 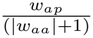. In this case, we can easily conclude that

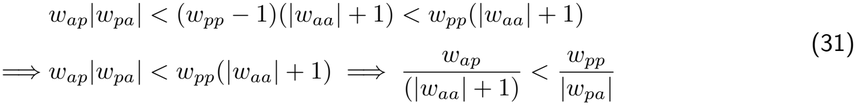

therefore, any trajectory which starts far enough from *H*_*p*_ will never cross it. The same holds for *H*_*a*_, because the slope of *H*_*a*_ is 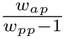, and in case *T <* 0 and the slope of the eigenvector is smaller than 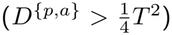, we have that

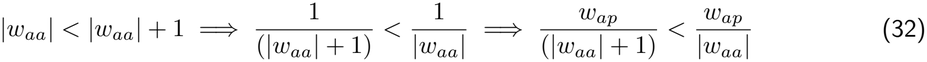

while if *T >* 0 and the slope of the eigenvector is bounded by 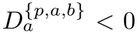, then we can write

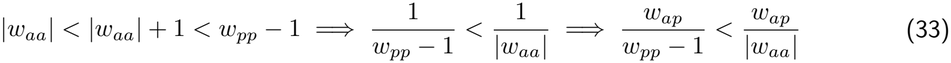

Therefore, any trajectory that starts far enough from *H*_*p*_ and *H*_*a*_ will stay in *L*^{*p,a*}^ forever (because it will become asymptotic to the eigenvector and stay away from both hyperplanes) and diverge. It is possible to prove the same when the supercritical point {*p, a*} exists and has a positive determinant, a positive trace, and a positive discriminant. Otherwise, if the discriminant is negative 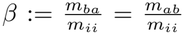, the system has a limit cycle, which is conceptually analogous to a PING oscillation (Tsodyks et al., 1997).

As a conclusion, we wish to prove that recurrent inhibition *w*_*aa*_ broadens the range of “useful inhibition”, for which the rate *x*_*p*_ in the supercritical point {*p, a*} stays within two arbitrary values *x*_*min*_ and *x*_*max*_. This requires (as long as the determinant is positive)

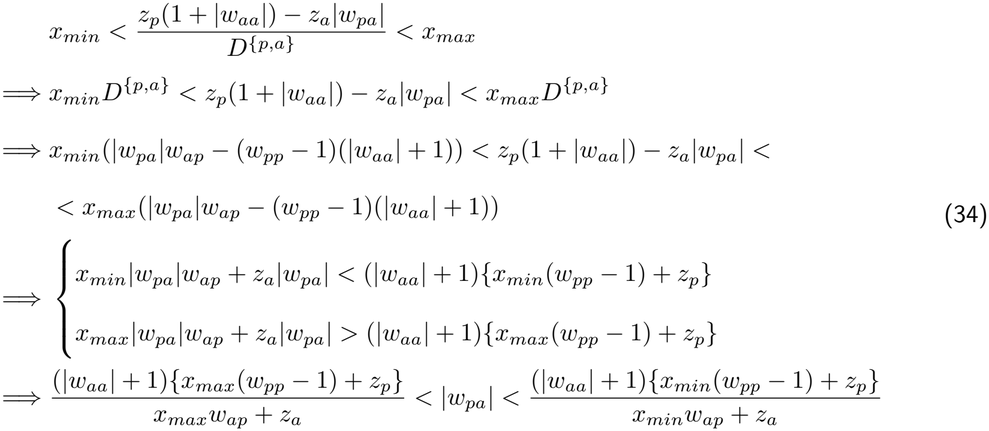

Therefore, the range of values |*w*_*pa*_| that result in a firing rate that is neither too low nor too high has the width

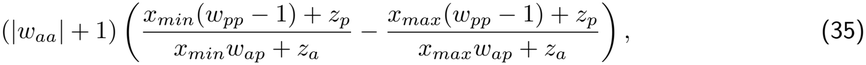

which increases monotonically with |*w*_*aa*_|. Note that we have assumed “supercritical” *p*, i.e. *w*_*pp*_ *>* 1.

### Canonical Circuit

For homogeneity with respect to the other sections of the SM, and unlike the corresponding Results section, here we use the following notations: *p* for E cells, *a* for SOM interneurons and *b* for PV interneurons. We first examine the fixed point {*p, a, b*}, for which the Jacobian is

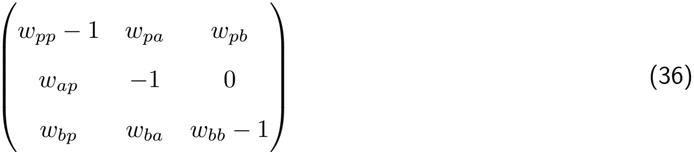

The coefficients of the characteristic polynomial of this matrix (here we take the Jacobian and not its negative, to avoid confusion) are the trace *t*, the cofactor sum *c* (sum of the three 2 *×* 2 principal minors) and the determinant *d*. Hurwitz’s stability criteria are

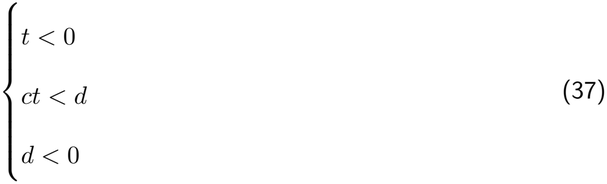

The conditions for a Hopf bifurcation instead are (Liu, 1994)

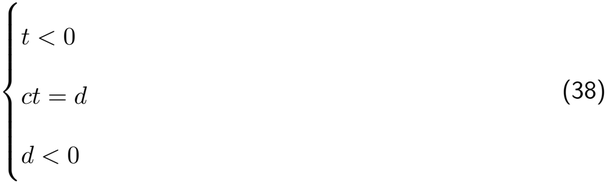

with the complex eigenvalues taking a positive real part when *ct > d*. We will occasionally refer to this condition as the condition for “oscillatory instability”, and simulations confirm that on this side of the Hopf bifurcation there is indeed a (3-dimensional) limit cycle that persists far away from the bifurcation. We now examine in more detail what this condition means, first in a generic *p*-*a*-*b* system, and then in the canonical circuit. Oscillatory instability is achieved when

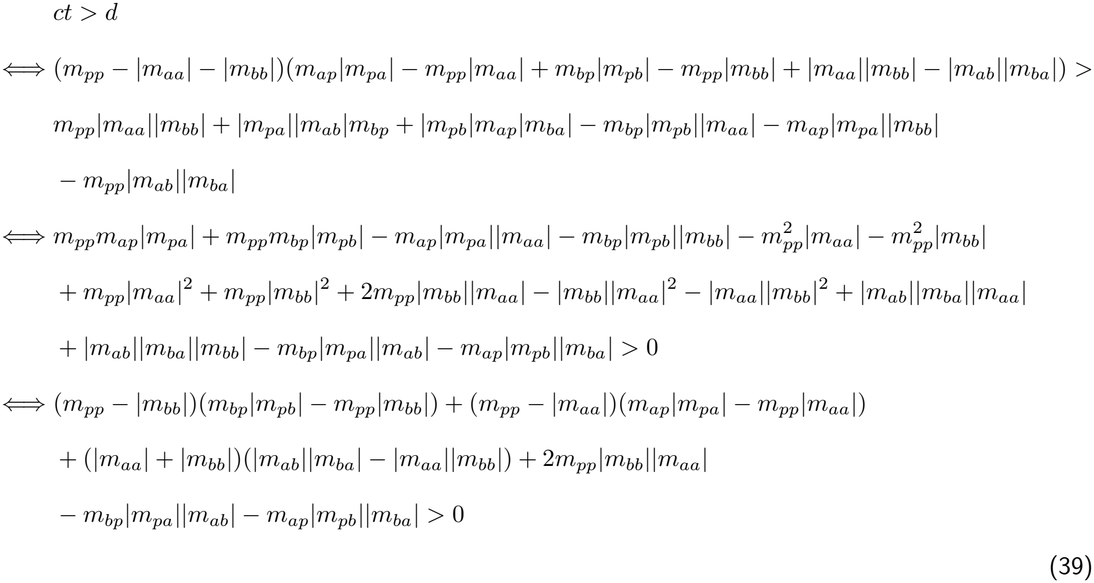

where, to get a more synthetic expression, we use the terms with *m* to indicate the full entries of the Jacobian, including the relaxation, so for example *m*_*pp*_ = *w*_*pp*_ *−* 1, while *m*_*pa*_ is just *w*_*pa*_. This expression summarizes as a positive sum between three terms representing the product of trace and determinant of each subsystem, plus three additional cross-talk terms. For the canonical circuit, the expression simplifies to

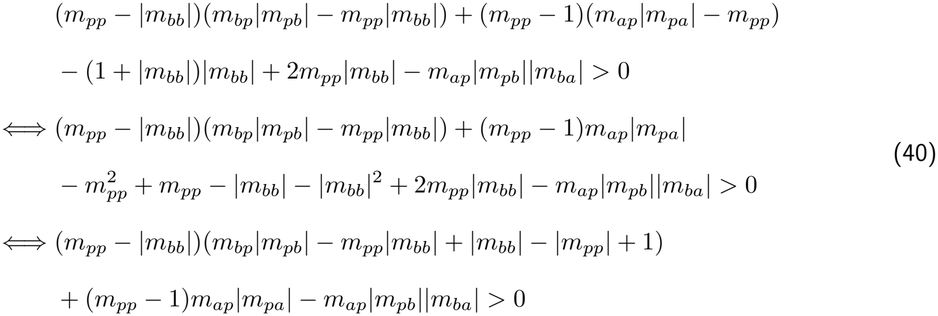

The whole term (*m*_*pp*_*−*|*m*_*bb*_|)(*m*_*bp*_|*m*_*pb*_|*−m*_*pp*_|*m*_*bb*_|+|*m*_*bb*_|*−*|*m*_*pp*_|+1) can be summarized as *T* ^{*p,b*}^(*D*^{*p,b*}^*− T* ^{*p,b*}^ + 1), which is always negative when {*p, b*} is stable (as this implies *T* ^{*p,b*}^ *<* 0 and *D*^{*p,b*}^ *>* 0), while the other two terms depend on the SOM interneurons *a*. Specifically, these interneurons induce oscillations in the system through their connections to the pyramidal neurons, an effect that is amplified by the recurrent excitation *m*_*pp*_ (provided that this is strong enough, but the threshold is much lower than the one required to make the *p*–*b* system oscillatory, due to the lack of recurrent inhibition among SOM interneurons). On the contrary, the connection from *a* to *b* (SOM to PV) hinders the oscillation: the stronger the *a → b → p → a* loop is, the weaker the oscillation. Moving along this loop clarifies why it cannot act as an oscillator: *p* excites *a, a* inhibits *b, p* is disinhibited from *b*, and it excites *a* even more, and so on until a fixed point is reached: no 3-dimensional oscillation can emerge due to this loop.

External inputs do not directly play a role in this bifurcation, but, as seen in Figure 3Ciii–v, they can stop the oscillation by turning off population *p* at the fixed point. This happens when

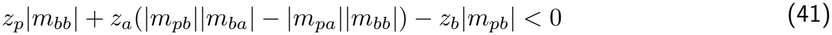

an expression which is greatly simplified by the lack of PV→SOM and SOM→SOM connections. It is straightforward to see that inputs to *p* favor the oscillation and inputs to *b* hinder it, while the effect of inputs to *a* depends on whether the disinhibitory pathway *a → b → p* or the inhibitory pathway *a → p* (scaled by |*m*_*bb*_|) is stronger. However, since the oscillation needs large *m*_*pa*_ and small *m*_*ba*_, and since we assume *m*_*bb*_ to be large, *z*_*a*_ is more likely to have a negative effect on *p* and to ultimately turn off this population when it increases too much (as it does in Figure 3Ciii, top left region, or on the right of Figure S1Ci). Alternatively, the oscillation can also stop when *a* turns off at the fixed point, which happens when

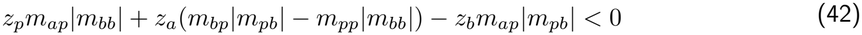

where *z*_*p*_ and *z*_*a*_ always have a positive effect and *z*_*b*_ always a negative one. An example of the oscillation turning off in this way can be seen in the bottom region of Figure 3Ciii or on the left of Figure S1Ci.

A different, 2-dimensional, limit cycle can be achieved when the fixed point {*p, a, b*} is replaced by {*p, a*}: in this case the oscillation is centered in *X*^*p,a*^ and it is essentially a 2-dimensional oscillation, although *b* can still activate in part of the trajectory. Prerequisites for this oscillation to exist are on the one hand that {*p, a*} is in the oscillatory state, on the other hand that its determinant is positive (whether this assumption is reasonable in the context of the canonical circuit is discussed in the Results) and neither *p* nor *a* is suppressed (see E-I section of the SM). In addition, *b* needs to be suppressed at the oscillation focus. The condition for this is

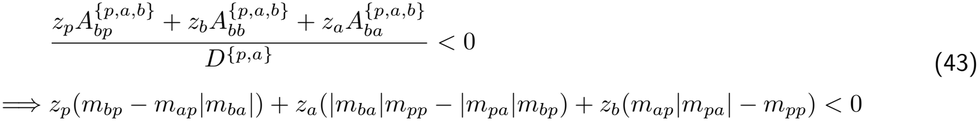

The term scaling *z*_*b*_ is the determinant of the *p*–*a* subsystem, which we assumed to be positive, so this input non-paradoxically works against the suppression of *b*. The sign of term scaling *z*_*p*_ depends on whether the inhibitory *p → a → b* pathway or the excitatory *p → b* pathway is stronger. In this case, it is more likely negative (like in Figure 3Civ) due to the lack of recurrent inhibition in SOM interneurons, which would otherwise multiply the *p → b* pathway because of comparably reducing the efficacy of *p → a → b*. The term scaling *z*_*a*_ is usually negative (as in Figure 3Ciii, S1Ci–ii), but it can turn positive, if *p* is very supercritical, due to the paradoxical excitatory effect of the increased *m*_*ba*_ connection, which can surpass the effect of the *a → p → b* pathway, which is always inhibitory.

In this way, we obtained insights on the effects of the external inputs, while to understand the contributions of single connections, it would help to rearrange this expression in different ways. For example, although *m*_*ba*_ can have a paradoxical excitatory effect within the minor scaling *z*_*b*_ (as we have just discussed), its overall effect on *b* is always negative, as we can see by re-arranging the terms as

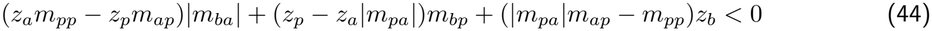

where (*z*_*a*_*m*_*pp*_ *− z*_*p*_*m*_*ap*_) must be negative in order for {*p, a*} to exist at all. In addition, *z*_*p*_ *− z*_*a*_|*m*_*pa*_| also needs to be positive, so the effect of *m*_*bp*_ is always positive. The effect of *m*_*ap*_ scales with *z*_*b*_|*m*_*pa*_|*−z*_*p*_|*m*_*ba*_|, so it becomes more negative with increasing |*m*_*ba*_|, as we can see in Figure 3Cii. The effect of |*m*_*pa*_| scales with *z*_*b*_*m*_*ap*_ *− z*_*a*_*m*_*bp*_, so it is positive as long as we assume, as we do in most default cases, that SOM neurons receive zero or negative external inputs (since SOM neurons are known to be primarily locally driven (Beierlein et al., 2003), and also receive additional inhibition from VIP interneurons). The effect of *m*_*pp*_ scales with *z*_*a*_|*m*_*ba*_| *− z*_*b*_, which again is mostly negative for us.

If we assume that the external drive to SOM is exactly 0, we simplify the expression for this second limit cycle as

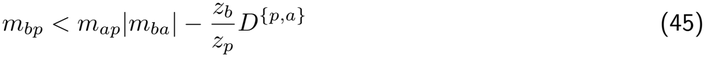

which is a comparison between the excitation that *b* receives from *p* and the excitation that *a* receives from *p*, minus the ratio between the inputs to *b* and *p* times the determinant of the *p*–*a* subsystem.

These conditions inform us about the fixed points, but in order to determine whether limit cycles indeed appear through the bifurcations and are maintained far away from them, we perform a semi-analytical bifurcation analysis in XPPAUT (Ermentrout and Mahajan, 2003). Since the software cannot handle differential equations with a TL activation function, due to non-differentiability at the threshold point, we replace it with its smooth approximation, the so-called softplus function

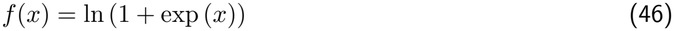

where the argument *x* is the total input. In general, we find an excellent agreement between the two models (see e.g. the comparison in Figure S1A). The most remarkable difference is that the softplus network does not need a population to inactivate completely in order for the 3-dimensional limit cycle to disappear due to varying external inputs (e.g. 3C or S1C), resulting in smoother and more biologically realistic transitions. All the dynamical regimes are otherwise maintained and they are found for the same combinations of parameters. The results of the bifurcation analysis of this system are reported in the Results section.

### Limit Cycle Bistability in E-I-I Circuit

We consider the same system examined in the section “Canonical Circuit”, but under the assumption that *a* and *b* are identical, except for the drives and the reciprocal inhibitions *w*_*ab*_ and *w*_*ba*_. In order to indicate the identical inhibitory connections we use the index *i* (specifically *m*_*ii*_ := *w*_*ii*_ *−* 1). Although general conditions for the existence of different limit cycles in this system cannot be given analytically, a necessary condition in order for population *b* (*a*) to be fully suppressed by a two-dimensional limit cycle created by the other two populations is that the unstable point {*p, a*} ({*p, b*}) exists. In fact, if the whole trajectory lies in *X*^{*p,a*}^, it must enclose an unstable fixed point due to a corollary of the Poincaré–Bendixson theorem, and this point can only be {*p, a*}. Such a point must be of the oscillatory-unstable kind, i.e. it must have positive trace and negative discriminant (and positive determinant). Therefore, a necessary condition for bistability between limit cycles in which the competing population is suppressed is the “bi-instability” of the fixed points {*p, a*} and {*p, b*}. The conditions for one of these points with an oscillatory instability are, for example for {*p, a*}:

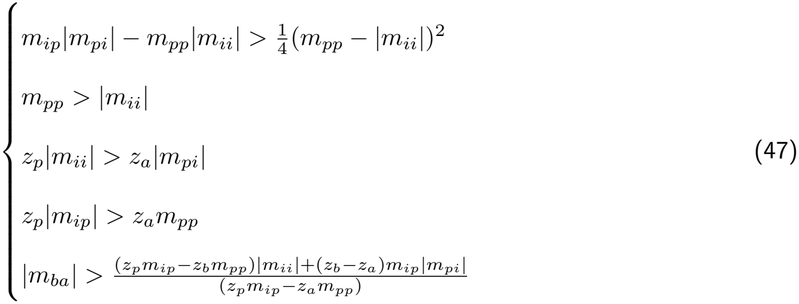

where the first conditions are taken from the E-I network, and the last one is Condition (43) adapted for when *a* and *b* are identical except for the drives and the cross-connections. For {*p, b*} the last condition is

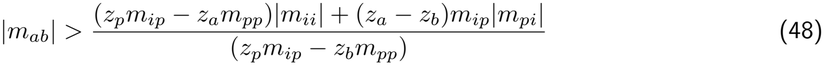

which is 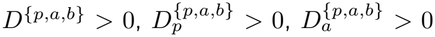. These two expressions indicate that limit-cycle bistability with suppression of the competing population requires strong cross-inhibition in both directions. The existence of the oscillatory-unstable {*p, a*} and {*p, b*}, however, does not guarantee that the whole trajectory of the limit cycles remains two-dimensional, as this could cross the hyperplane that marks the activation of the competing population, if the fixed point is close enough to it. Activation of the competing population does not automatically mean that the bistability is lost and the two cycles merge in a single one: especially if in *X*^{*p,a,b*}^ there is a saddle, with eigenvectors pointing toward *X*^{*p,a*}^ and *X*^{*p,b*}^, then the trajectories can be pushed back to their plane and the bistability be maintained.

If 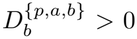 and *z*_*i*_ := *z*_*a*_ = *z*_*b*_, then the two conditions in Equations (47) and (48) are the same and simplify to *β >* 1 (which is the same condition which delivers bistability in the I-I network). The unstable foci {*p, a*} and {*p, b*} both exist for *β >* 1 and neither exists for *β <* 1. In addition, we wish to show that {*p, a, b*} exists on both sides of the bifurcation, but with opposite determinants. The conditions are 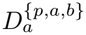, and 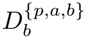 for it to exist with positive determinant, and all the opposite to exist with negative determinant. 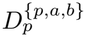 and 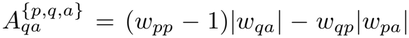 are the same quantities considered above, while for the determinant *D*^{*p,a,b*}^ we get

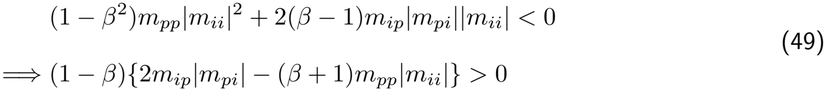

For *β <* 1, *D*^{*p,a,b*}^ is always positive, while it is negative in an interval for *β >* 1, before the second term grows enough to change sign. For 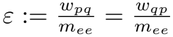 we get a very similar expression:

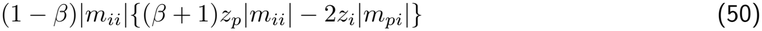

from which we derive the same conclusions. Therefore, at the bifurcation point *β* = 1, all the relevant quantities for {*p, a, b*} switch sign, and the point keeps existing on the other side of the bifurcation. If all the additional stability conditions for {*p, a, b}, {p, a*}, and {*p, b*} are met, this produces a classic pitchfork bifurcation analogous to what we have seen in the I-I model. Otherwise, any of the stable points involved in the pitchfork can instead be unstable and likely surrounded by its own limit cycle. In the semi-analytical bifurcation analyses of the equivalent system with softplus activation functions, we see that this pitchfork-like bifurcation coincides with an analogous bifurcation of limit cycles, both when {*p, a, b*} is stable and {*p, a*} and {*p, b*} are unstable (Figure 4Ei) and when all the involved points are unstable (Figure S2Di). Which one of these two scenarios occurs depends on whether the Hopf bifurcation which stabilizes the 3-dimensional limit cycles occurs for a value of *β* larger or smaller than 1 (we recall that both *w*_*ab*_ and *w*_*ba*_ have a desynchronizing effect, due to their involvement in non-oscillatory loops, Equation (39)). In order to determine this, we rewrite Equation (39) for our symmetrical case

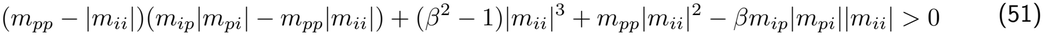

This is still true at *β* = 1 if and only if

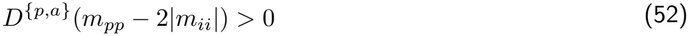

Therefore, since we assumed *D*^{*p,a*}^ *>* 0, we get the direct transition from the 3-dimensional oscillation to the 2-dimensional ones (as in Figure S2A–C) when *m*_*pp*_ *>* 2|*m*_*ii*_|. If instead |*m*_*ii*_| *< m*_*pp*_ *<* 2|*m*_*ii*_|, an intermediate non-oscillatory region emerges (as in Figure 4D).

### E-E-I Network

We consider a general E-E-I system, in which two E assemblies or populations *p* and *q* are connected to unspecific interneurons *a*

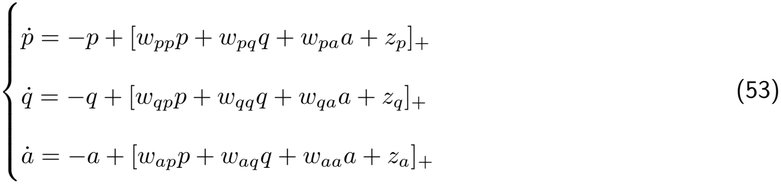

The Jacobian of the full system *L*^{*p,q,a*}^ is

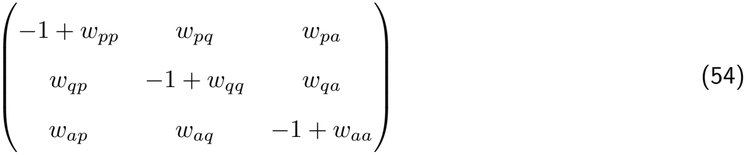

We assume *p* and *q* to be supercritical. Provided that the point {*p, q, a*}, in which all the populations are active, exists, the sign of the effect that an increased input to population *p* has on *q* is given by the entry of the adjugate matrix

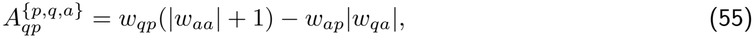

which has been extensively commented on, as a general example in the Methods (section “Perturbations”). One excitatory population can also be completely suppressed by the other one: for example, the stable point {*p, a*}, in which the competing assembly *q* is suppressed, exists when the usual conditions for the stable supercritical E-I point are met, and in addition

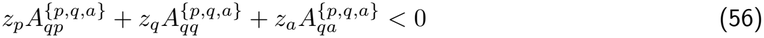

The first term is the already considered minor (55), expressing the competitive or cooperative effect of *p* on *q*. The second is the determinant *D*^{*p,a*}^, which needs to be positive in order for {*p, a*} to be stable.

The last term, 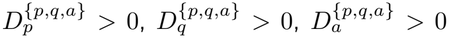, expresses the effect of *a* on *q*, which one would expect to be inhibitory and indeed, if *w*_*pp*_ *<* 1, always is. If *p* is inhibition-stabilized, however, this term could turn positive because a high input to *a* would reduce its firing due to the paradoxical effect, and therefore disinhibit *q*. In order to understand whether the resting state of this network, when *z*_*p*_ and *z*_*q*_ are close to each other, is co-active or bistable, we assume *z*_*e*_ := *z*_*p*_ = *z*_*q*_. We also use the notation *z*_*i*_ for *z*_*a*_ and *m*_*ij*_ for the (*i, j*) entry of the Jacobian. Then the suppression condition for *q* is

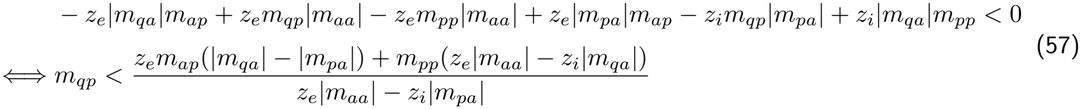

where the denominator and the term *m*_*pp*_(*z*_*e*_|*m*_*aa*_|*−z*_*i*_|*m*_*qa*_|) are positive for supercritical *p*. Ideal conditions for the suppression of *q* are when *w*_*qp*_ is small compared to *w*_*pp*_ *−* 1, and when *a* preferentially targets *q*. However, just one of these combinations is sufficient for suppression, as, for *m*_*qp*_ = *m*_*pp*_ the condition becomes exactly

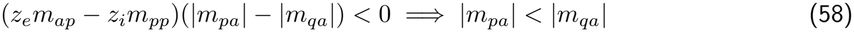

where we used that the first term must be positive. For |*w*_*qa*_| = |*w*_*pa*_|, instead, the condition is just

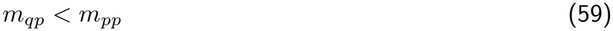

which means that asymmetries in the E→I connections only matter for suppression if there are also asymmetries in the I→E connections — and in this case an increase in either of the E→I connections always works against the population which is preferentially targeted by inhibition.

Now we turn to symmetric networks (with asymmetric drives), where the *p*–*a* and *q*–*a* subnetworks are identical copies, so *m*_*ee*_ := *w*_*qq*_ *−* 1 = *w*_*pp*_ *−* 1, *m*_*ie*_ := *w*_*aq*_ = *w*_*ap*_, *m*_*ei*_ := *w*_*qa*_ = *w*_*pa*_, 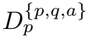 *m*_*ii*_ := *w*_*aa*_ *−* 1. Note that *ε* scales *w*_*pp*_ *−* 1 and not *w*_*pp*_, so *ε* = 1 is achieved for values of the cross-connections *w*_*pq*_ and *w*_*qp*_ which are still smaller than *w*_*pp*_ and *w*_*qq*_. Then the condition for *q* to be suppressed is

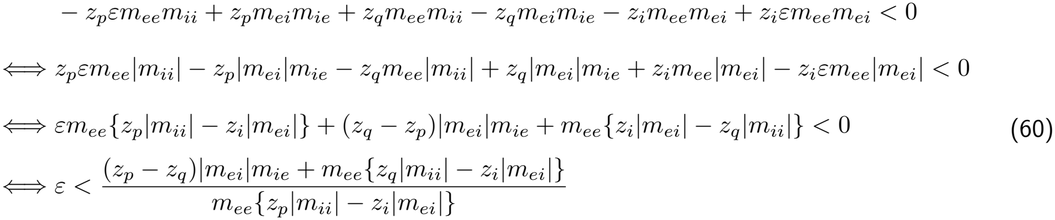

where in the last step we used that *m*_*ee*_{*z*_*p*_|*m*_*ii*_| *− z*_*i*_|*m*_*ei*_|*} >* 0, as one of the other conditions for {*p, a*}. This condition is satisfied when the cross-excitation coefficient *ε* is smaller than a threshold value set by the other parameters. In particular, the threshold is the ratio between the sum of two terms and a denominator which would be *<* 0 if *p* were suppressed by *a* and is the larger the less *a* suppresses *p*, so we refer to this as the binary non-suppression term for *p*. The two terms in the numerator are: the product of the E→I and I→E pathways scaled by the input difference between the two excitatory populations, and the binary non-suppression term for *q*. The condition for the competing point {*q, a*} is instead

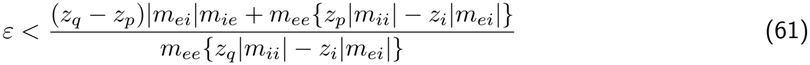

In the special case in which the inputs are the same, like in a resting state, then the conditions for {*p, a*} and for {*q, a*} are the same and simplify to

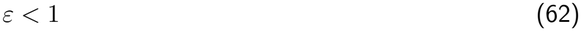

We have shown that both points {*p, a*} and {*q, a*} always exist and are stable (if the other E-I conditions are met), when *ε <* 1, and they both do not exist when *ε >* 1. Now we want to prove that at *ε* = 1 the system has a pitchfork bifurcation, so we need to show that {*p, q, a*} always exists and is stable when *ε >* 1, and that it always exists and is unstable when *ε <* 1. The conditions for this are: 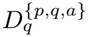 when *ε >* 1, and all the opposite when *ε <* 1. For 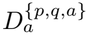 and 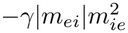, we have just shown that. Now we consider the determinant *D*^{*p,q,a*}^ (determinant of the *opposite* of the Jacobian)

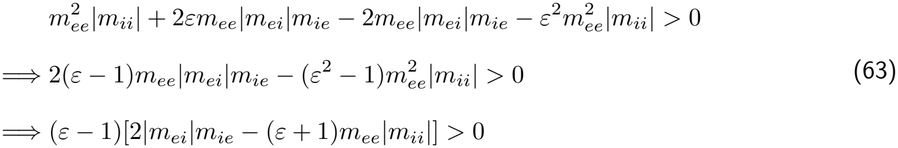

Since *m*_*ie*_|*m*_*ei*_| *> m*_*ee*_|*m*_*ii*_|, the determinant is negative for every *ε <* 1, while when *ε* becomes *>* 1 there is at least an interval in which *D*^{*p,q,a*}^ *>* 0 (until *ε* gets so big that the second term switches sign: this is where the point disappears and activity diverges). For what concerns 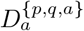, we basically have the same calculations again, just with *−z*_*i*_ instead of |*m*_*ii*_| and the (negative of the) common excitatory drive *−z*_*e*_ instead of |*m*_*ei*_|. Then we get

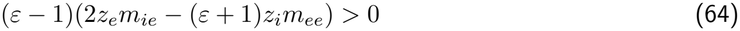

where we can draw the same conclusions since *z*_*e*_*m*_*ie*_ *> z*_*i*_*m*_*ee*_. Therefore, we have proved that this system has a pitchfork bifurcation for *ε* = 1. Above this value, the network then transitions to the co-active regime, but interactions between the two populations are still competitive. The perturbation effect, in the symmetric network with equal drives, becomes positive only when

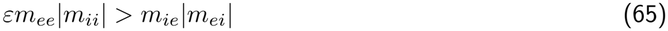

which always happens for a value *ε >* 1, because the E-I determinant must be positive. Therefore, bistability in a symmetric network always requires that the excitatory populations have a negative perturbation effect on each other.

Now, we call *ε*^{*p,a*}^ the threshold below which {*p, a*} exists (Equation (60)), and *ε*^{*q,a*}^ the threshold below which {*q, a*} exists (Equation (61)). When taking the derivative of *ε*^{*p,a*}^ with respect to the input *z*_*p*_, at the numerator we get (calling *N* the numerator of Equation (60))

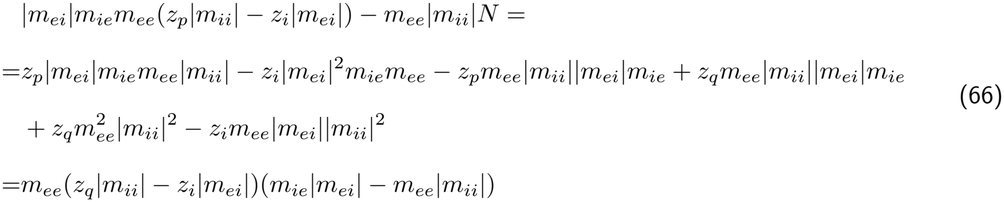

which is positive for every input for which the E-I points can exist. So this threshold is a monotonically increasing function of *z*_*p*_. It is also a monotonically decreasing function of *z*_*q*_, as the numerator of its derivative with respect to it is

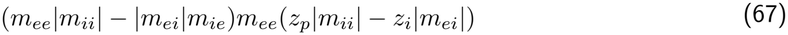

which is negative, as we can immediately see. Clearly, *ε*^{*q,a*}^ is instead a monotonically increasing function of *z*_*q*_ and a monotonically decreasing function of *z*_*p*_. Since when the inputs *z*_*p*_ and *z*_*q*_ are the same, the threshold values *ε*^{*p,a*}^ and *ε*^{*q,a*}^ are also the same and equal to 1, we expect mono-stable regions {*p, a*} and {*q, a*} to appear at *ε* = 1 for asymmetrical inputs, in-between a {*p, q, a*} region, in which neither of these points exists, and a bistable region, in which they both exist. The two mono-stable regions expand monotonically to comprise a larger range of *ε* values, the more different the two drives become. This is indeed what we see in the simulations (Figure 5B). It would also be possible to prove that, assuming without loss of generality that *z*_*p*_ *> z*_*q*_, the mono-stable region {*p, a*} shrinks, for the benefit of the {*p, q, a*} region and of the bistable region, when decreasing either *m*_*ie*_ or |*m*_*ei*_|, until the mono-stable region disappears as soon as they are as small as they can be in a balanced system (so when *m*_*ie*_|*m*_*ei*_| = *m*_*ee*_|*m*_*ii*_|). We omit the proof and just refer to Figure 5Di–ii.

### E-E-I-I Network

We consider a TLN with two excitatory populations *p* and *q* and two inhibitory populations *a* and *b*. We think of *p* as paired with *a* and *q* as paired with *b*, and indicate the connections within one excitatory-inhibitory assembly as *w*_*ee*_, *w*_*ei*_, *w*_*ie*_, and *w*_*ii*_, and the cross-assembly connections as the same variables scaled by a positive parameter. Cross-connections, however, matter in comparison to the full self-side entries of the Jacobian, including the relaxation, so we scale the cross E→E and I→I terms as *εm*_*ee*_ := *ε*(*w*_*ee*_ *−* 1) and *βm*_*ii*_ := *β*(*w*_*ii*_ *−* 1). Since we restrict ourselves to cases in which the excitatory populations are supercritical, we always have *ε >* 0. We note that we also assume that the usual E-I non-suppression and non-explosion conditions (E-I section) hold for each same-side E-I pair. In this way, we obtain the following Jacobian

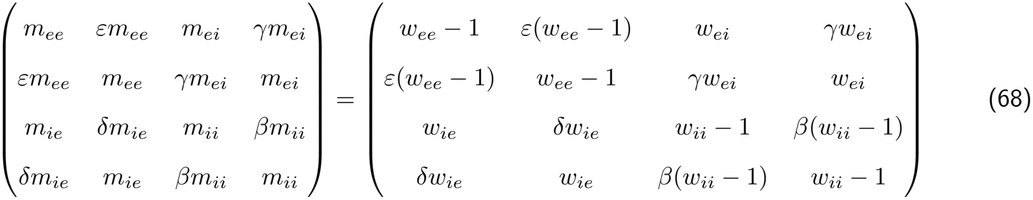

The fixed point {*p, q, a, b*}, in which all the populations are co-active, is stable when

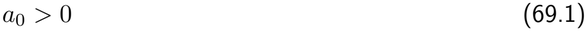

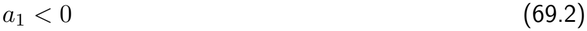

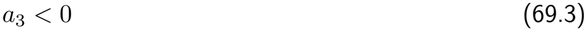

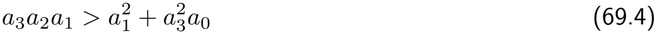

where

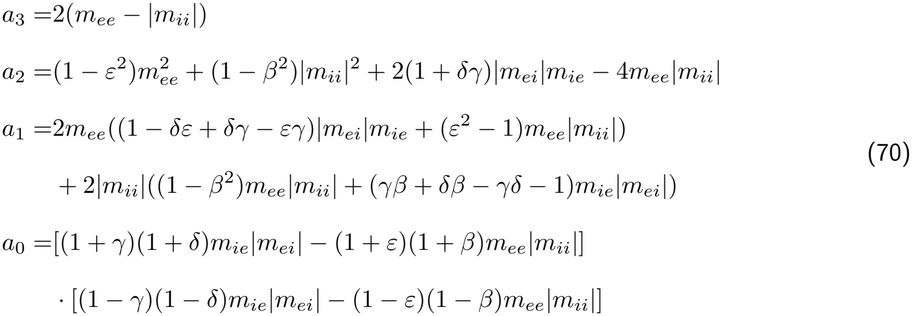

which are, respectively, the trace, the sum of the 2- and 3-dimensional principal minors, and the determinant of the Jacobian of the full system. We can simplify these expressions by defining *W* := (1 + *γ*)(1 + *δ*)*m*_*ie*_|*m*_*ei*_| *−* (1 + *ε*)(1 + *β*)*m*_*ee*_|*m*_*ii*_|, *Z* := (1 *− γ*)(1 *− δ*)*m*_*ie*_|*m*_*ei*_| *−* (1 *− ε*)(1 *− β*)*m*_*ee*_|*m*_*ii*_|, *U* := (1 + *ε*)*m*_*ee*_ *−* (1 + *β*)|*m*_*ii*_|, *V* := (1 *− ε*)*m*_*ee*_ *−* (1 *− β*)|*m*_*ii*_|. In this way, we obtain

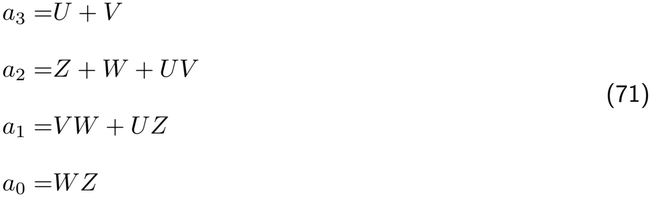

From Equations 69.1–69.4, it follows that *a*_2_ *>* 0. From *a*_0_ *>* 0, it follows that *W* and *Z* must have the same sign. If they are both negative, *a*_2_ *>* 0 implies *UV <* 0. However, since *a*_1_ *<* 0, at least one between *U* and *V* must be positive. Since *a*_3_ *<* 0, the other one must be negative. Hence, *UV <* 0, and we reach a contradiction, as *a*_2_ = *Z* + *W* + *UV* would be negative. Therefore, we conclude that W and Z must both be positive, in order for the co-active state to be stable. The existence conditions of the co-active point are instead

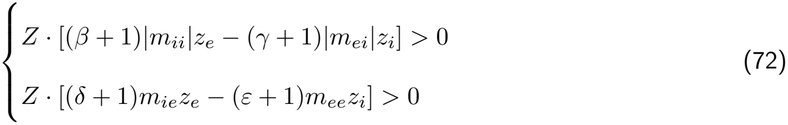

if *a*_0_ (i.e. *D*^*p,q,a,b*^) is positive, and with opposite signs if it is negative. Therefore, the existence of a stable {*p, q, a, b*} also implies the positivity of the second terms in these conditions. These two conditions, together with *W >* 0, act as a two-module equivalent of the existence and stability conditions of a simple E-I network. The condition *Z >* 0, instead, points at specific pairwise relations: existence and stability of the co-active point are favored by “coherent” *γ* and *δ* (both *>* 1 or both *<* 1, such as in balanced inhibition), and not coherent *ε* and *β*.

If the point {*p, q, a, b*} exists and is stable, stimulation of an excitatory population has a positive effect on the other one if

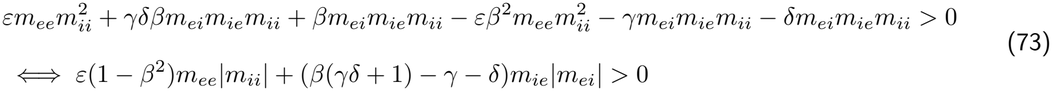

This expression summarizes the effects of all the possible pathways and their paradoxical effects (Figure S4A). Taking for example the perturbation from *p* to *q*, the *p → q* pathway produces the term *εm*_*ee*_, scaled by the determinant of the I-I subsystem, (1*− β*^2^)|*m*_*ii*_|, which determines whether the direct input incoming from *p* has an excitatory or inhibitory effect. The latter occurs when *β >* 1, because in this case the I-I subsystem is in the bistable configuration and excitation-stabilized. The inhibitory *p → a → q* pathway contributes the term *−γ*|*m*_*ei*_|*m*_*ie*_, scaled by |*m*_*ii*_|, representing the determinant of *b* as the only population not already taken into account in this pathway. The inhibitory *p → b → q* pathway is represented by the term *−δ*|*m*_*ei*_|*m*_*ie*_, scaled by |*m*_*ii*_|, the excitatory *p → a → b → q* pathway by +*β*|*m*_*ei*_|*m*_*ie*_|*m*_*ii*_|, and the excitatory *p → b → a → q* pathway by +*γδβ*|*m*_*ei*_|*m*_*ie*_|*m*_*ii*_|.

The cross-connection parameters determine which of these pathways are the most relevant. For example, if *β* = 0, then the last two pathways are irrelevant, and the *p → q* pathway has a non-paradoxical excitatory effect, which competes with the inhibitory effect mediated by both *γ* and *δ*. Specifically, if *β* = *γ* = *δ* = 0 the effect is given by *εm*_*ee*_|*m*_*ii*_|, so it is positive even with a very small *ε*, in accordance with Negrón et al. (2024). With growing *β*, the effect of the *p → a → b → q* and *p → b → a → q* pathways grow: *p* does not only inhibit *q* through the interneurons, but also disinhibit it by inhibiting the competing interneurons. Specifically, for *γ* = *δ* = 0, the perturbation response is

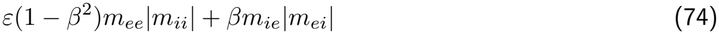

which is always positive for *β ≤* 1. When *β* = 1, the sign of the interaction depends on the product (*δ −* 1)(*γ −* 1): if *δ* and *γ* are coherent, including if they are both *>* 1, the perturbation is positive. For *β >* 1, instead, a growing *ε* has a paradoxical negative effect on the perturbation (provided that the point {*p, q, a, b*} still exists).

The sign of the response of an inhibitory population to the stimulation of the opposite excitatory population is instead given by

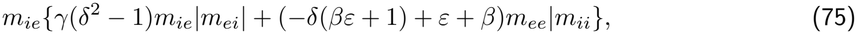

In this case, taking for example the effect of *p* on *b*, we have the *p → b* pathway *δm*_*ie*_, scaled by the *q*–*a* determinant (*γδ*)*m*_*ie*_|*m*_*ei*_| *− m*_*ee*_|*m*_*ii*_|, the inhibition-mediated *p → a → b* pathway +*βm*_*ie*_|*m*_*ii*_|, which however has a positive excitatory effect which scales with *m*_*ee*_, because it acts like an external input in an inhibition-stabilized E-I network, the excitatory *p → q → b* pathway +*εm*_*ie*_|*m*_*ii*_|, scaled by |*m*_*ii*_|, the inhibitory *p → a → q → b* pathway 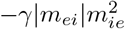, and the inhibitory *p → q → a → b* pathway *−εδβm*_*ee*_*m*_*ie*_|*m*_*ii*_|.

The sign of the effect of an inhibitory population on the opposite excitatory population is given by

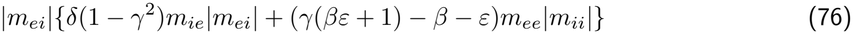

whose meaning is illustrated in Figure S4B. Specifically, when *γ* = 0, *δ* = 1, this becomes

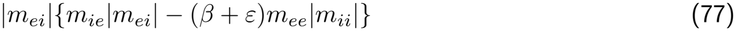

which is positive whenever *β* + *ε <* 1 (provided that the E-I determinant *m*_*ie*_|*m*_*ei*_| *− m*_*ee*_|*m*_*ii*_| is positive).

The interaction between the inhibitory populations is instead

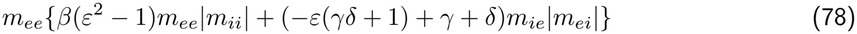

which can also be interpreted in analogous ways.

We now wish to prove that, if the co-active point exists, then the two requirements “*z*_*p*_ decreases the rate of *q*”, and “*z*_*a*_ decreases the rate of *q*” can only be fulfilled at the same time if *γ > β*. The requirements are:

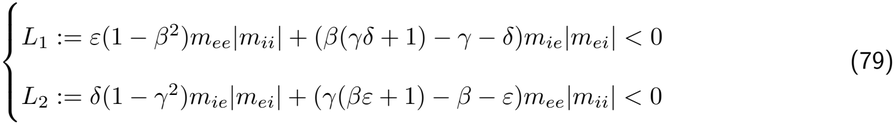

If the requirements are satisfied, it follows that

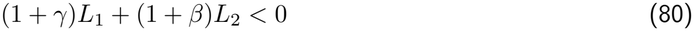

which, with *A* := *m*_*ie*_|*m*_*ei*_| and *B* := *m*_*ee*_|*m*_*ii*_|, factorizes as

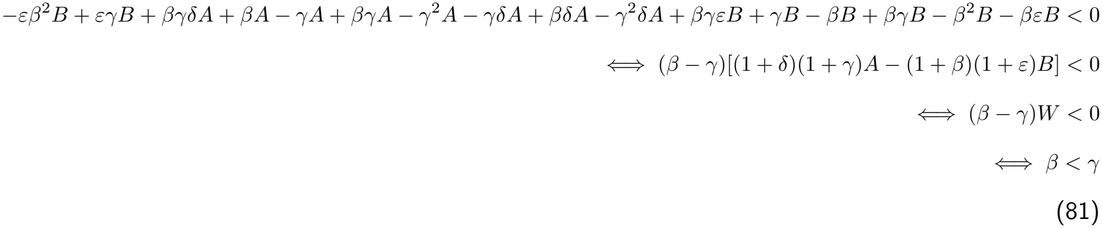

where we have used that *W >* 0.

Since the network is symmetric, conditions for the existence and stability of the E-I-I points {*p, a, b*} and {*q, a, b*} are the same. The existence conditions are

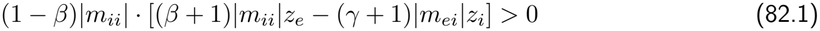

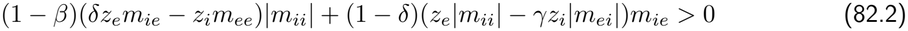

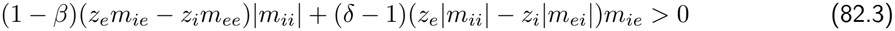

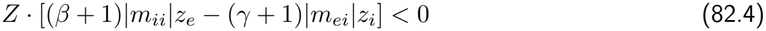

if the determinant *D*^{*p,a,b*}^ is positive (stable E-I-I points), and with reversed signs if it is negative. Condition (82.4) is the opposite of the first one in Equation (72) and it ensures the suppression of the competing E population. This condition can be satisfied in two ways.

If *Z >* 0 and the second factor of Condition (82.4), *N* := (*β* + 1)|*m*_*ii*_|*z*_*e*_ *−* (*γ* + 1)|*m*_*ei*_|*z*_*i*_ is negative, then *β >* 1 (Condition (82.1)). In addition, for this combination the co-active state cannot exist even in its unstable version (Equations (69.1)-(69.4) and (72)), while the unstable state {*a, b*} is the saddle separating the two stable states. This holds because *D*^*a,b*^ has the sign of 1 *− β*, and because Condition (82.1) is the suppression condition for both *p* and *q* in the state {*a, b*}. Therefore, in this case the instability that separates the two E-I-I states originates in the competition between the inhibitory populations.

On the contrary, if *Z <* 0 and *N >* 0, then *β <* 1. In addition, in this case the co-active state always exists and it is the saddle separating the two E-I-I states. This can be seen by adding Condition (82.2) and Condition (82.3), which implies that the term (*δ* + 1)*m*_*ie*_*z*_*e*_ *−* (*ε* + 1)*m*_*ee*_*z*_*i*_, appearing in the second condition of Equation (72), is positive. In addition, the determinant *D*^{*p,a,b*}^, which needs to be positive, can be written as 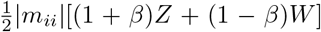, and since *Z <* 0 and *β <* 1, then *W >* 0, which guarantees *D*^{*p,q,a,b*}^ *<* 0. This also shows that the system has a pitchfork bifurcation at *Z* = 0, if the other conditions hold: on the one side there is the stable point {*p, q, a, b*}, while on the other side this point becomes unstable, and the E-I-I points appear. In this case, the instability that separates the two E-I-I states originates in the interactions between excitatory and inhibitory populations, expressed by *Z <* 0. We recall that *Z* := (1 *− γ*)(1 *− δ*)*m*_*ie*_|*m*_*ei*_| *−* (1 *− ε*)(1 *− β*)*m*_*ee*_|*m*_*ii*_| and it is ensured to be negative if *γ* and *δ* are not coherent and *β* and *ε* are coherent (specifically, *<* 1, since *β <* 1). This is an ideal network structure to suppress the competing excitatory population while keeping both I populations active (Figure S4A).

In this setting, we additionally need none of the inhibitory populations to be suppressed: for the “unpaired” population (for example, *b* in the point {*p, a, b*}) this entails an upper threshold on *β* and a lower threshold on *δ* (condition (82.3)), while for the “paired” population (*a* in our example) it also entails an upper threshold on *β* and a more nuanced dependence on *δ*. Specifically, if *γ* is big enough, there is a lower threshold on *δ*, as it can make both terms in condition (82.3) negative, while if *γ* is small, then the effect of *δ* depends on the other parameters. *γ*, in turn, has an upper threshold in case *δ <* 1. There are several configurations of the cross-parameters which can satisfy all the requirements, but one that ensures that this regime emerges is *ε <* 1, *γ < β <* 1, *δ* = 1 (Figure S4Ci).

In particular, if *β* = 0, as considered in Figure 7F–ii, the conditions become

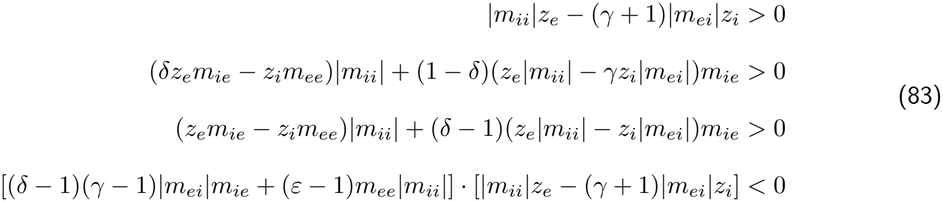

If *z*_*e*_ is large enough and *ε <* 1, the last condition is always satisfied when *γ* and *δ* have different polarities, and for *ε* = 1 (like in Figure 7Fi), the condition becomes exactly (*δ −* 1)(*γ −* 1) *<* 0, which explains why this figure is exactly divided in two coactive and two bistable quadrants by the lines *γ* = 1 and *δ* = 1. As for conditions 2 and 3, they are ensured when *δ* is not too far from 1.

Another possible dynamical regime is the E-E-I bistability between {*p, q, a*} and {*p, q, b*}, whose existence conditions are

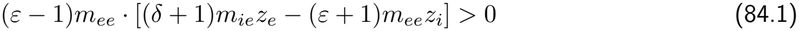

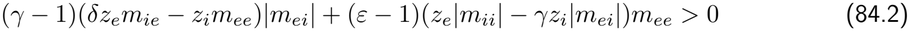

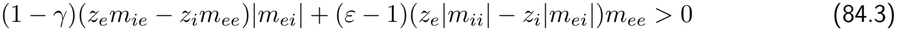

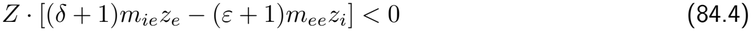

if the determinant *D*^{*p,q,a*}^ is positive (stable E-E-I points), and with reversed signs if it is negative. Also in this case, we have two ways in which condition (84.4) can be satisfied: with *Z >* 0, *ε <* 1, and the unstable saddle {*p, q*}, or with *Z <* 0, *ε >* 1, and the unstable saddle {*p, q, a, b*}. Focusing on the second case, we additionally need the non-suppression of the “unpaired” excitatory population, which requires a lower threshold on *ε* and an upper threshold on *γ*, while non-suppression of the “paired” one also requires a lower threshold on *ε*, an upper threshold on *δ* in case *γ <* 1, and an upper threshold on *γ* in case *δz*_*e*_*m*_*ie*_ *< εz*_*i*_*m*_*ee*_. A combination that ensures that all the conditions are satisfied is *δ > ε >* 1, *β >* 1, *γ* = 1 (Figure S4Cii).

We note that *Z <* 0 (*γ* and *δ* not coherent and/or *β* and *ε* coherent) can contribute to both Conditions (82.4) and (85.4) being satisfied: whether E-I-I bistability, E-E-I bistability, both or none emerges, depends on the other conditions. Namely, if *ε <* 1 and *β <* 1, there can be E-I-I but not E-E-I; if *ε >* 1 and *β >* 1, there can be E-E-I but not E-I-I; if *ε >* 1 and *β <* 1, often the condition will not be satisfied, but there can either be E-I-I, E-E-I, or both. If *ε <* 1 and *β >* 1, instead, none of these points can exist. The fact that these conditions are also associated with the suppression of both one E and one I population, however, makes these conditions ideal for fixed points of the E-I kind.

The conditions for the existence of the “natural” E-I fixed points {*p, a*} and {*q, b*} are given by the opposite of conditions (82.3) and (84.3):

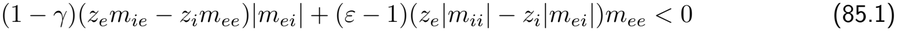

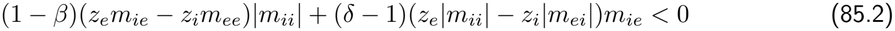

In this case, the conditions *ε <* 1, *γ >* 1, *δ <* 1, and *β >* 1 ensure that the resting state of the network exhibits bistability between the two E-I assemblies (Figure S4Ciii), because of the positivity of the terms that they scale. At least one between *ε <* 1 and *γ >* 1, and one between *δ <* 1 and *β >* 1 is required in order to achieve this regime.

The opposite of conditions (82.2) and (84.2), instead, clarify when the “exchanged” E-I fixed points {*p, b*} and {*q, a*} occur

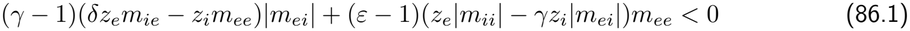

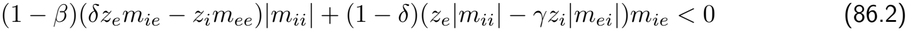

In this case, *δz*_*e*_*m*_*ie*_ *− z*_*i*_*m*_*ee*_ *>* 0 and *z*_*e*_|*m*_*ii*_| *− γz*_*i*_|*m*_*ei*_| *>* 0 are also required for the existence of these points. Provided that these conditions are satisfied, it is clear that *ε <* 1, *γ <* 1, *δ >* 1, and *β >* 1 are the ideal choices for a bistability with this kind of coupling to emerge (Figure S4Civ). Conditions (85) and (86) are not incompatible, and the four E-I combinations can all exist and be stable at the same time, which happens for example when *γ* = *δ* = 1 (Figure S4Cv).

## Supplementary Materials 2: Tables

### Supplementary Materials 3: Supplementary Figures

**Figure S1:**
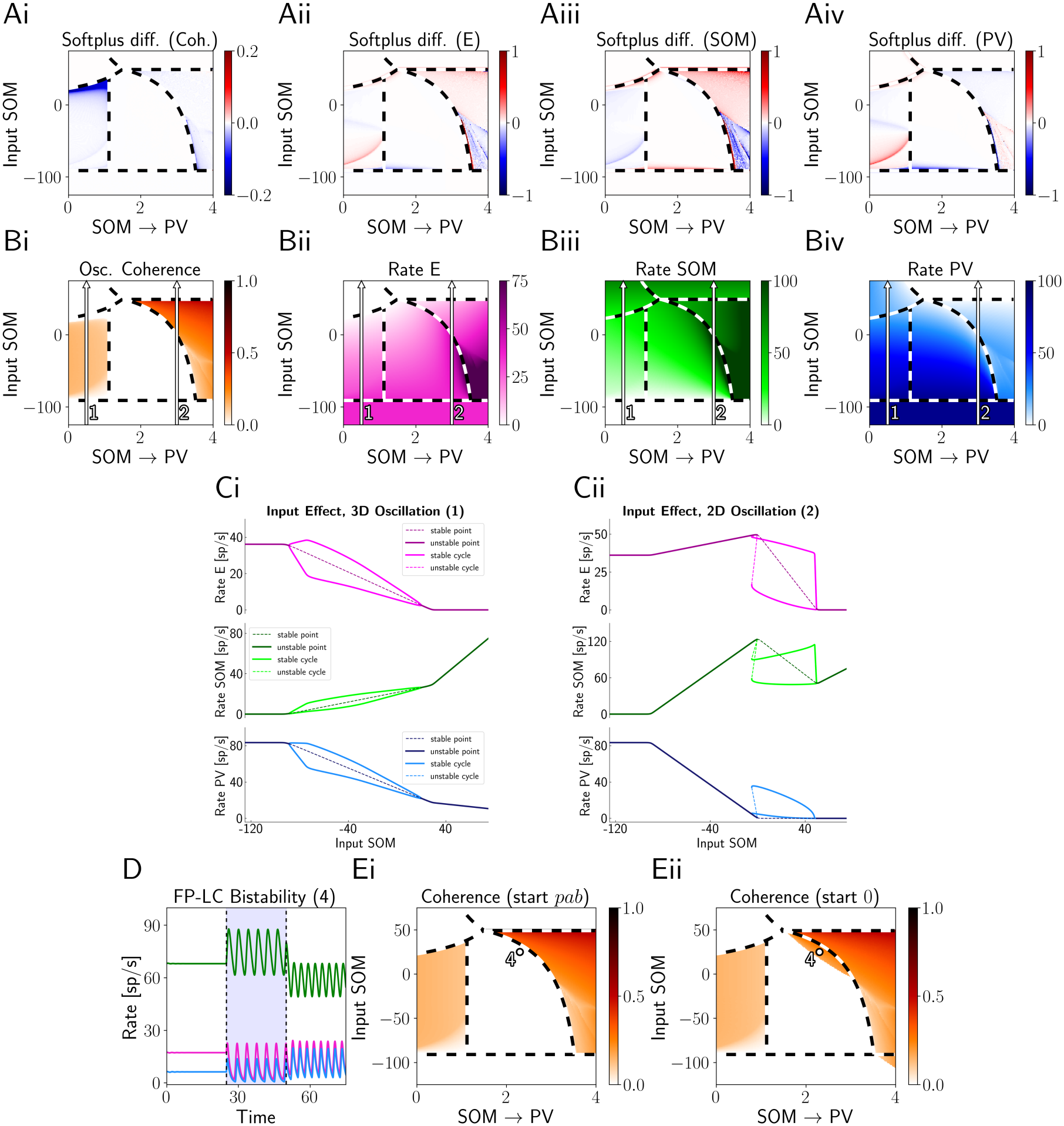
Canonical circuit, supplement to Figure 3. (A) Difference between relevant quantities in the canonical circuit with the softplus activation and in the same network with the threshold-linear activation. (B) Same quantities in the softplus network. Note that the oscillatory region on the left stops before reaching the inactivation lines above and below. (C) Bifurcation diagrams reporting the fixed points and limit cycles encountered when increasing the input to the SOM population, for two different values of the SOM →PV connection, corresponding to arrows 1 and 2 in B. (D) Network activity for dot 4 in E. The shaded blue region represents a transient stimulation of the PV population, highlighting bistability between the fixed point and the limit cycle. (E) Power spectrum coherence over the same parameter space for two different initial conditions, revealing bistability. Parameters in Table 3 of the SM.

**Figure S2:**
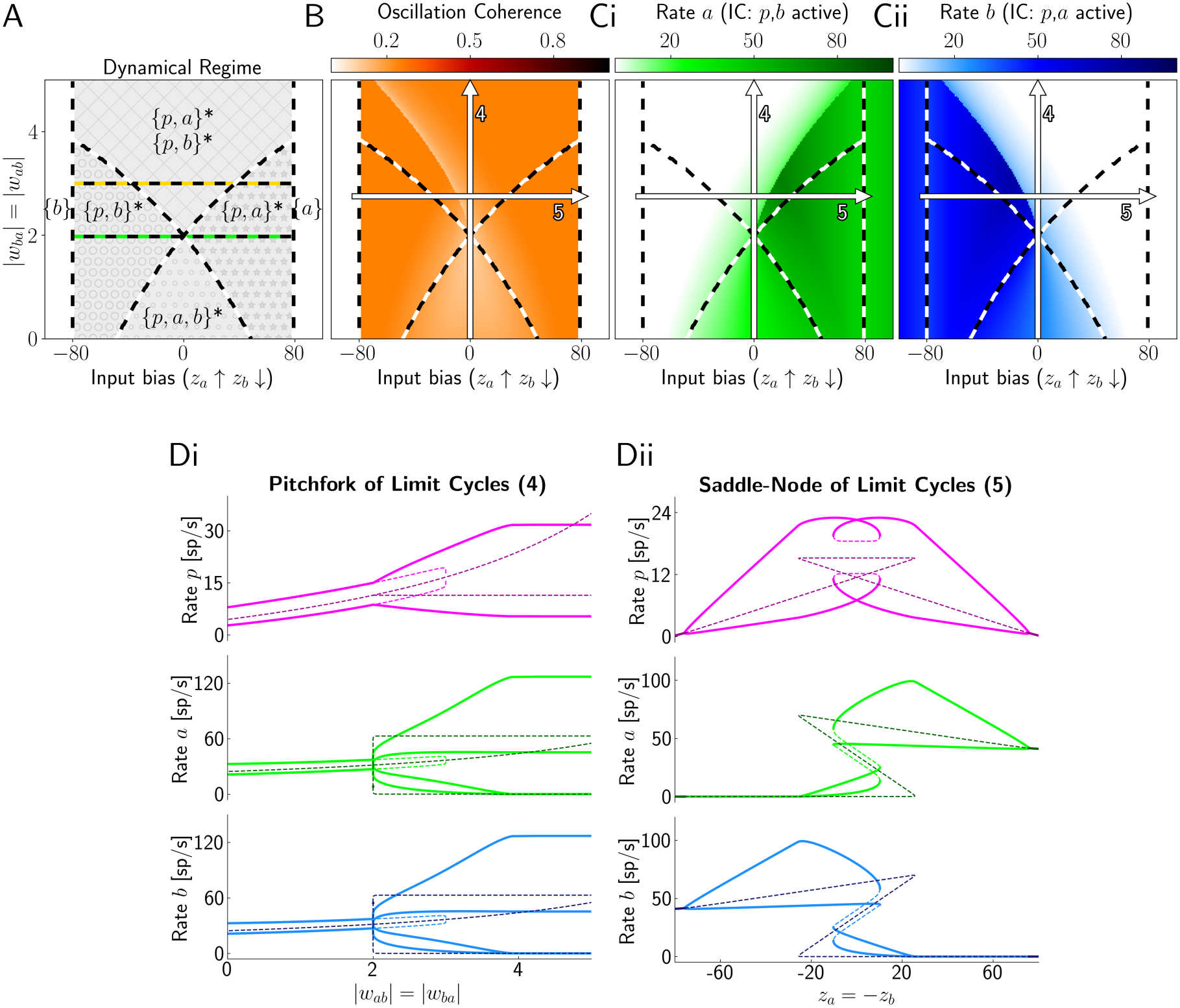
E-I-I network with strong recurrent excitation, supplement to Figure 4. (A) Phase diagram, reporting fixed points and their stability, but not limit cycles. At the black-green line the determinant of {*p, a, b*} changes sign, while at the black-gold line the oscillatory-instability condition for {*p, a, b*} stops holding. Which line is higher determines whether the intermediate non-oscillatory region exists. (B) Power spectrum coherence. (C) Mean steady state firing rates for different initial conditions (IC). (D) Bifurcation diagrams corresponding to arrows 4 and 5 in B–C. Parameters in Table 3 of the SM.

**Figure S3:**
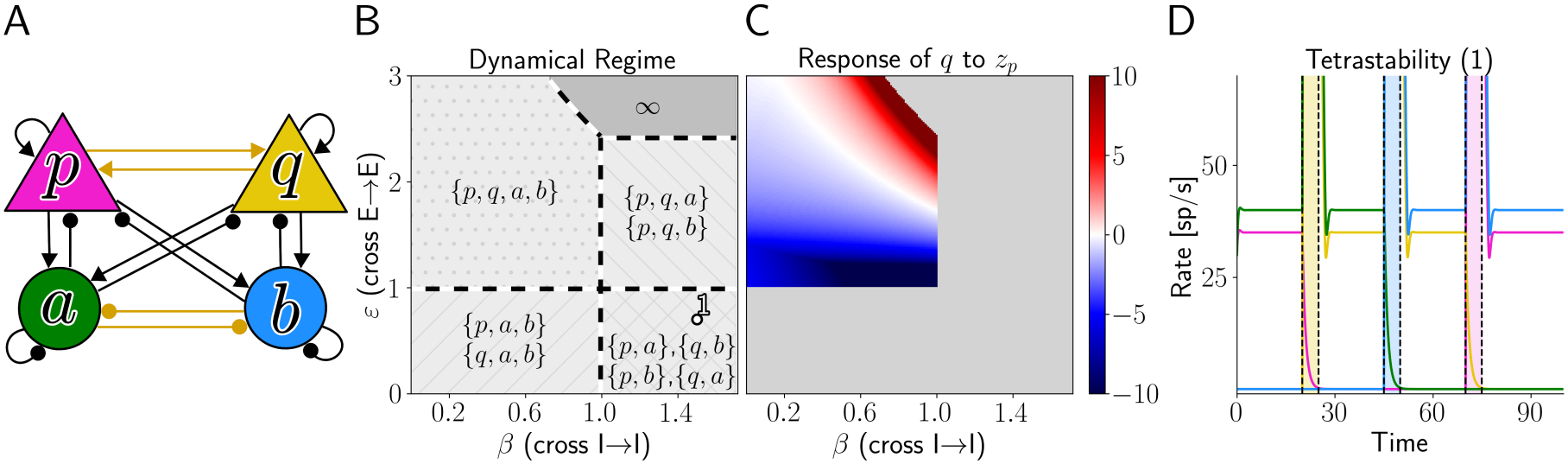
E-E-I-I network, supplement to Figure 7. (A) Network sketch, highlighting in gold the connections that are varied in the simulations. (B) Phase diagram (only stable states are displayed), including tetra-stability in the bottom right corner. (C) Response of *q* to an extra input of 10 delivered to *p*. (D) Network activity for the dot in B. The colored regions represent transient stimulations of the population with the respective color. Parameters in Table 5 of the SM.

**Figure S4:**
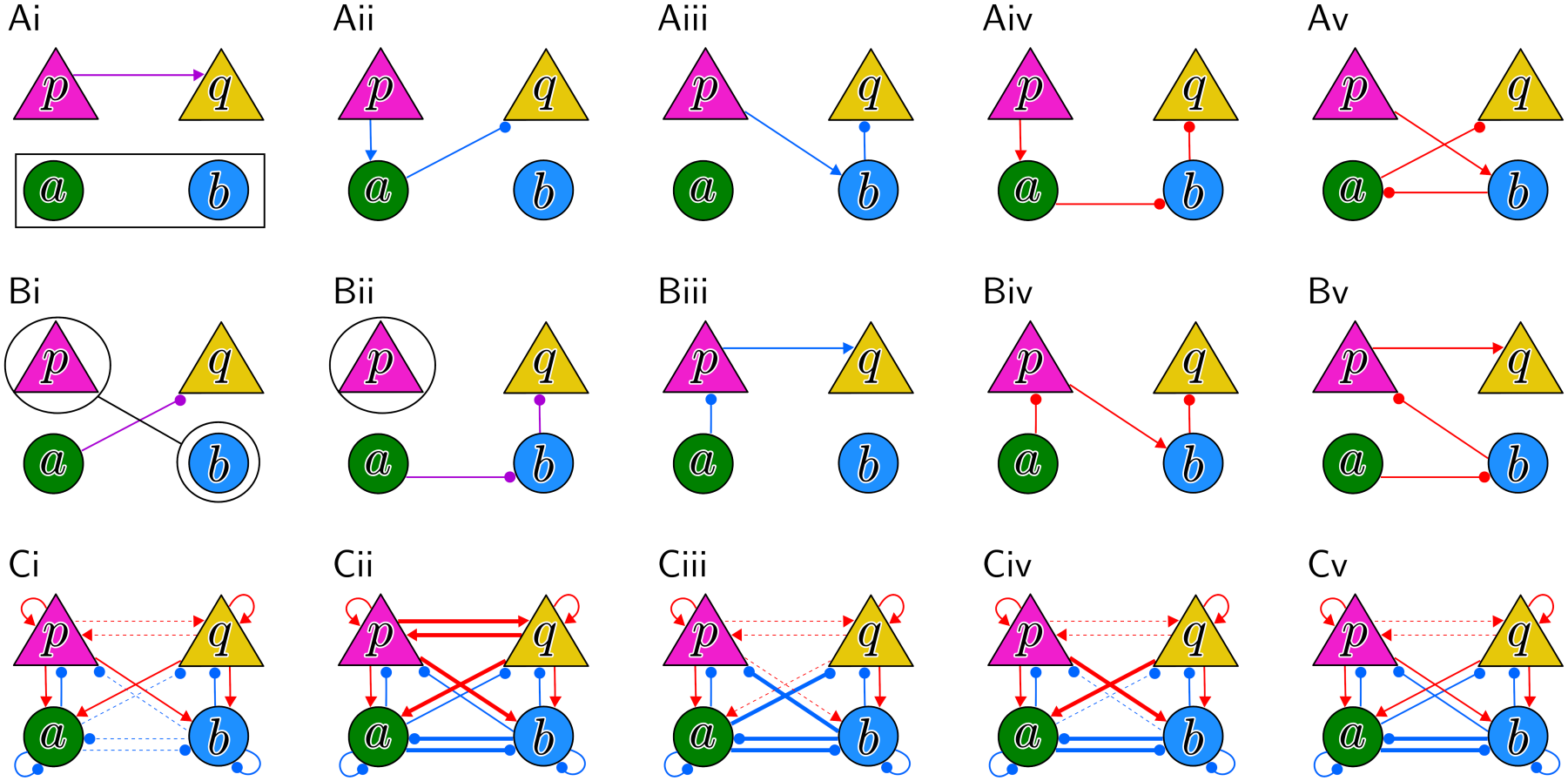
Pathways contributing to perturbation responses and dynamical landscapes of E-E-I-I networks. (A) Pathways that contribute to the *p→ q* perturbation (SM, Equation (73)). The color indicates the excitatory (red) or inhibitory (blue) contribution of the whole pathway. The effect of the purple pathway depends on the determinant of the *a*–*b* subsystem: excitatory if positive, inhibitory if negative. (B) Pathways that contribute to the *a→ q* perturbation (SM, Equation (76)). The effect of the first purple pathway depends on the determinant of the *p*–*b* subsystem: inhibitory if positive, excitatory if negative. The effect of the second pathway depends on the determinant of *p*: excitatory if subcritical, inhibitory if supercritical. (C) Ideal network configurations that support different kinds of bistability. In order: {*p, a, b }* vs {*q, a, b}* (SM, Equation (82)); {*p, q, a }* vs {*p, q, b}* (SM, Equation (84)); {*p, a}* vs {*q, b}* (SM, Equation (85)); {*p, b}* vs {*q, a}* (SM, Equation (86)); {*p, a}* vs {*p, b}* vs {*q, a}* vs {*q, b}* . Thin dashed lines indicate a cross-connection weaker than the self-connection, regular continuous lines indicate a cross-connection equal to the self-connection, and thick lines indicate a cross-connection stronger than the self-connection.

**Figure S5:**
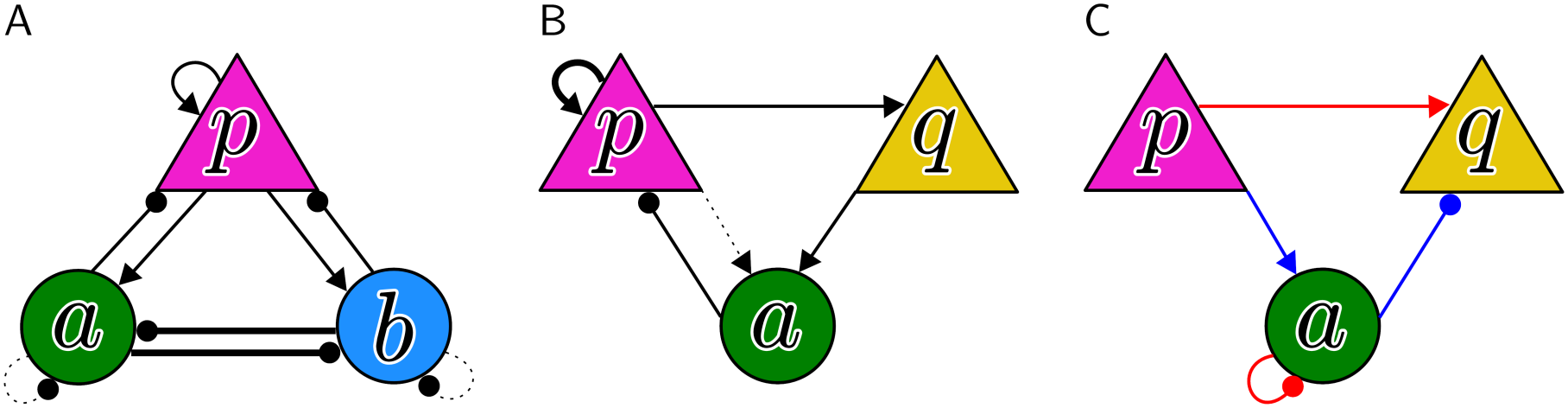
Sketches of microcircuits employed in the Methods to exemplify perturbation responses. (A) Diagram of a {*p, a, b*} network with the *a*–*b* subnetwork in the unstable configuration, where bold lines represent strong connections, dashed lines weak ones. If population *p* stabilizes this network, it presents a paradoxical response to its own input, a case of “excitation-stabilized” regime. (B) Different case of an excitation-stabilized regime. In this case, the *p*–*a* subnetwork is unstable and characterized by divergent activity in both populations. The additional excitatory population *q*, by receiving excitation from *p* and exciting *a*, can stabilize this network. If this is the case, *q* also exhibits a paradoxical response to its own input. (C) Graphical depiction of the pathways involved in the response of *q* to the perturbation of *p*, with the red color representing the overall excitatory pathway and the blue color representing the overall inhibitory pathway. The participation of the *a →a* connection in the excitatory pathway can in this case be understood as a weakening of the competing inhibitory pathway. However, in more complex networks, such scalings involve multiple pathways and are better understood in the framework of Equation (9).

**Figure S6:**
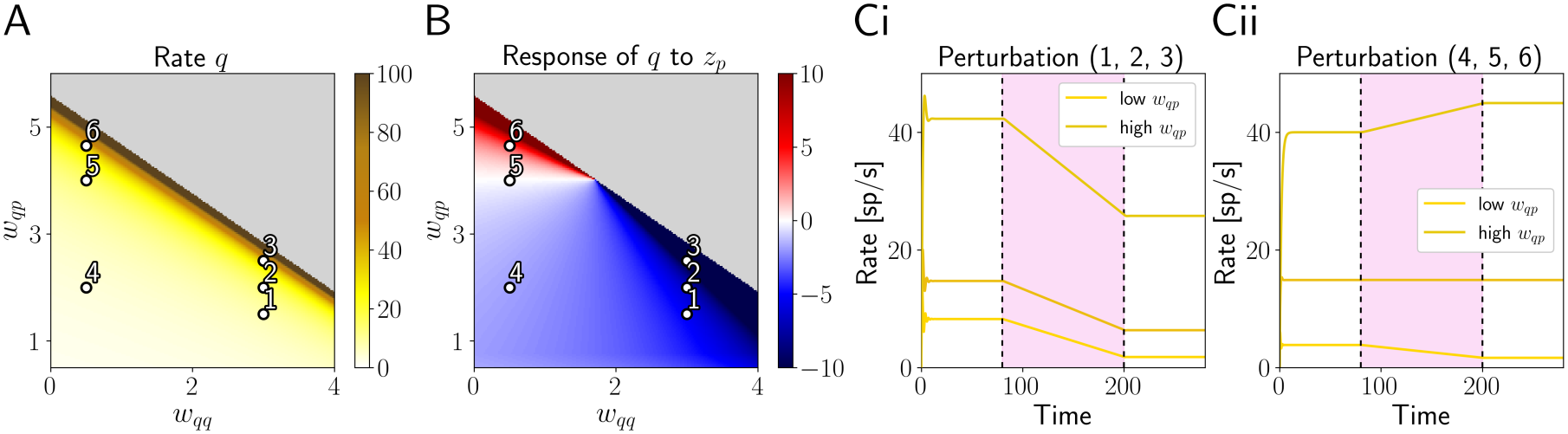
Example of paradoxical modulations of perturbation responses in E-E-I network. (A) Steady state firing rate of *q* in the *p*–*q*–*a* network, as a function of *w*_*qq*_ and *w*_*qp*_. (B) Response of *q* to an extra input of 10 delivered to *p*. (Ci) Evolution of the firing rate over time for the three vertically arranged dots on the right, with lighter gold shades representing lower values of *w*_*qp*_. The red-shaded region indicates the timing of a stimulation of population *p* with an input that gradually increases from 0 to 12 and is then fixed at 12. (Cii) Same, but for the three dots on the left.

**Figure S7:**
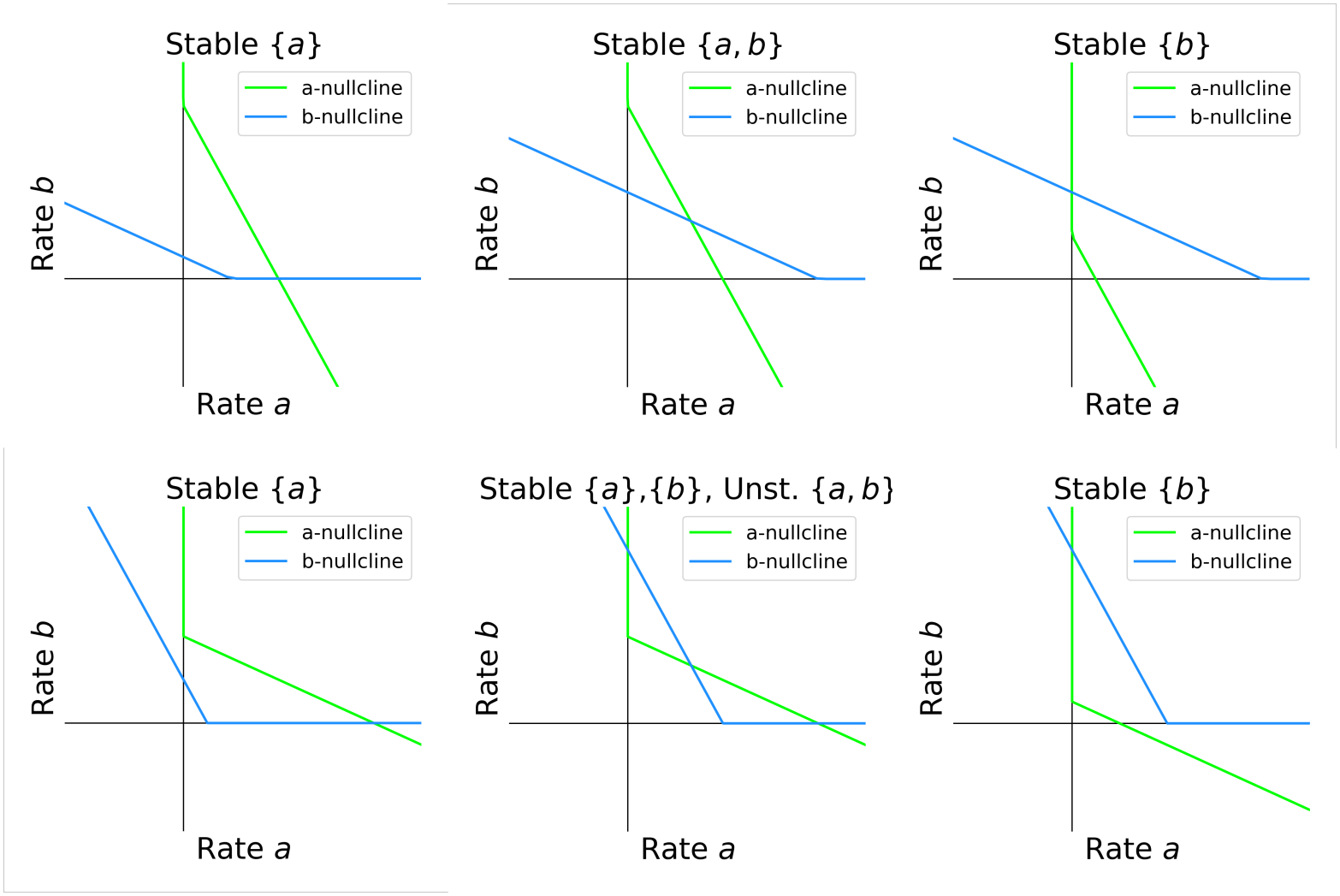
Nullclines of the I-I network. Phase planes depicting the possible nullcline configurations in the *a*–*b* network, with either a positive (top row) or negative (bottom row) determinant *D*^{*a,b*}^. In the latter case, both mono-stable scenarios arise from the bistable one through SN bifurcations.

**Figure S8:**
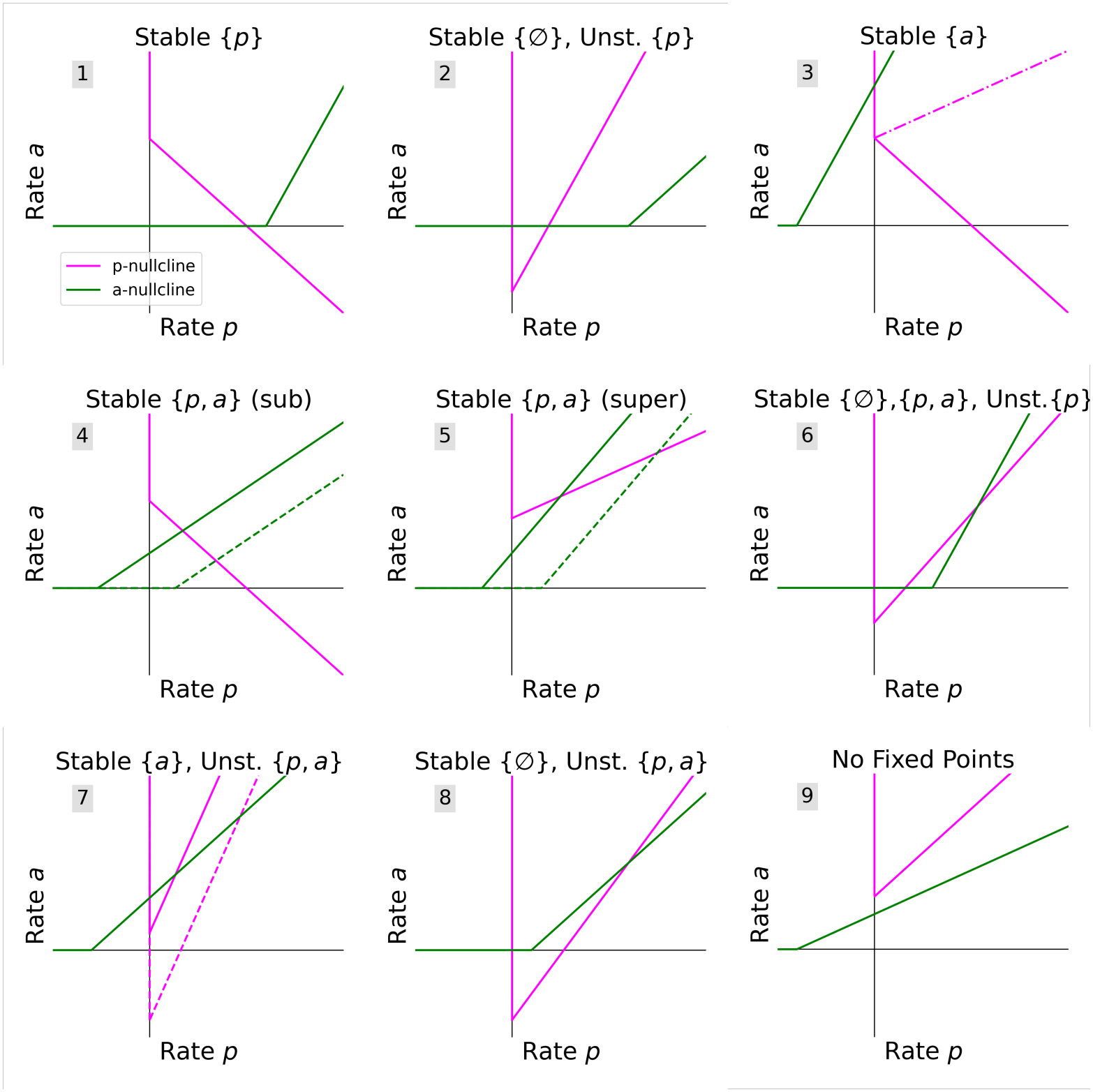
Nullclines of the E-I network. Phase planes depicting the possible nullcline configurations in the *p*–*a* network. Alternative nullclines show when it is possible to have the same fixed point constellation even if one input has the opposite sign (dashed), or *p* is supercritical (dot-dashed) instead of subcritical. When *p* is subcritical, the *p*-nullcline has a negative slope (as a function of *a*); when it is supercritical, a positive one. Panel 6 reports the only bistable configuration. In scenarios 2, 7, 8, and 9 there are divergent trajectories. In scenarios 3 (dashed case), 5, and 6 there can also be divergent trajectories, if the trace and the discriminant are positive. Scenarios 5 and 6 are also compatible with a limit cycle, if the trace is positive and the discriminant is negative.

## References

Adesnik, H. (2017). Synaptic mechanisms of feature coding in the visual cortex of awake mice. Neuron, 95(5):1147–1159.

Beierlein, M., Gibson, J. R., and Connors, B. W. (2003). Two dynamically distinct inhibitory networks in layer 4 of the neocortex. Journal of neurophysiology, 90(5):2987–3000.

Bienenstock, E. (1995). A model of neocortex. Network: Computation in neural systems, 6(2):179–224.

Brunel, N. (2000). Dynamics of sparsely connected networks of excitatory and inhibitory spiking neurons. Journal of computational neuroscience, 8:183–208.

Buzsáki, G. and Draguhn, A. (2004). Neuronal oscillations in cortical networks. Science, 304(5679):1926–1929.

Campagnola, L., Seeman, S. C., Chartrand, T., Kim, L., Hoggarth, A., Gamlin, C., Ito, S., Trinh, J., Davoudian, P., Radaelli, C., et al. (2022). Local connectivity and synaptic dynamics in mouse and human neocortex. Science, 375(6585):eabj5861.

Canto-Bustos, M., Friason, F. K., Bassi, C., and Oswald, A.-M. M. (2022). Disinhibitory circuitry gates associative synaptic plasticity in olfactory cortex. Journal of Neuroscience, 42(14):2942–2950.

Colgin, L. L., Denninger, T., Fyhn, M., Hafting, T., Bonnevie, T., Jensen, O., Moser, M.-B., and Moser,E. I. (2009). Frequency of gamma oscillations routes flow of information in the hippocampus. Nature, 462(7271):353–357.

Curto, C., Geneson, J., and Morrison, K. (2019). Fixed points of competitive threshold-linear networks.Neural computation, 31(1):94–155.

Curto, C., Geneson, J., and Morrison, K. (2024). Stable fixed points of combinatorial threshold-linear networks. Advances in applied mathematics, 154:102652.

Curto, C., Langdon, C., and Morrison, K. (2020). Combinatorial geometry of threshold-linear networks.arXiv preprint arXiv:2008.01032.

Curto, C. and Morrison, K. (2016). Pattern completion in symmetric threshold-linear networks. Neural computation, 28(12):2825–2852.

De Kock, C. P., Pie, J., Pieneman, A. W., Mease, R. A., Bast, A., Guest, J. M., Oberlaender, M., Mansvelder, H. D., and Sakmann, B. (2021). High-frequency burst spiking in layer 5 thick-tufted pyramids of rat primary somatosensory cortex encodes exploratory touch. Communications biology, 4(1):709.

Deuchars, J. and Thomson, A. (1996). CA1 pyramid-pyramid connections in rat hippocampus in vitro: dual intracellular recordings with biocytin filling. Neuroscience, 74(4):1009–1018.

Eckhorn, R., Bauer, R., Jordan, W., Brosch, M., Kruse, W., Munk, M., and Reitboeck, H. (1988). Coherent oscillations: a mechanism of feature linking in the visual cortex? Multiple electrode and correlation analyses in the cat. Biological cybernetics, 60:121–130.

Ermentrout, B. and Mahajan, A. (2003). Simulating, analyzing, and animating dynamical systems: a guide to XPPAUT for researchers and students. Appl. Mech. Rev., 56(4):B53–B53.

Evangelista, R., Cano, G., Cooper, C., Schmitz, D., Maier, N., and Kempter, R. (2020). Generation of sharp wave-ripple events by disinhibition. Journal of Neuroscience, 40(41):7811–7836.

Fries, P. (2009). Neuronal gamma-band synchronization as a fundamental process in cortical computation.Annual review of neuroscience, 32(1):209–224.

Hunt, D. L., Linaro, D., Si, B., Romani, S., and Spruston, N. (2018). A novel pyramidal cell type promotes sharp-wave synchronization in the hippocampus. Nature neuroscience, 21(7):985–995.

Jercog, D., Roxin, A., Bartho, P., Luczak, A., Compte, A., and De La Rocha, J. (2017). Up-down cortical dynamics reflect state transitions in a bistable network. eLife, 6:e22425.

Karnani, M. M., Jackson, J., Ayzenshtat, I., Sichani, A. H., Manoocheri, K., Kim, S., and Yuste, R. (2016). Opening holes in the blanket of inhibition: localized lateral disinhibition by VIP interneurons. Journal of neuroscience, 36(12):3471–3480.

Kato, H. K., Asinof, S. K., and Isaacson, J. S. (2017). Network-level control of frequency tuning in auditory cortex. Neuron, 95(2):412–423.

Keeley, S., Fenton, A. A., and Rinzel, J. (2017). Modeling fast and slow gamma oscillations with interneurons of different subtype. Journal of neurophysiology, 117(3):950–965.

Kenet, T., Bibitchkov, D., Tsodyks, M., Grinvald, A., and Arieli, A. (2003). Spontaneously emerging cortical representations of visual attributes. Nature, 425(6961):954–956.

Kim, M.-H., Znamenskiy, P., Iacaruso, M. F., and Mrsic-Flogel, T. D. (2018). Segregated subnetworks of intracortical projection neurons in primary visual cortex. Neuron, 100(6):1313–1321.

Kuan, A. T., Bondanelli, G., Driscoll, L. N., Han, J., Kim, M., Hildebrand, D. G., Graham, B. J., Wilson,D. E., Thomas, L. A., Panzeri, S., et al. (2024). Synaptic wiring motifs in posterior parietal cortex support decision-making. Nature, 627(8003):367–373.

LaFosse, P. K., Zhou, Z., O’Rawe, J. F., Friedman, N. G., Scott, V. M., Deng, Y., and Histed, M. H. (2024). Cellular-resolution optogenetics reveals attenuation-by-suppression in visual cortical neurons. Proceedings of the National Academy of Sciences, 121(45):e2318837121.

Lagzi, F., Bustos, M. C., Oswald, A.-M., and Doiron, B. (2021). Assembly formation is stabilized by parvalbumin neurons and accelerated by somatostatin neurons. BioRxiv, pages 2021–09.

Lagzi, F. and Fairhall, A. L. (2024). Emergence of co-tuning in inhibitory neurons as a network phenomenon mediated by randomness, correlations, and homeostatic plasticity. Science Advances, 10(12):eadi4350.

Laing, C. R. and Chow, C. C. (2002). A spiking neuron model for binocular rivalry. Journal of computational neuroscience, 12:39–53.

Lein, E., Borm, L. E., and Linnarsson, S. (2017). The promise of spatial transcriptomics for neuroscience in the era of molecular cell typing. Science, 358(6359):64–69.

Letzkus, J. J., Wolff, S. B., and Lüthi, A. (2015). Disinhibition, a circuit mechanism for associative learning and memory. Neuron, 88(2):264–276.

Litwin-Kumar, A., Rosenbaum, R., and Doiron, B. (2016). Inhibitory stabilization and visual coding in cortical circuits with multiple interneuron subtypes. Journal of neurophysiology, 115(3):1399–1409.

Liu, W.-M. (1994). Criterion of Hopf bifurcations without using eigenvalues. Journal of Mathematical Analysis and Applications, 182(1):250–256.

Lu, Y. and Rinzel, J. (2024). Firing rate models for gamma oscillations in I-I and E-I networks. Journal of Computational Neuroscience, 52(4):247–266.

Madisen, L., Zwingman, T. A., Sunkin, S. M., Oh, S. W., Zariwala, H. A., Gu, H., Ng, L. L., Palmiter,R. D., Hawrylycz, M. J., Jones, A. R., et al. (2010). A robust and high-throughput Cre reporting and characterization system for the whole mouse brain. Nature neuroscience, 13(1):133–140.

Miller, K. D. and Palmigiano, A. (2020). Generalized paradoxical effects in excitatory/inhibitory networks.BioRxiv, pages 2020–10.

Moreno-Bote, R., Rinzel, J., and Rubin, N. (2007). Noise-induced alternations in an attractor network model of perceptual bistability. Journal of neurophysiology, 98(3):1125–1139.

Morrison, K., Degeratu, A., Itskov, V., and Curto, C. (2024). Diversity of emergent dynamics in competitive threshold-linear networks. SIAM Journal on Applied Dynamical Systems, 23(1):855–884.

Negrón, A., Getz, M. P., Handy, G., and Doiron, B. (2024). The mechanics of correlated variability in segregated cortical excitatory subnetworks. Proceedings of the National Academy of Sciences, 121(28):e2306800121.

Ogando, M. B., Abdeladim, L., Sit, K. K., Shin, H., Sridharan, S., Gopakumar, K., and Adesnik, H. (2025).Feature-specific inhibitory connectivity augments the accuracy of cortical representations. bioRxiv.

Okun, M. and Lampl, I. (2008). Instantaneous correlation of excitation and inhibition during ongoing and sensory-evoked activities. Nature neuroscience, 11(5):535–537.

O’Hashi, K., Fekete, T., Deneux, T., Hildesheim, R., Van Leeuwen, C., and Grinvald, A. (2018). Interhemispheric synchrony of spontaneous cortical states at the cortical column level. Cerebral Cortex, 28(5):1794–1807.

Palmigiano, A., Fumarola, F., Mossing, D. P., Kraynyukova, N., Adesnik, H., and Miller, K. D. (2020). Common rules underlying optogenetic and behavioral modulation of responses in multi-cell-type V1 circuits. bioRxiv, pages 2020–11.

Parker, M. M., Rubin, J. E., and Huang, C. (2025). State modulation in spatial networks with three interneuron subtypes. Science Advances, 11(26):eads9134.

Pfeffer, C. K., Xue, M., He, M., Huang, Z. J., and Scanziani, M. (2013). Inhibition of inhibition in visual cortex: the logic of connections between molecularly distinct interneurons. Nature neuroscience, 16(8):1068–1076.

Richter, L. M. and Gjorgjieva, J. (2022). A circuit mechanism for independent modulation of excitatory and inhibitory firing rates after sensory deprivation. Proceedings of the National Academy of Sciences, 119(32):e2116895119.

Routh, E. J. (1877). A treatise on the stability of a given state of motion, particularly steady motion: being the essay to which the Adams prize was adjudged in 1877, in the University of Cambridge. Macmillan and Company.

Sadeh, S. and Clopath, C. (2020). Theory of neuronal perturbome in cortical networks. Proceedings of the National Academy of Sciences, 117(43):26966–26976.

Sadeh, S. and Clopath, C. (2021). Inhibitory stabilization and cortical computation. Nature Reviews Neuroscience, 22(1):21–37.

Sammons, R. P., Masserini, S., Moreno Velasquez, L., Metodieva, V. D., Cano, G., Sannio, A., Orlando, M., Maier, N., Kempter, R., and Schmitz, D. (2025). Sub-type specific connectivity between CA3 pyramidal neurons may underlie their sequential activation during sharp waves. eLife, 13:RP98653.

Sanzeni, A., Akitake, B., Goldbach, H. C., Leedy, C. E., Brunel, N., and Histed, M. H. (2020). Inhibition stabilization is a widespread property of cortical networks. eLife, 9:e54875.

Sasaki, T., Matsuki, N., and Ikegaya, Y. (2007). Metastability of active CA3 networks. Journal of Neuroscience, 27(3):517–528.

Shpiro, A., Curtu, R., Rinzel, J., and Rubin, N. (2007). Dynamical characteristics common to neuronal competition models. Journal of neurophysiology, 97(1):462–473.

Shpiro, A., Moreno-Bote, R., Rubin, N., and Rinzel, J. (2009). Balance between noise and adaptation in competition models of perceptual bistability. Journal of computational neuroscience, 27:37–54.

Tahvili, F., Vinck, M., and di Volo, M. (2025). PV and SOM cells play distinct causal roles in controlling network oscillations and stability. Cell Reports, 44(8).

Tahvili, F., Vinck, M., and Di Volo, M. (2026). A mean-field model of neural networks with PV and SOM interneurons reveals connectivity-based mechanisms of gamma oscillations. PLOS Computational Biology, 22(6):e1014378.

Tasic, B., Yao, Z., Graybuck, L. T., Smith, K. A., Nguyen, T. N., Bertagnolli, D., Goldy, J., Garren, E., Economo, M. N., Viswanathan, S., et al. (2018). Shared and distinct transcriptomic cell types across neocortical areas. Nature, 563(7729):72–78.

Tsodyks, M. V., Skaggs, W. E., Sejnowski, T. J., and McNaughton, B. L. (1997). Paradoxical effects of external modulation of inhibitory interneurons. Journal of neuroscience, 17(11):4382–4388.

Van Vreeswijk, C., Abbott, L., and Bard Ermentrout, G. (1994). When inhibition not excitation synchronizes neural firing. Journal of computational neuroscience, 1:313–321.

Veit, J., Hakim, R., Jadi, M. P., Sejnowski, T. J., and Adesnik, H. (2017). Cortical gamma band synchronization through somatostatin interneurons. Nature neuroscience, 20(7):951–959.

Veit, J., Handy, G., Mossing, D. P., Doiron, B., and Adesnik, H. (2023). Cortical VIP neurons locally control the gain but globally control the coherence of gamma band rhythms. Neuron, 111(3):405–417.

Vogels, T. P., Sprekeler, H., Zenke, F., Clopath, C., and Gerstner, W. (2011). Inhibitory plasticity balances excitation and inhibition in sensory pathways and memory networks. Science, 334(6062):1569–1573.

Waitzmann, F., Wu, Y. K., and Gjorgjieva, J. (2024). Top–down modulation in canonical cortical circuits with short-term plasticity. Proceedings of the National Academy of Sciences, 121(16):e2311040121.

Wang, X.-J. and Buzsáki, G. (1996). Gamma oscillation by synaptic inhibition in a hippocampal interneuronal network model. Journal of neuroscience, 16(20):6402–6413.

Wehr, M. and Zador, A. M. (2003). Balanced inhibition underlies tuning and sharpens spike timing in auditory cortex. Nature, 426(6965):442–446.

Whittington, M. A., Traub, R. D., Kopell, N., Ermentrout, B., and Buhl, E. H. (2000). Inhibition-based rhythms: experimental and mathematical observations on network dynamics. International journal of psychophysiology, 38(3):315–336.

Wilson, H. R. and Cowan, J. D. (1972). Excitatory and inhibitory interactions in localized populations of model neurons. Biophysical journal, 12(1):1–24.

Wu, Y. K. and Gjorgjieva, J. (2023). Inhibition stabilization and paradoxical effects in recurrent neural networks with short-term plasticity. Physical Review Research, 5(3):033023.

Znamenskiy, P., Kim, M.-H., Muir, D. R., Iacaruso, M. F., Hofer, S. B., and Mrsic-Flogel, T. D. (2024).Functional specificity of recurrent inhibition in visual cortex. Neuron, 112(6):991–1000.

